# Quantitative whole-tissue 3D imaging reveals bacteria in close association with mouse jejunum mucosa

**DOI:** 10.1101/2022.06.17.496478

**Authors:** Roberta Poceviciute, Said R. Bogatyrev, Anna E. Romano, Amanda H. Dilmore, Octavio Mondragón-Palomino, Heli Takko, Rustem F. Ismagilov

## Abstract

**Background:** The small intestine (SI) is the primary site of nutrient absorption, so its large surface area lacks the thick protective mucus that is characteristic of the large intestine. Because the SI epithelium is relatively exposed, any microbes that colonize the thin mucosa of the SI may exert a substantial effect on the host. Thus far, potential bacterial colonization of the SI mucosa has only been documented in disease states, suggesting mucosal colonization is a rare occurrence, likely requiring multiple perturbations.

**Results:** Here, we tested whether we could induce bacterial association with jejunum mucosa by a combination of malnutrition and oral co-gavage with a specific bacterial cocktail (*E. coli* and *Bacteroides* spp.) that has previously induced environmental enteropathy in mouse models. To overcome the current limitations in imaging and allow definite determination of whether bacterial colonization of the SI mucosa is occurring, we optimized our previously developed whole-tissue three-dimensional (3D) imaging tools with third-generation hybridization chain reaction (HCR v3.0) probes. Only in mice that were malnourished and gavaged with the bacterial cocktail did we detect dense bacterial clusters surrounding intestinal villi suggestive of colonization. Healthy mice gavaged with bacteria and malnourished mice not gavaged with bacteria showed no evidence of mucosal colonization. Furthermore, in malnourished mice gavaged with bacteria we detected villus loss, which may represent one possible consequence that bacterial colonization of the SI mucosa has on the host.

**Conclusions:** Our results suggest that dense bacterial colonization of jejunum mucosa is possible in the presence of multiple perturbations and that villus loss may be one possible consequence to such colonization. Furthermore, our results demonstrate the utility of whole-tissue 3D imaging tools. Although 2D imaging of thin sections may have failed to detect and capture the full spatial complexity of such rare events, whole-tissue 3D imaging tools enabled their detection over large areas of intestinal mucosa and visualization of their spatial complexity in 3D.

## Background

Unlike the large intestine (LI), which is dispensable [1], the small intestine (SI) cannot be fully removed [2] because it performs the essential function of food digestion and nutrient absorption. However, compared with LI microbiota, SI microbiota remains underrepresented in microbiota research, in part because it is less accessible: although non-invasive sampling of feces can be leveraged to study LI microbiota, invasive endoscopy procedures are required to access SI microbiota in humans [3]. The SI is also longer than the LI [4, 5]; as a result, although endoscopy can be used to access the upper SI (duodenum) and colonoscopy can be used to sample the lower SI (ileum), even more invasive or expensive technologies are required to access the middle SI (jejunum) [3, 6]. Furthermore, the underrepresentation of the SI in microbiome research may be explained by its orders of magnitude lower microbial loads compared with the LI (10^2^-10^7^ colony forming units per gram (CFU/g) in the SI and 10^10^-10^13^ CFU/g in the LI [6–8]). At these lower microbial loads of the SI, any low-abundance taxa are also more challenging to detect and quantify [9, 10]. Even less is known about microbial colonization of the mucosa of the SI; however, one can extrapolate that any microbes that do colonize the mucosa of the SI may have a disproportionate impact on the host. The SI has a large surface area (15 times larger than the LI [4]), and the SI epithelium is exposed with only a loose discontinuous mucus, as opposed to the additional dense protective mucus layer characteristic of the LI [5, 11, 12]). Thus, bacterial colonization of SI mucosa would result in a large, direct host-microbe contact area where the host and the microbe may intimately interact.

To protect the SI mucosa from bacterial colonizers and pathogens, the host is armed with a wide range of mucosal defenses, including abundant intraepithelial lymphocytes and other immune cells [5], antimicrobials produced by Paneth cells [13], secretory immunoglobulin A (IgA) [14], mucus (which does not form a dense mucus layer but protects mucosa by other mechanisms [5]), and oxygen (which should suppress the growth of strict anaerobes [15]). Given these defenses, one may anticipate that bacterial colonization of SI mucosa would be rare across space and/or over time in healthy organisms and would perhaps be more prevalent when mucosal defenses are weakened by genetic or environmental perturbations. For example, bacteria were detected by 2D imaging of thin sections in SI mucosa of a genetic cystic fibrosis mouse model but not wild-type controls [16]; in cystic fibrosis, the altered physiology of mucus [16] and Paneth cells [17] is thought to permit bacterial invasion of the mucosa. Environmental factors, such as age, diet, xenobiotics, and chemical or biological contamination, may also increase the likelihood of bacterial colonization of SI mucosa. For example, bacteria have been detected in the ileal mucosa of old mice [18] or mice fed a high-fat diet [19], as well as the jejunum mucosa of mice fed malnourishing diets [20]. Finally, synergistic bacterial interactions [21] may enable bacteria to overcome mucosal defenses and colonize the mucosa. For example, to recapitulate features of environmental enteropathy (EE), a sub-clinical disorder of the SI characterized by inflammation and damage to the SI epithelium and implicated in childhood malnutrition [22–24], both *Escherichia coli* and *Bacteroides/Parabacteroides* spp. had to be gavaged to malnourished mice [20]. In these malnourished co-gavaged mice, an increase in tissue-adherent *Enterobacteriaceae* was also reported [20], which led us to hypothesize that the EE phenotype results at least partially from the colonization of SI mucosa by the gavaged bacteria. However, it remains unknown whether the bacteria detected in the jejunum mucosa of those EE mice [20] represented true mucosal colonizers or lumenal bacteria present in the mucosa as an artifact of mixing of lumenal contents [25]. Furthermore, because the studies examined only a few representative thin sections, the extent, heterogeneity, and 3D spatial structure of the host-microbe mucosal interface remains poorly understood.

Whole-tissue 3D imaging tools may be best suited to detect and characterize rare events (e.g. mucosal colonization) in the large surface area of the SI. A breakthrough in 3D imaging took place in the field of neuroscience in which various hydrogel tissue stabilization, lipid clearing, and refractive index matching modalities, such as CLARITY, CUBIC, and others, were developed to reduce light scattering and increase imaging depth [26]. Steps have been taken to translate CLARITY to 3D of imaging host-microbe interface in the gut. For example, in combination with sensitive detection of bacteria by hybridization chain reaction (HCR v2.0) [27, 28], CLARITY has been used to visualize bacteria in the sputum of cystic fibrosis patients [29]. However, that application of CLARITY imaged bacteria in mucosal excretions, but did not image directly in the mucosa, and HCR v2.0 suffered from background amplification at depth [30]. CLARITY has been also applied to image the intact mouse gut [31, 32]; however, neither bacteria nor mucus were visualized by these methods. CUBIC has been used to visualize host-microbe interface in 3D along the entire gut of mice, however, this method did not allow to image native bacteria with taxonomic resolution [33]. Finally, we have recently developed CLARITY-based imaging tools that allow one to scan large areas of intestinal mucosa and profile heterogeneous bacterial association with intestinal mucosa in 3D with highly sensitive and specific taxonomic resolution of bacteria [34]. Thus, we reasoned that our tools are suited to study rare events of bacterial colonization of SI mucosa.

As a first step toward mechanistic understanding of host-microbe interactions on the SI mucosa, we set out to answer whether our 3D imaging tools can be used to detect bacterial association with the SI mucosa indicative of colonization. We focused on the jejunum to reduce the risk that duodenum microbiota could be contaminated with oral or stomach bacteria due to gastric emptying [35, 36], and that ileum microbiota may contain colonic bacteria due to retrograde transport [37]. To study bacterial association with jejunum mucosa and look for evidence of mucosal colonization, we leveraged three strategies. First, we used the previously reported EE mouse model [20]. We expected that the gavaged *E. coli* and *Bacteroides/Parabacteroides* spp. cocktail would not only recapitulate EE features in malnourished mice [20] but that these microbes would colonize the jejunum mucosa. Second, we applied our whole-tissue 3D imaging tools and HCR v3.0 technology [30] to image any mucosa-associated microbes with high sensitivity and specificity over large areas of the mucosa and in 3D. Finally, we performed a series of experiments to evaluate whether bacterial colonization of jejunum mucosa was plausible in malnourished co-gavaged mice. With these approaches, we documented the association of bacterial taxa with jejunum mucosa in mice and observed one possible host response to this association.

## Results

### EE phenotype is not fully reproduced in malnourished co-gavaged mice

We anticipated that the hypothesized bacterial colonization of jejunum mucosa is too rare in healthy mice to be detected in practice, so we investigated the published mouse model of environmental enteropathy (EE) [20] in which mice were weakened by malnutrition and orally challenged with a cocktail of *E. coli* and *Bacteroides/Parabacteroides* spp. Briefly, at 21 days of age, mice were placed on either a malnourishing (MAL) or a nutritionally complete (COM) control diet (Fig. 1A). Diet formulations were identical to those in the original EE study [20] except that food dyes were omitted to minimize sample autofluorescence during imaging. The human gut bacterial isolates – a mixture two *E. coli* isolates, and a mixture of five *Bacteroides/Parabacteroides* spp. isolates (*Bacteroides dorei*, *Bacteroides fragilis, Bacteroides ovatus, Bacteroides vulgatus*, and *Parabacteroides distasonis*) – were identical to the previously used isolates [20]. During the third week on the experimental diets, mice were orally gavaged three times (once every other day) with one of the following bacterial cocktails: *E. coli* and *Bacteroides/Parabacteroides* spp. mixture (EC&BAC), *E. coli* only (EC), *Bacteroides/Parabacteroides* spp. only (BAC), or phosphate buffered saline (PBS) as a control (Fig. 1A). During the fifth week, mice were euthanized (Fig. 1A), and small and large intestines were analyzed. Analysis of the LI was included because colonic microbiota may impact SI microbiota as a result of retrograde transport from cecum to ileum [37] or coprophagy (fecal ingestion) [38].

**Figure 1.**
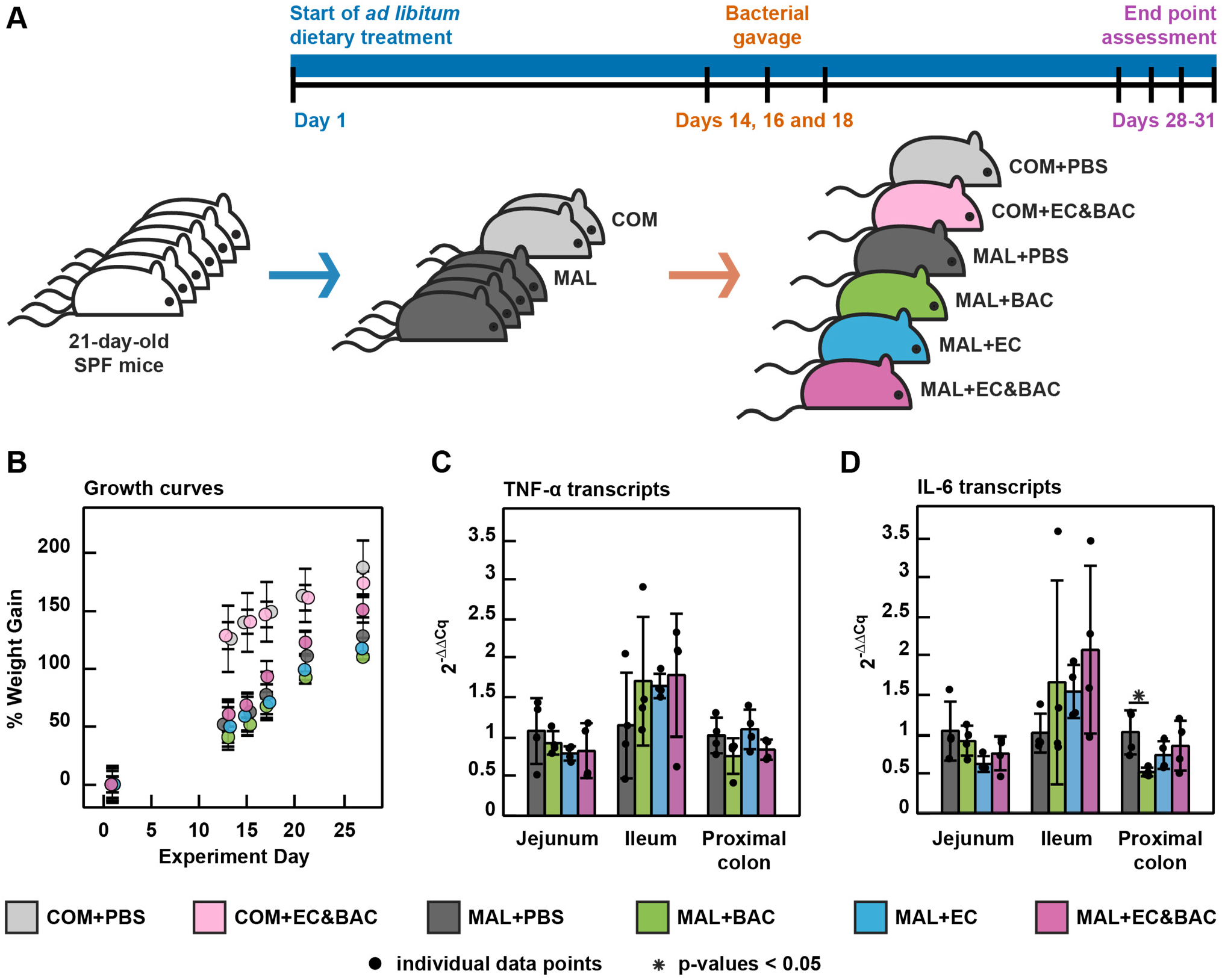
Study design and mouse response to dietary and bacterial interventions. (A) The setup of the animal experiment. SPF mice at 21-days-of-age were placed on one of the experimental diets on day 1 of the experiment and then gavaged with bacterial cocktails or PBS control on days 14, 16, and 18. Mice were euthanized for examination on days 28, 29, 30, and 31. COM: nutritionally complete diet. MAL: malnourishing diet low in protein (7%) and fat (5%), but containing the same amounts of calories, vitamins, and minerals on the basis of mass. PBS: phosphate buffered saline gavage as a no-bacteria control. BAC: gavage with five *Bacteroides/Parabacteroides* spp. isolates. EC: gavage with two *E. coli* isolates. EC&BAC: gavage with all seven bacterial isolates. (B) Mean weight gain data showing how malnutrition and bacterial gavage affected animal growth. Bars are ±S.D. (C–D) Relative quantification of TNF-α (C) and IL-6 (D) transcripts normalized to GAPDH housekeeping gene and with respect to the MAL+PBS group[39] using a TaqMan assay. Transcripts were quantified in the jejunum (2nd quartile of the SI), ileum (4th quartile of the SI), and proximal colon tissues in malnourished mice only. Each group contained four mice, which were euthanized over the course of 4 days, one mouse per group per day. The averages of the biological replicates (each of which is the average of triplicate qPCR runs) are plotted. Kruskal–Wallis test was used to evaluate statistical significance between treatments within each tissue type, and Benjamini–Hochberg correction was used to account for multiple comparisons.

Consistent with the original EE study [20], all MAL diet treatments resulted in retarded growth; for example, mice on the MAL diet gained 2.5 times less weight than mice fed the COM diet by day 13 (Fig. 1B). However, in contrast to the original EE study [20], oral gavage of bacteria did not further retard weight gain nor lead to weight loss (Fig. 1B). Further, among the four MAL groups, differences in the transcription of inflammatory cytokines tumor necrosis factor alpha (TNF-α) and interleukin 6 (IL-6) in jejunal, ileal, and colonic tissues were modest, and most differences were not statistically significant (Fig. 1 C-D). Therefore, the previously reported [20] wasting and intestinal inflammation characteristics of EE were not reproduced.

### Gavaged bacterial isolates persist in jejunum digesta of malnourished co-gavaged mice

Although the published EE phenotype was not fully reproduced, we hypothesized that the gavaged *E. coli* and *Bacteroides*/*Parabacteroides* spp. isolates may have still colonized the jejunum mucosa of malnourished mice. We used 16S rRNA amplicon sequencing to inform whether bacterial colonization of jejunum mucosa was plausible. If gavaged isolates colonized jejunum mucosa, we expected to detect their signatures in jejunum lumenal contents (hereafter digesta) or feces. To test whether the gavaged bacterial isolates were still present in the mouse gut 12-14 days after gavage, we performed 16S rRNA gene amplicon sequencing (Fig. 2) and compared the obtained amplicon sequence variants (ASV) to the full 16S rRNA gene sequences of the gavaged isolates. We identified four unique ASVs assigned to *Bacteroidaceae*,the family of gavaged *B. fragilis, B. dorei, B. ovatus*, and *B. vulgatus* isolates (Table S1). Of these four ASVs, one ASV was assigned to the resident *B. thetaiotaomicron* (hereafter *B. theta*) species and was present in all six groups, whereas three ASVs aligned perfectly with a region of full 16S rRNA sequences of the gavaged *Bacteroides* spp. and were present only in groups gavaged with *Bacteroides* spp. (Table S1). Furthermore, we identified one ASV assigned to *Tannerellaceae* (Table S1), the family of the gavaged *P. distasonis* isolate, and this ASV aligned perfectly with a region of full 16S rRNA sequence of the gavaged *P. distasonis* isolate and could only be detected in mice gavaged with *P. distasonis* (Table S1 and Fig. 2A). Both *Bacteroidaceae* and *Tannerellaceae* were detected more abundantly in feces than in jejunum lumenal contents (hereafter jejunum digesta) (Fig. 2A). Finally, we detected one ASV assigned to *Enterobacteriaceae* (Table S1), the family of the gavaged *E. coli*, and this ASV aligned perfectly with a region of full 16S rRNA sequences of the gavaged *E. coli* isolates. Remarkably, this ASV was only detected in MAL+EC&BAC mice (but not in mice gavaged solely with *E. coli*), in both feces and jejunum digesta (Fig. 2A). We concluded that gavaged bacterial isolates persisted in the gut for at least 12-14 days.

**Figure 2.**
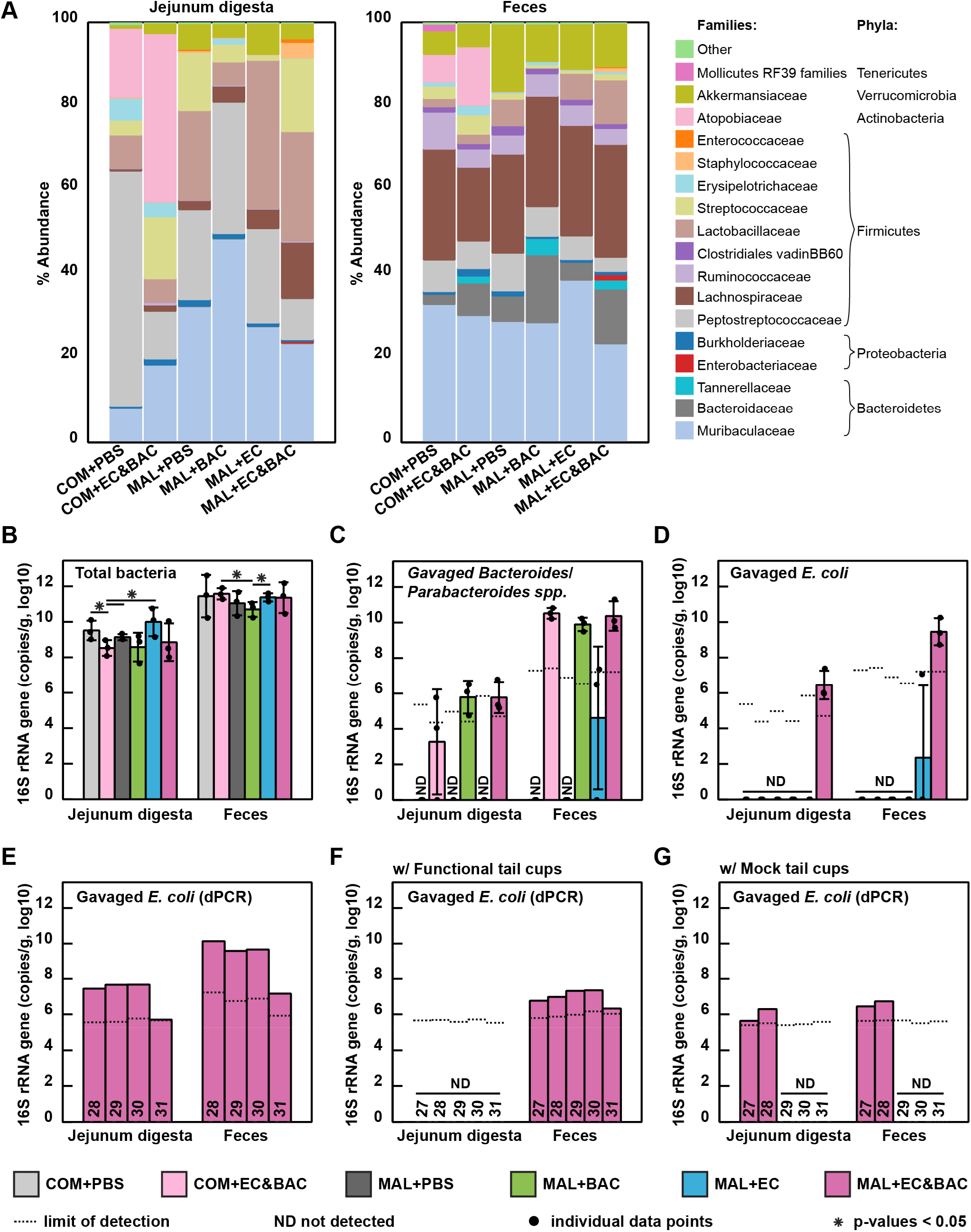
Relative and absolute bacterial 16S rRNA gene copy abundance in jejunum digesta and feces across treatment groups. (A) Relative family-level microbiota composition profiled by 16S rRNA gene amplicon sequencing. (B) Log-absolute total 16S rRNA gene copy abundance quantified by digital PCR (dPCR). (C-D) Extrapolation of log-absolute 16S rRNA gene abundance of (C) gavaged *Bacteroides/Parabacteroides* spp. and (D) gavaged *E. coli*. In A-D, the analysis consisted of six groups: control mice not gavaged with bacteria (COM+PBS), control mice gavaged with an *E. coli* and *Bacteroides/Parabacteroides* spp. cocktail (COM+EC&BAC), malnourished mice not gavaged with bacteria (MAL+PBS), malnourished mice gavaged only with *Bacteroides/Parabacteroides* spp. isolates (MAL+BAC), malnourished mice gavaged only with *E. coli* isolates (MAL+EC), and malnourished mice gavaged with the full bacterial cocktail (MAL+EC&BAC). Each group contained three mice, which were euthanized on days 28, 29, and 30, with one mouse per group per day. Limit of detection was expressed as group average. Statistical significance was evaluated using Kruskal-Wallis tests. (E-F) Log-absolute 16S rRNA gene copy abundance of gavaged *E. coli* quantified by dPCR with *Enterobacteriaceae* primers in MAL+EC&BAC group (E) without tail cups (identical biological replicates as shown in A-D), (F) with functional tail cups that effectively prevented coprophagy, and (G) with mock tail cups that did not prevent coprophagy but recapitulated the stress of wearing them. In E-G, each bar represents an individual mouse euthanized on the specified day of the experiment (27-31).

Knowing that gavaged bacterial isolates persisted in the gut for 12-14 days after gavage, we next asked whether their presence depended on each other (co-gavage) or on malnutrition, suggestive of a potential synergy. We quantified the absolute abundance of each microbial taxon because, unlike relative-abundance quantification, absolute abundance quantification is not biased by differences in total bacterial loads or the loads of other taxa [38, 40, 41]. Following our lab’s quantitative sequencing protocol [38, 40, 41], the fractional composition obtained by 16S rRNA gene amplicon sequencing was anchored by the absolute load of total bacterial 16S rRNA gene copy numbers in each sample (Fig. 2B). Consistent with other reports [7, 8], total bacterial loads were lower in the jejunum digesta than in feces. Among fecal samples, only the MAL+BAC group displayed a significant 4-fold lower total abundance relative to the COM+EC&BAC and MAL+EC groups (Fig. 2B). Among jejunum digesta samples, only the COM+EC&BAC group exhibited significantly lower total 16S rRNA gene copy loads than the three groups not exposed to *Bacteroides*/*Parabacteroides* spp. during gavage (Fig. 2B). Therefore, absolute quantification captured differences in total bacterial loads along the gut and established that neither malnutrition nor bacterial gavage introduced stark differences in total bacterial loads across treatment groups.

We next investigated how malnutrition and co-gavage affected the absolute loads of the gavaged bacteria in the jejunum digesta and feces. Although 16S rRNA gene amplicon sequencing is generally not suitable for species-level analysis [42], the well-defined *Bacteroides* species pool in the non-gavaged mice and the availability of complete 16S rRNA gene sequences of gavaged isolates permitted it in this situation. We found that, in feces, neither the MAL diet nor co-gavage with *E. coli* were required for gavaged *Bacteroides*/*Parabacteroides* spp. to reach abundances comparable to the levels of resident *B. theta*, which was present at consistent abundance across all six treatments (Fig. 2C and Table S1). However, in the jejunum digesta, the two MAL treatments that received the *Bacteroides*/*Parabacteroides* spp. gavage had the greatest abundance of *Bacteroides*/*Parabacteroides* spp. (10^6^ 16S rRNA gene copies per gram of jejunum digesta) (Fig. 2C and Table S1). In contrast, both the MAL diet and co-gavage with *Bacteroides/Parabacteroides* spp. were required for *E. coli* to colonize the mice and reach 10^6^ – 10^7^ (in jejunum digesta) and ~10^10^ (in feces) 16S rRNA gene copies per gram (Fig. 2D and Table S1). We confirmed the latter findings by digital PCR (dPCR) with *Enterobacteriaceae* primers, which detected *E. coli* in MAL+EC&BAC group at 2 orders of magnitude greater abundance than in COM+EC&BAC or MAL+EC groups (Fig. S1A and Table S2). However, both 16S rRNA gene amplicon sequencing and dPCR with *Enterobacteriaceae* primers showed that *E. coli* loads in MAL+EC&BAC group decreased by day 31 of the experiment, dropping below the limit of detection (LOD) in 16S rRNA gene amplicon sequencing (Fig. S1D and Table S1) and by 2 orders of magnitude in dPCR (Fig. 2E, Fig. S1C, and Table S2); this drop motivated us to only consider mice euthanized on days 28-30 to calculate group averages (Fig. 2, A-D, and Fig. S1, A and B). We made two observations in support of synergy: (1) *Bacteroides/Parabacteroides* spp. required malnutrition to persist for 2 weeks in the jejunum, and (2) *E. coli* required both malnutrition and co-gavage with *Bacteroides*/*Parabacteroides* spp. to persist in the jejunum and other parts of the gut.

The bacterial loads we observed in the jejunum were orders of magnitude higher than what has been reported in the human jejunum [7, 8], likely due to coprophagy [38], a common behavior in lab mice. Thus, we next wished to determine whether coprophagy might be contributing to the differences we observed among the gavage treatments. Reingestion of feces could further alter the SI microbiota composition in the gavage treatments by continuously re-introducing the gavaged isolates from the feces to the jejunum. To test this potential effect, we leveraged mouse tail cups previously designed in our lab to prevent coprophagy and eliminate its effect on SI microbiota [38]. In this experiment, MAL+EC&BAC treatment was split across two groups after the gavage: one group received functional tail cups that prevented fecal reingestion (Fig. 2F), whereas the other group received mock tail cups that permitted coprophagy but recapitulated the stress of wearing them (Fig. 2G). In both groups, the previously observed high *E. coli* loads were not recapitulated (Fig. 2, E-G), possibly because tail-cup intervention that involved wearing bulky tail cups and being socially isolated altered mouse physiology [38]. In the functional tail-cup group, *E. coli* could be detected in feces but not jejunum digesta by dPCR (Fig. 2F), suggesting that coprophagy played a role in *E. coli* persisting and reaching high loads in the jejunum digesta. In contrast, in the mock tail-cup group, *E. coli* could be detected in both feces and jejunum digesta but only at earlier time points (Fig. 2G). At later time points, *E. coli* loads in both feces and jejunum digesta dropped below the LOD (Fig. 2G), and such drop may be related to the previously observed drop in *E. coli* loads by day 31 of the experiment (Fig. 2E). Overall, the tail-cup intervention suggested that the continuous re-exposure to *E. coli* and *Bacteroides*/*Parabacteroides* spp. via fecal reingestion enabled *E. coli* to persist and reach high loads in the jejunum digesta of malnourished co-gavaged mice.

### Whole-tissue 3D imaging can detect and profile bacteria remaining in the jejunum after digesta passage

Having established that persistence of gavaged bacteria in jejunum digesta may have been the result of fecal reingestion and not necessarily bacterial colonization of jejunum mucosa, we next wished to test for evidence of bacterial resistance to washout with digesta passage. The SI is not continuously filled with digesta (Fig. 3A); potentially, such digesta separation marks meals ingested at different times. We reasoned that bacterial retention after digesta passage would indicate bacterial resistance to washout, likely by adherence to and colonization of jejunum mucosa. Thus, if bacteria were able to resist washout, we should detect them in the empty segments of the SI, whereas in segments containing digesta it would be difficult to differentiate bacteria moving with the digesta from those being retained in the SI [25].

**Figure 3.**
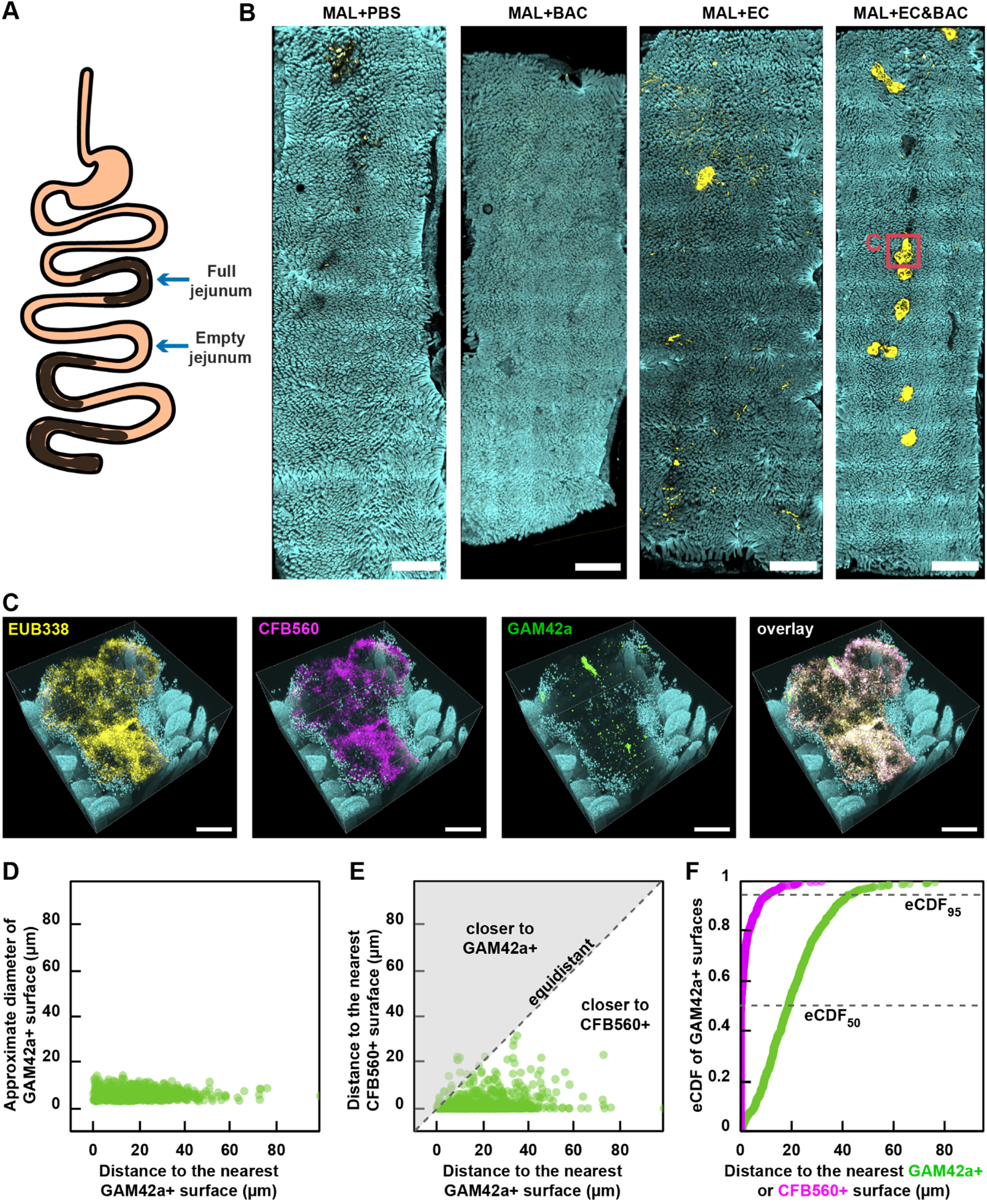
Large-scale 3D imaging of bacteria in empty jejunum segments and image analysis of bacterial distribution. (A) A diagram illustrating the movement of digesta between empty jejunum segments. Empty jejunum segments were isolated from the middle third of the SI, with the exact sampling location depending on the distribution of digesta along the SI. (B) Large-scale low-magnification fluorescence tile scans of whole empty jejunum segments showing DAPI staining of host nuclei that marks epithelial surface (cyan) and HCR v2.0 staining of total bacteria with EUB338 probe (yellow) across malnourished (MAL) treatment groups. All scale bars 2 mm. (C) High-magnification 3D fluorescence image of a large surface aggregate seen in B (red square) additionally showing HCR v2.0 staining of *Bacteroidetes* with CFB560 probe (magenta) and *Gammaproteobacteria* with GAM42a probe (green). Two large surface aggregates in one MAL+EC&BAC mouse were tile-scanned in 3D, with four fields of view per scan. Only one aggregate is displayed in C, but both are analyzed in panels D-F. All scale bars 200 μm. (D-F) Image analysis of *Gammaproteobacteria* distribution in large surface bacterial aggregates seen in MAL+EC&BAC mouse. (D) The size of GAM42a+ surfaces plotted against their shortest distance to another GAM42a+ surface. (E) The shortest distance of GAM42a+ surfaces to CFB560+ surface plotted against their shortest distance to another GAM42a+ surface. (F) Empirical cumulative distribution function (eCDF) of GAM42a+ surfaces as a factor of the shortest surface-to-surface distance to either GAM42a+ (green) or CFB560+ (magenta) surfaces. In D-F, the distance was expressed as the shortest surface-to-surface distance.

We used our 3D imaging tools [34] to map the exact bacterial location with respect to the complex mucosal landscape of the host and look for evidence of bacterial adhesion to the mucosa. First, to evaluate imaging workflow, we performed a pilot experiment by taking one empty jejunum tissue sample from each malnourished group through the whole-tissue 3D imaging pipeline (Fig. 3). Briefly, empty jejunum segments identified in the middle third of the SI were embedded into an acrylamide hydrogel in whole-mount, the resulting hydrogel-tissue hybrids were permeabilized with lysozyme for bacterial staining, cleared with SDS, and stained with DAPI for DNA (which marks epithelium), and HCR v2.0 for bacteria with one universal EUB338 probe and two taxon-specific CFB560 and GAM42a probes (see Methods for details). In our study, taxon-specific probes recognized not only the gavaged isolates but also resident taxa. For example, the *Bacteroidetes-specific* CFB560 probe only targeted the *Bacteroidales* order, including *Bacteroidaceae*, *Tannerellaceae* and *Muribaculaceae* families, without a mismatch and *Lachnospiraceae* family with one mismatch. Similarly, *Gammaproteobacteria*-specific GAM42a probe only targeted the *Enterobacteriaceae* family, solely represented by gavaged *E. coli*, without a mismatch and the resident *Burkholderiaceae* family with one mismatch.

Whole-tissue samples were first tile-scanned using a confocal fluorescence microscope at low magnification (5x) for DAPI staining of epithelium and HCR v2.0 staining of total bacterial (Fig. 3B). Despite high total bacterial loads in jejunal digesta (10^8^-10^10^ 16S rRNA gene copies per gram; Fig. 2B), empty jejunum tissue segments were mostly devoid of bacteria (Fig. 3B). Bacteria were predominantly observed in scattered aggregates, which were detected in MAL+PBS, MAL+EC, and MAL+EC&BAC mice (Fig. 3B). These aggregates showed stronger eubacterial signal in in MAL+EC and MAL+EC&BAC mice and were more abundant in MAL+EC&BAC mouse, suggestive of bacterial retention after digesta passage (Fig. 3B). We imaged two of these large surface aggregates from the MAL+EC&BAC mouse at higher magnification (20x), revealing that these aggregates contained not only bacteria but also host cells (Fig. 3C, Fig. S2, and Fig. S3). Imaging at higher magnification with taxonomic resolution of two large surface aggregates further revealed that CFB560 and GAM42a targets were both present in the aggregates (Fig. 3C, Fig. S2, and Fig. S3). To assess the abundance of each target, the images were segmented in Imaris to obtain surfaces positive for the EUB338, CFB560, and GAM42a probes and the segmented surfaces were filtered to remove noise (Fig. S4–S6). Quantification of segmented and filtered surface volume revealed that, averaged over two aggregates, CFB560+ and GAM42a+ volume fractions were 35% and 0.5%, respectively. These abundance estimates were in agreement with 16S rRNA gene amplicon sequencing, which detected *Bacteroidales* and *Enterobacteriaceae* in the jejunum digesta of MAL+EC&BAC treatment at 20–27% and 0.2–0.9% relative abundance, respectively (Fig. 2A). This experiment confirmed that the imaging workflow enabled us to visualize bacteria retained in the empty jejunum after digesta passage and that taxon-specific probes could identify the taxonomic groups of the retained bacteria.

Sequencing had suggested that *E. coli* required co-gavage with *Bacteroides/Parabacteroides* spp. to persist for at least 12-14 days in the jejunum of malnourished mice (Fig. 2D), suggesting potential synergy. Thus, next we analyzed spatial structure of taxonomic groups within the bacterial aggregates to determine whether GAM42a+ bacteria were well-mixed with or segregated from CFB560+ bacteria (Fig. 3, D–F). We found that GAM42a+ bacteria existed predominantly as single cells separated from each other by a distance larger than the estimated cell diameter (Fig. 3D). In contrast, CFB560+ bacteria were so tightly packed that they could not be segmented into individual cells (Fig. 3C and Fig. S5). GAM42a+ bacteria associated more closely with CFB560+ bacteria than with each other (Fig. 3, E–F), with 95% of GAM42a+ bacteria located less than 11 μm away from CFB560+ bacterial surface but less than 44 μm away from GAM42a+ bacterial surface (Fig. 3F). Such close association of GAM42a+ bacteria with CFB560+ bacteria supported the dependency of *E. coli* on *Bacteroides/Parabacteroides* spp., as previously hypothesized[21].

### Bacterial retention is most prevalent in malnourished and co-gavaged mice

Having established that whole-tissue 3D imaging can detect and profile bacterial retention after digesta passage, we next set out to answer three biological questions: (1) Does the MAL+EC&BAC group retain more bacteria in the jejunum after digesta passage than the other MAL groups? (2) Can bacteria closely associate with the jejunum mucosa, which is indicative of bacterial colonization of the jejunum mucosa? (3) What are some possible host responses to bacterial association with the mucosa in the jejunum? To answer these questions, we further improved experimental design and whole-tissue 3D imaging tools.

First, to reduce potential sampling bias related to the timing of food ingestion, in this experiment, the mice were fasted for 1 h prior to euthanasia and sample collection. Because feeding behavior may be regulated by circadian rhythms [43], all mice were also euthanized at the same time of day. Furthermore, transcardial perfusion of the vasculature (used to eliminate blood, which is strongly autofluorescent) can cause the gut to swell [44], potentially disrupting mucosal colonizers, so we omitted transcardial perfusion in this experiment. To minimize amplification of background and false-positive HCR signal, which increases with depth and complicates specific detection of mucosal colonizers deep in the mucosa, we adopted HCR v3.0 technology [30]. Compared with HCR v2.0, these HCR v3.0 probes reduce off-target signal amplification and increase the signal-to-background ratio [30]. We designed a universal degenerate HCR v3.0 probe set (Fig. S7) that, compared with HCR v2.0 EUB338 probe, successfully suppressed background signal amplification in the gut tissue (Fig. S8). We also designed two new HCR v3.0 probes to specifically target either the gavaged *E. coli* or *Bacteroides/Parabacteroides* spp. isolates without a mismatch using their full 16S rRNA sequences (Fig. S9); these probes successfully resolved false-positive signal amplification at depth associated with CFB560 probe (Fig. S10). In our study, *E. coli* probe only recognized the gavaged *E. coli* isolates. In contrast, *Bacteroides*/*Parabacteroides* spp. probe recognized not only the gavaged *Bacteroides*/*Parabacteroides* spp. isolates but also the resident *B. theta* without a mismatch and the resident *Muribaculaceae* with one mismatch; thus, this probe was referred to as *Bacteroidales* probe hereafter. Finally, to improve antibody entry into the hydrogel-tissue hybrids, we adopted a new more porous tissue preservation chemistry for antibody staining (Fig. S11).

With these new advances, in this experiment we again observed that large surface aggregates were present in empty jejunum segments, and these aggregates were most common in the MAL+EC&BAC group (3 out of 4 mice had aggregates), followed by the MAL+BAC group (1 out of 4 mice had aggregates). None of the mice in the MAL+PBS or the MAL+EC groups had aggregates when they were fasted for 1 h (Fig. 4A), which was in contrast to the nonfasted mice in these same treatments (Fig. 3B). These observations support our prediction that fasting reduces sampling bias with respect to the timing of food ingestion. Notably, these aggregates were opaque and large enough to be visible to the naked eye after tissue clearing (Fig. 4A). To confirm that bacteria were present in these aggregates, cleared hydrogel-tissue hybrids were stained with DAPI for epithelium and with HCR v3.0 for total bacteria, *Bacteroidales*,and *E. coli* (Fig. 4B and Figs. S12–S27). Low-magnification (5x) tile-scanning of whole-tissue samples for epithelium and total bacteria detected bacteria only in samples with large surface aggregates (Fig. 4, A–B, and Figs. S12–S27). Bacteria were predominantly found in large surface aggregates; only one mouse in the MAL+EC&BAC treatment additionally showed small bacterial clusters on the tissue (Fig. 4B and Fig. S24). Hydrogel-tissue hybrids from two mice per group were also imaged at high magnification (20x) and with taxonomic resolution (Fig. 4C and Figs. S28-S35). High-magnification images were acquired at the positions of large surface aggregates except for samples where large surface aggregates could not be detected. After thorough exploration, tissue samples lacking aggregates were imaged at random, representative locations. At high magnification, bacteria in large surface aggregates were clearly visible even in areas where they appeared dim at low magnification (Fig. 4, B (purple squares) and C). The samples that contained large surface aggregates retained the greatest amounts of bacteria after digesta passage, whereas the samples lacking these surface aggregates contained only a few sparse bacterial cells (Fig. 4D). This experiment suggests that, in malnourished mice not gavaged with bacterial isolates, the jejunum mucosa was devoid of bacteria after a 1-hour fast, whereas in malnourished mice gavaged with the EC&BAC bacterial cocktail, the jejunum mucosa retained bacteria.

**Figure 4.**
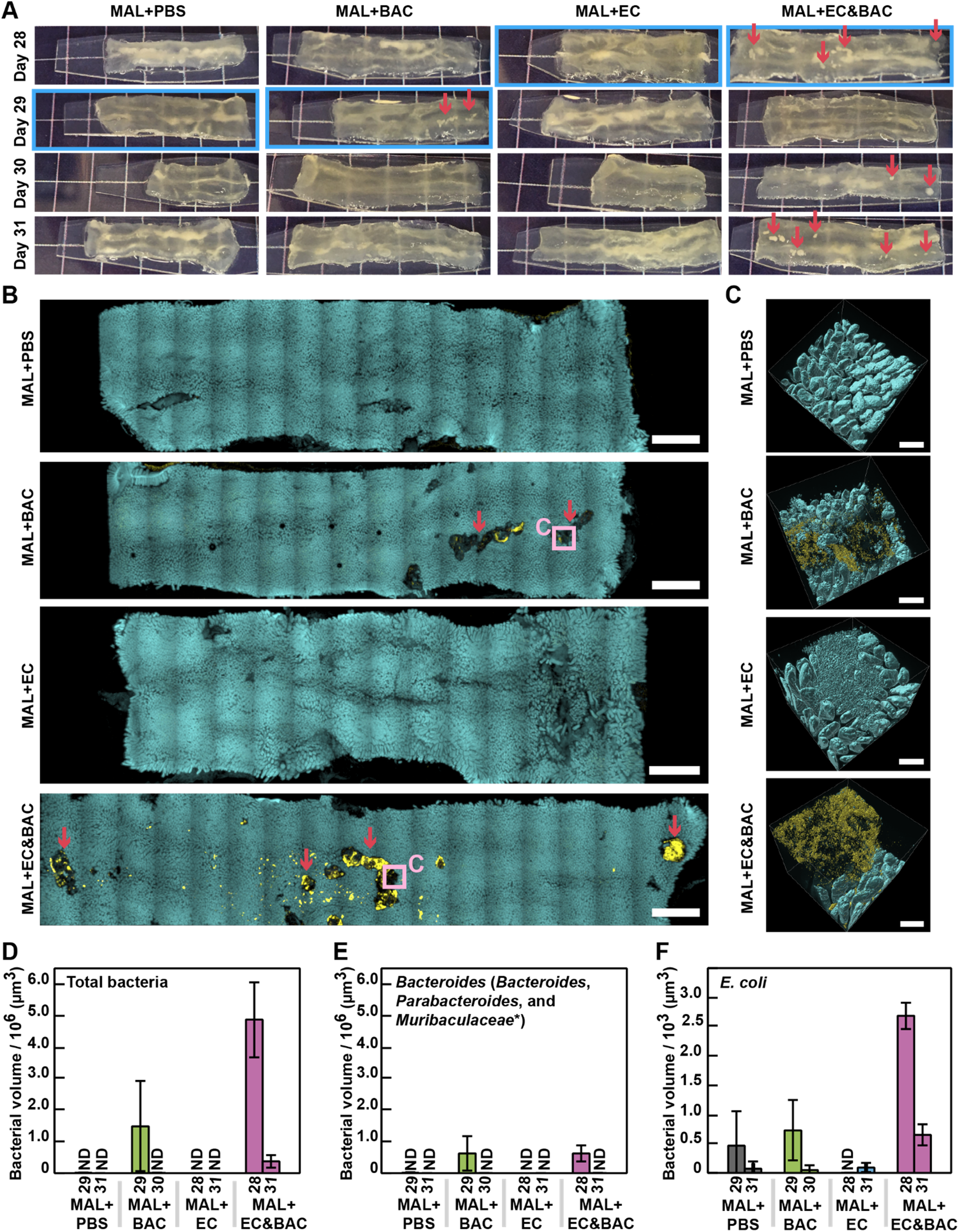
Large-scale 3D imaging of bacteria in empty jejunum segments after 1-h fast and quantification of bacterial retention. (A) Photos of empty jejunum segments after clearing. Large surface aggregates indicative of bacteria remained opaque after clearing and were visible by eye (red arrows). The mice were euthanized over the course of 4 consecutive experimental days with one mouse per group per day. All mice were fasted for 1 h prior to euthanasia and sample collection and all procedures began at 11 am. Empty jejunum segments were identified in the 3^rd^ quartile of the SI. Panels B and C display only one example tissue per group (samples indicated by the blue rectangles in panel A). (B) Example large-scale low-magnification fluorescence tile scans of whole empty jejunum segments showing DAPI staining of epithelial surface (cyan) and HCR v3.0 staining of total bacteria (yellow) across malnourished treatment groups. The rest of the tile scans are provided in Figs. S12–S27. Opaque large surface aggregates observed in panel A can also be identified in panel B (red arrows). Large surface aggregates (violet squares) were selected for high-magnification imaging. All scale bars 2 mm. (C) Example segmented and filtered high-magnification 3D fluorescence images of the same empty jejunum segments displayed in B showing DAPI staining of host nuclei that marks epithelial surface (cyan) and HCR v3.0 staining of total bacteria (yellow). Tissue samples were also imaged for HCR v3.0 staining of *Bacteroidales* and *E. coli*, but only total bacteria are displayed in C. Two mice from each group were imaged at high magnification; for each mouse, three 3D images were acquired with four tiles per image. If present, aggregates were imaged. Additional images, with one example image per mouse, are provided in Figs. S28-S35. All scale bars 200 μm. (D-F) Quantification of (D) total bacteria, (E) *Bacteroidales*, and (F) *E. coli* volumes in high magnification 3D images. X-axis specifies the treatment group and the day of euthanasia of mice under consideration. Y-axis shows bacterial volumes averaged over three 3D images for each mouse (N = one mouse per treatment). The star denotes probe targets with one mismatch. ND: no significant amount of bacteria detected.

Next, to test whether the gavaged bacterial isolates were present in the aggregates suggestive of their resistance to washout with digesta passage, we analyzed high-magnification images for the presence of *Bacteroidales* and *E. coli* (Fig. 4, E-F). *Bacteroidales* were detected in large surface aggregates in both MAL groups gavaged with *Bacteroides*/*Parabacteroides* spp. (specifically, in one MAL+BAC mouse and one MAL+EC&BAC mouse (Fig. 4E)). In contrast, *E. coli* could be detected above the background noise only in one MAL+EC&BAC mouse (Fig. 4F). Inability to detect *E. coli* in the MAL+EC&BAC day 31 mouse (Fig. 4F) was in line with the 16S rRNA analysis, which suggested that *E. coli* loads fell below the LOD of sequencing by day 31 of the experiment (Fig. 2E). *E. coli* was not found in any of the MAL+EC mice (Fig. 4F). In the two MAL+BAC and MAL+EC&BAC mice in which *Bacteroidales* were abundant, *Bacteroidales* represented 42% and 13% of total bacterial volume, respectively (Fig. 4, D-E), whereas in one MAL+EC&BAC mouse where *E. coli* was detected above background noise, *E. coli* amounted to 0.06% of total bacterial volume (Fig. 4, D and F). In comparison, 16S rRNA gene amplicon sequencing detected *Bacteroidales* at 11%–73% of total 16S rRNA load in the jejunum digesta across all mice on MAL diet, and *E. coli* at 0.2–0.9% of total 16S rRNA load in the jejunum digesta across mice in MAL+EC&BAC treatment (Fig. 2A). Overall, imaging of bacterial aggregates in digesta-free jejunum sections was able to detect *Bacteroidales* (which includes gavaged *Bacteroides* spp.) in MAL+EC&BAC and MAL+BAC groups, as well as gavaged *E. coli* in MAL+EC&BAC group, which supported our prediction that gavaged bacterial isolates resisted washout with digesta passage in these groups.

### Bacterial association with jejunum mucosa is detected in malnourished co-gavaged mice

Next, we asked whether the bacteria retained in the jejunum of the MAL+EC&BAC group after digesta passage showed evidence of colonization of the jejunum mucosa (e.g. consistent spatial association of microbes with mucosa). The published EE model detected bacteria by 2D imaging in between the villi in the jejunum of malnourished mice and malnourished mice co-gavaged with *E. coli* and *Bacteroides/Parabacteroides* spp. [20]; however, the images were acquired at locations with digesta. Where digesta is present, bacterial detection in the intervillus spaces could also be explained by partitioning of lumenal bacteria to the mucosa [25] or by sectioning artifacts that could physically push lumenal bacteria into the intervillus spaces. Fasting animals prior to euthanasia and selecting empty jejunum segments provided a unique opportunity to detect and visualize in 3D bacterial colonization of SI mucosa not biased by the presence of digesta.

Imaging revealed evidence for bacterial association in the jejunum mucosa of malnourished co-gavaged mice. We zoomed in to examine small bacterial clusters detected at low magnification (5x) in one of the MAL+EC&BAC mice (Fig. 5A) at higher magnification (20x). We observed that these dense clusters of bacteria penetrated deeply in between the villi and, in some areas, even appeared to anchor to and wrap around the villi (Fig. 5B and Video S1). Such high local bacterial density in the jejunum mucosa contrasted with the previously reported sparse distribution of individual cells and small aggregates in the jejunum mucosa of SPF mice [33]. Although the mixing of lumenal contents by peristalsis [25] may have moved these aggregates from the lumen into the mucosa, the high bacterial density and abundance of aggregates across the SI mucosa only in this treatment suggested bacterial growth in the mucosa. This single observation does not prove mucosal colonization, however, it shows that bacteria can be abundant in the intervillus spaces of empty jejunum segments (independent of the digesta presence) and that our experimental design and imaging technology were able to capture such events.

**Figure 5.**
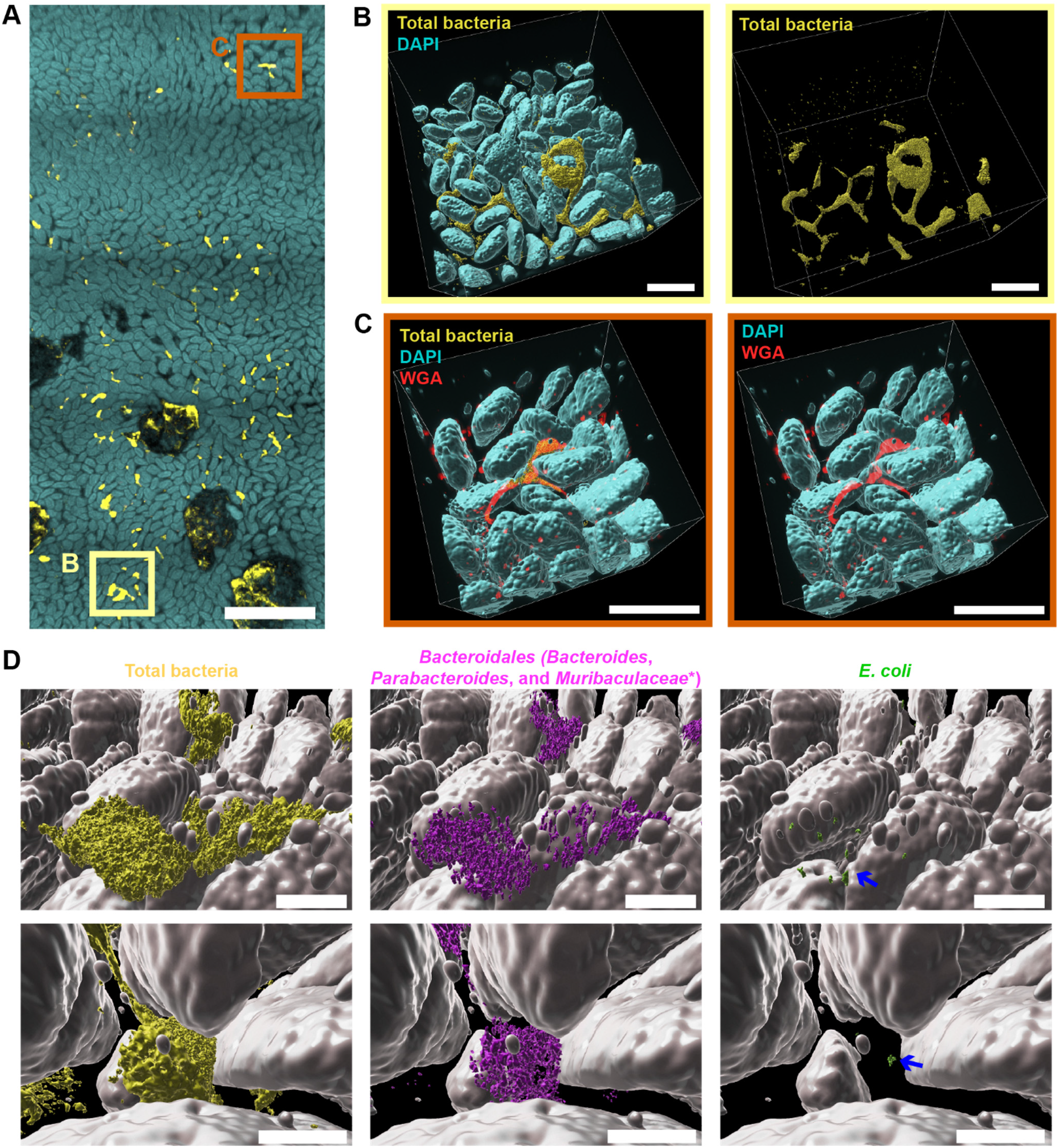
Large-scale 3D imaging of bacterial colonization of SI mucosa in the empty jejunum of a MAL+EC&BAC mouse. (A) Large-scale low-magnification fluorescence tile scan of the empty jejunum segment from a MAL+EC&BAC mouse marking the positions of the zoomed-in 3D images in panels B (yellow square) and C (orange square). Cyan: DAPI staining of epithelium. Yellow: HCR v3.0 staining of total bacteria. Scale bar = 1 mm. (B) Segmented and filtered high-magnification 3D fluorescence image of a position highlighted in A showing DAPI staining of host nuclei that marks epithelial surface and HCR v3.0 staining of total bacteria (yellow). Left: overlay between DAPI and HCR v3.0 staining. Right: display of HCR v3.0 staining only. Three images were acquired, with four tile scans per image. One of the three images is displayed in B. All scale bars 200 μm. (C) Segmented and filtered high-magnification fluorescence image of a position highlighted in C additionally showing wheat germ agglutinin (WGA) staining of mucus and other glycans (red). Left: overlay between DAPI, HCR v3.0 and WGA staining. Right: overlay between DAPI and WGA staining only. Two images were acquired in spectral mode with one field of view per image. One of the two images is displayed in C. All scale bars 200 μm. (D) Zoomed-in display of mucosal bacteria in panel B additionally showing HCR v3.0 staining of *Bacteroidales* (magenta) and *E. coli* (green). Blue arrows point to the sparse *E. coli* cells. All scale bars 50 μm.

We next assessed the mucosal bacterial community detected in MAL+EC&BAC day 28 mouse for the presence of gavaged bacteria and host mucus (Fig. 5, C and D). *Bacteroidales* were abundant in the mucosa (Fig. 5D, center). Consistent with large surface aggregates (Fig. 4, D and E), *Bacteroidales* amounted to 14% of total bacterial volume in the mucosa. *E. coli* were rare in the mucosa (Fig. 5C, right), amounting to less than 0.1% of total bacterial volume. Notably, they were always found in close association with *Bacteroidales* (Fig. 5C, left); 80% of *E. coli* surfaces were located less than 6 μm away from *Bacteroidales*. We speculate that the ability of *Bacteroides* spp. to adhere to [45] and forage on [46, 47] mucus may enable them to colonize SI mucosa. Indeed, WGA staining for N-acetylglucosamine (which is abundant in mucus; [48, 49]) and HCR staining for bacteria showed that bacteria were co-localized with N-acetyleglucosamine (Fig. 5C). These results show that *Bacteroides/Parabacteroides* spp. colonization of the intervillus spaces of malnourished mice is possible, and that *E. coli* may closely associate with *Bacteroides/Parabacteroides* spp. in the mucosa.

### Loss of villi may be one consequence of bacterial colonization of jejunum mucosa

Finally, we investigated host response to bacterial colonization of jejunum mucosa. In our pilot experiment (Fig. 3C) and in the MAL+EC&BAC day 31 mouse (Fig. 6, A-C) we had observed that host nuclei seemed to associate with large surface bacterial aggregates. To better characterize this association, we stained large surface aggregates in the MAL+EC&BAC day 31 mouse with anti-EpCAM antibody for epithelial cells and anti-CD45 antibody for total immune cells (Fig. 6D). Free nuclei can represent epithelial damage (for example, villus blunting is characteristic of EE [20]), or immune-cell infiltration (for example, intraluminal neutrophil cast formation in response to *Enterobacteriaceae* overgrowth following acute *Toxoplasma gondii* infection [50]). Additionally, we stained the aggregates with WGA lectin that recognizes mucus; we predicted that mucus may also be present in large surface aggregates because it is known to mediate aggregation in the gut [51]. Based on visual inspection, antibody staining showed that epithelial cells were abundant in large surface aggregates whereas immune cells could be barely detected (Fig. 6D, i and iv). To quantify this difference, we selected a region of interest (ROI) in each image to include the large surface aggregates but exclude the surrounding villi and segmented the obtained ROIs in the antibody staining channel (Table S6). Normalized for ROI volume, we detected 2 orders of magnitude more antibody positive surfaces (by net volume) for anti-EpCAM staining than for anti-CD45 staining. Furthermore, in some fields of view, entire dislodged villi could be detected, again supporting that the observed free mammalian nuclei represented epithelial damage (Fig. 6D, iv). WGA staining was strong, suggesting that mucus associated with bacteria and cellular debris in large surface aggregates (Fig. 6D, ii and v). Overall, profiling of host components in large surface aggregates identified that dislodging villi may be one consequence of bacterial gavage and the subsequent bacterial colonization of jejunum mucosa.

**Figure 6.**
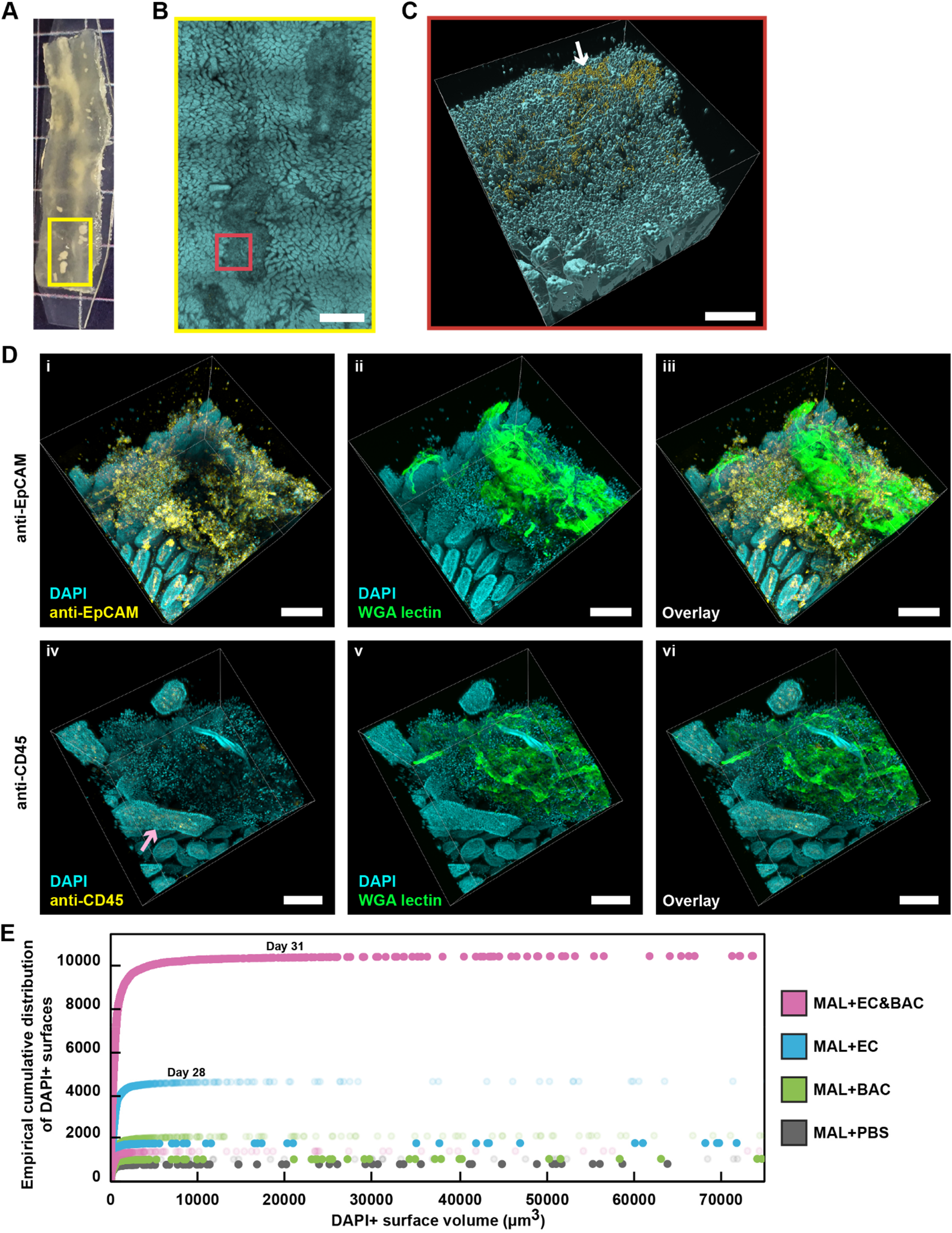
Large-scale 3D imaging and image analysis of a possible host response to bacterial colonization of SI mucosa. (A) A photo of a cleared empty jejunum segment from the MAL+EC&BAC day 31 mouse with large surface aggregates marking the position of the low-magnification tile scan shown in panel B. (B) Large-scale low-magnification fluorescence tile scan of the area highlighted in panel A displaying DAPI staining of epithelium (cyan) and HCR v3.0 staining of total bacteria (yellow) and marking the position of the high-magnification 3D fluorescence image shown in panel C (red square). At low magnification, bacteria appeared faint because of abundant mammalian nuclei in the large surface aggregates. Scale bars = 1 mm. (C) Segmented and filtered high-magnification 3D fluorescence image of the area highlighted in B displaying DAPI staining of the epithelium (cyan) and HCR v3.0 staining of total bacteria (yellow). Bacteria become clearly visible at high magnification (white arrow). Scale bars = 200 μm. (D) High-magnification 3D imaging of large surface aggregates detected in MAL+EC&BAC day 31 mouse for antibody and lectin staining of host components. Left (i and iv): Overlay between DAPI staining of the epithelium (cyan) and antibody staining of either epithelial cells with anti-EpCAM antibody (i; yellow) or total immune cells with anti-CD45 antibody (iv; yellow). Middle (ii and v): Overlay between DAPI staining of the epithelium (cyan) and WGA lectin staining of mucus (green). Right (iii and vi): Overlay between all three staining modalities. In some fields of view, entire dislodged villi could be detected (violet arrow). All scale bars 200 μm. (E) Quantification of DAPI+ surfaces as an empirical cumulative sum versus DAPI+ surface size. High-magnification images were segmented in Imaris to identify DAPI+surfaces, sorted based on size, and counted from the smallest (left) to the largest (right) objects. We deduce that objects on the far left represent individual nuclei, whereas objects on the far right represent intact villi. The same high-resolution images were analyzed as in Figure 4 D-F, with two mice per group and three images per mouse (for each mouse, empirical cumulative distribution was calculated over all three images). The values in the plot correspond to the number of DAPI+ surfaces (y-axis values) up to a certain size (x-axis values).

Finally, we quantified epithelial damage in the MAL groups, specifically the two groups in which epithelial damage was most pronounced: the MAL+EC and MAL+EC&BAC groups (Fig. 6E). High-magnification (20x) images were first segmented in Imaris to identify DAPI+ surfaces; the same images were analyzed as in Fig. 4, D-F, with two mice per group and three images per mouse. After segmentation, an empirical cumulative sum of DAPI+ surfaces was calculated over all three images from each mouse by sorting DAPI+ surfaces based on size and then counting them in the order from small to large (Fig. 6E). As suggested by the steeper early rise in the empirical cumulative count, small DAPI+ surfaces suggestive of free nuclei were more abundant in malnourished mice exposed to bacteria compared with the PBS treatment (Fig. 6E). Thus, quantitative analysis supported that bacterial gavage increased the extent of epithelial damage in malnourished mice.

## Discussion

Using a mouse model of environmental enteropathy and building on our whole-tissue clearing and 3D-imaging tools [34], we documented dense bacterial association with the jejunum mucosa, suggestive of colonization. We anticipated that bacterial colonization of jejunum mucosa would be rare across space and over time in healthy organisms, therefore, we leveraged three strategies. First, we reasoned that factors that weaken or overcome host defenses would be required to induce bacterial colonization of jejunum mucosa. We focused on EE mouse model, in which mice are stressed by a malnourishing diet and co-gavaged with a cocktail of *E. coli* and *Bacteroides*/*Parabacteroides* spp. [20]. Second, because rare events can be challenging to detect by 2D imaging of thin sections, we employed our whole-tissue 3D imaging tools to detect rare events over large areas of mucosal surface and with depth into the mucosa. Third, we performed a series of experiments to test whether bacterial colonization of jejunum mucosa was plausible in malnourished co-gavaged mice. Specifically, we used sequencing to test whether gavaged bacterial isolates persisted in the gut (jejunum digesta or feces) after the gavage. Then, we used whole-tissue 3D imaging to determine whether bacteria, including gavaged bacterial isolates, were retained in the jejunum after digesta passage. We reasoned that, if the gavaged bacterial isolates colonized jejunum mucosa, they should be detectable in the gut and should be able to resist washout with digesta passage.

With these approaches, we established that bacterial colonization of jejunum mucosa was plausible. Based on 16S rRNA gene amplicon sequencing, gavaged bacterial isolates persisted in the jejunum digesta 12-14 days after the gavage. Although tail-cup intervention established that this persistence required fecal reingestion and, as a result, may have been transient, whole-tissue 3D imaging suggested that gavaged bacterial isolates led to bacterial retention after digesta passage and were themselves (or their higher order taxonomic groups) retained in the empty jejunum. In contrast, in the empty jejunum of malnourished mice not gavaged with bacterial isolates, whole-tissue 3D imaging detected few to no bacteria, suggestive of effective bacterial clearance with digesta passage. In the empty jejunum of malnourished mice gavaged with *Bacteroides/Parabacteroides* spp., with or without *E. coli*, bacteria, if detected, appeared predominantly in large surface aggregates; possibly, these aggregates represented the clearance of remaining bacteria by the host rather than mucosal colonization. However, in the empty jejunum of malnourished co-gavaged group, dense bacterial clusters could be detected in the intervillus spaces suggestive of close bacterial association with and growth in the mucosa, further supporting that bacterial colonization of jejunum mucosa was plausible.

We acknowledge that bacterial colonization of jejunum mucosa or other mucosal surfaces is challenging to establish conclusively. Close spatial association can be evidence of colonization, e.g. mucosal colonizers should be present in mucosal crevices, such as crypts or intervillus spaces. Furthermore, mucosal colonizers should appear as multicellular clusters rather than individual cells, indicative of cellular division and growth in the mucosa. Here, we leveraged whole-tissue 3D imaging tools [34] that allowed us to map both microbe-microbe and host-microbe spatial organization at the mucosal interface over large areas and in 3D. For example, leveraging a protective surface gel [34], we successfully retained large surface aggregates and bacteria, which can otherwise be easily lost during sample processing and evade detection. Furthermore, we visualized in 3D how bacterial clusters connect in between and wrap around the villi. In contrast, 2D imaging of thin sections has failed to reveal such complex spatial structure in the same mouse model [20]. However, spatial association alone does not prove colonization because, although some lumenal microbes may be able to adhere to and grow in the mucosa, others may be present in the mucosa only transiently during the mixing of lumenal contents [25]. To prevent lumenal digesta from potentially biasing our imaging results and to test whether bacteria can remain associated with the mucosa after digesta passage, we fasted the mice for 1 h and selected for imaging only jejunum segments without digesta. Another benefit to this approach is that our detected bacterial retention after digesta passage suggested that any retained bacteria were resistant to washout, possibly by adherence to and/or colonization of the mucosa.

Although our results cannot prove bacterial colonization of jejunum mucosa or convey its extent due to the low number of biological replicates, our observation of this rare phenomenon uncovers important concepts toward understanding host-microbe interactions in the jejunum and SI in general. Specifically, we detected dense clusters of bacteria in association with the jejunum mucosa only in one (out of four) malnourished co-gavaged mice and none of the controls. Low incidence of bacterial colonization of jejunum mucosa raises questions about its true extent, however such data are challenging to capture because sufficient sampling and complete imaging of the large surface area of the SI is both time- and cost-prohibitive. Furthermore, bacterial colonization of jejunum mucosa can also be short-lived, for example, mucosal colonizers may disperse or be eliminated by host mucosal defenses. Possibly, the rare occurrence of jejunum colonization only in malnourished, co-gavaged mice is a concept that can help explain other host-microbe interactions in the SI. The low incidence of dense bacterial clusters in the jejunum mucosa could also be an artifact of mucosa contamination with lumenal bacteria. To mitigate potential contamination, we imaged empty jejunum segments without digesta after fasting the mice for 1 h. Indeed, the dense bacterial clusters we detected in the mucosa were a unique finding; previous 3D imaging reported only sparse bacterial cells in the jejunum mucosa of SPF mice [33]. Although mucosal bacteria cannot be fully uncoupled from lumenal bacteria, we speculate that understanding the relationship between microbiota of the lumen and the mucosa in full and empty gut segments alone can be valuable. For example, lumenal bacteria can appear in the mucosa during digestion phase, grow there after digesta passage, and be eliminated by the host or disperse upon arrival of the next meal. Characterizing these dynamics and how the host responds to and/or modulates them can improve our understanding of host-microbe interactions in the SI in both health and disease.

Along with other recent studies [21, 52], our work supports that complex microbe-microbe interactions can be responsible for disease. We made several observations in support of synergy. First, *E. coli* required both malnutrition and co-gavage with *Bacteroides*/*Parabacteroides* spp. to reach higher loads in the gut. Second, in malnourished co-gavaged mice, *E. coli* associated closely with *Bacteroidales* in the empty jejunum, in both large surface aggregates and mucosa. Although our observations of the spatial relationships between these taxa are consistent with aggregation of a well-mixed bacterial community dominated by *Bacteroidales*, it does not rule out bacterial interactions. Some potential mechanisms for this synergy are metabolic coupling [21, 47, 53], mucosal adhesion [45, 54], and modulation of host immune response [55, 56]. For example, our previous genome-scale metabolic models and *in vitro* bioreactor studies identified metabolic coupling between *Klebsiella pneumoniae*, a model *Enterobacteriaceae*, and *B. theta*, a model *Bacteroidaceae*, with *K. pneumoniae* consuming oxygen and protecting *B. theta* from oxygen stress and *B. theta* degrading complex carbohydrates and supplying simple sugars and short-chain fatty acids to *K. pneumoniae* [21]. Furthermore, previous *in vivo* studies showed that degradation of mucus and release of sialic acid by *B. vulgatus* supported *E. coli* growth in mouse colon [47], and evasion of colonic mucus by *B. fragilis* was required for *E. coli* to colonize colonic mucosa [54]. In support that similar mucus-mediated interactions may take place in jejunum mucosa, our imaging identified that WGA staining of N-acetylglucosamine and sialic acid abundant in mucus co-localized with dense bacterial clusters detected in the jejunum mucosa. Curiously, *Enterobacteriaceae* and *Bacteroidaceae* were reported to be both more prevalent and abundant in the jejunum of small intestinal bacterial overgrowth (SIBO) patients [57], although they did not co-occur in the jejunum of SIBO patients when different exclusion criteria were applied [58], or the duodenum of SIBO [35] or EE [59] patients.

We predicted that large, direct contact area between mucosal colonizers and jejunum mucosa, which is not protected by a dense mucus layer [5, 11, 60], would impact the host, and detected dislodged dispersing villi as one possible consequence to the host. If such villus loss exceeded villus regeneration, over time it would reduce the absorptive surface area and, at the same time, absorptive capacity of the SI. In fact, our observed villus loss in the context of malnutrition and in the presence of certain bacterial taxa may represent the mechanism of villus blunting and epithelial damage characteristic of EE [22–24]. This mechanism may also be relevant to celiac disease, which is characterized by villus blunting and malabsorption [61].

The combination of whole-tissue 3D imaging coupled with HCR and a strategic sampling of the SI have enabled the detection of rare mucosa-associated bacteria in the SI as well as potential changes to host physiology, which may be broadly relevant. The EE mouse model used here was originally developed to mimic the ingestion of fecal bacteria in contaminated food and water, a pressing problem in the developing world [23]. In the developed world, ingestion of enteropathogens and fecal bacteria with contaminated food and water is perhaps unlikely; however, fecal bacteria may reach the jejunum by retrograde transport [37], and a number of other factors can predispose to SIBO [7]. Therefore, the tools and approaches we used and the observations we made may provide cues toward the understanding not only of EE [22–24] but also other diseases of the SI, including SIBO [7, 57, 62], irritable bowel syndrome (IBS) [63], and celiac disease [61]. Therefore, we envision that the capacity to detect rare bacterial colonization of SI mucosa and profile microbe-microbe and host-microbe interactions at these locations can enable mechanistic understanding of how SI microbiota impacts the host.

## Methods

### Mice

All animal husbandry and experiments were approved by the Caltech Institutional Animal Care and Use Committee (IACUC, protocol #1646). The EE mouse model was established following the previously published protocol [20] with slight adaptations. On day 1 of the experiment, just-weaned, specific pathogen free (SPF) C57BL/6J male mice were received from the Jackson Laboratory (Sacramento, CA, USA) at 21 days of age. Because the purpose of this work was to answer whether bacterial colonization of jejunum mucosa was plausible, we reasoned that gender comparison was not necessary at this stage and focused on a single gender for consistency. However, future studies into the phenomenon of bacterial colonization of jejunum mucosa would require gender comparison. When mice were received for more than one experimental condition, they were randomized upon arrival across those conditions. For the duration of the study, the mice were housed in sterile cages with 4–5 mice per cage. Mice had ad libitum access to one of the experimental diets, either complete (COM) or malnourishing (MAL) (Table S3). The malnourishing diet was deficient in fat and protein, but contained the same amounts of calories, vitamins, and minerals on a per-weight basis [20]. Diet formulations were consistent with the previous study [20] except that food dyes were omitted to reduce autofluorescence during imaging.

### Bacterial culture for gavage

Bacterial isolates of *Bacteroides dorei, Bacteroides ovatus, Bacteroides vulgatus, Parabacteroides distasonis*, and the two isolates of *Escherichia coli* were received from Brett Finley (University of British Columbia), and *Bacteroides fragilis* was received from Emma Allen Vercoe (University of Guelph). All isolates were identical to the previously used isolates in the Brown et al. study [20]. We confirmed isolate identities by Sanger sequencing; obtained amplicon sequences aligned perfectly with a region of full 16S rRNA sequences of the corresponding gavaged bacterial isolates (full 16S rRNA sequences were provided by E. A. Vercoe). All bacterial isolates were cultured at 37 °C in an anerobic chamber (Coy Laboratory Products, Grass Lake, MI, USA). Single colonies were cultured on Brucella blood agar plates (89405-032; VWR, Randor, PA, USA), and liquid cultures were grown in brain heart infusion (BHI) medium (90003-040; VWR) supplemented with 5 μg/mL hematin porcine (H3281; Sigma-Aldrich, St. Louis, MO, USA), 1 μg/mL vitamin K1 (L10575; ThermoFisher Scientific, Waltham, MA, USA) and 250 μg/mL L-cysteine (44889; ThermoFisher Scientific); hereafter, this supplemented BHI medium is referred as BHI-S. To minimize the number of passages, which may result in loss of phenotype and increase the risk of cross-contamination, bacteria were plated from frozen one-time-use glycerol stocks up to a week prior to the gavage and cultured for 1–2 days. The night before bacterial gavage, the isolates were cultured for ~14 h from single colonies in 5 mL of BHI-S. On the day of gavage, stationary overnight cultures were reinoculated into fresh 5 mL of BHI-S. The inoculum volume was selected so that all bacterial isolates reached midexponential phase within 2–3 h (40–50 μL for *E. coli* and 250–400 μL for *Bacteroides/Parabacteroides* spp. isolates). Bacterial growth was monitored by measuring optical density (OD) directly in transparent culture tubes (60819-524; VWR), with mid-exponential phase corresponding to OD ~ 1 (in the range 0.7–1.2). Exponential-phase cultures were pelleted, and the pellets were resuspended in PBS. Bacterial density in PBS suspensions was estimated using our established OD - CFU (colony forming units) correlations. Briefly, we established these correlations by assuming a linear relationship between OD and CFU for OD values below 0.5, which required two data points. For the first data point, we assumed that OD of 0 corresponded to 0 CFUs. For the second data point, we prepared bacterial suspensions in PBS with OD < 0.5 and plated these suspensions on Brucellar blood agar plates to enumerate CFUs. After estimation of bacterial density in PBS suspensions, bacterial cocktails were prepared in 2.5% sodium bicarbonate (25080094; ThermoFisher Scientific) in PBS at 1.43 · 10^8^ cells/mL density of each clinical isolate, with net bacterial density in *E. coli* + *Bacteroides*/*Parabacteroides* spp. (EC+BAC) cocktail adding up to 10^9^ cells/mL, in *E. coli* only cocktail (EC) – to 2.86· 10^8^ cells/mL, and in *Bacteroides/Parabacteroides* spp. – to 7.14· 10^8^ cells/mL. Bacterial cocktails were stored on ice and administered to mice within 1 h of preparation.

### Diet and gavage treatments

Six mouse groups were considered in this study: COM+PBS, COM+EC&BAC, MAL+PBS, MAL+BAC, MAL+EC, and MAL+EC&BAC. The mice were placed on one of the experimental diets (COM or MAL) starting day 1 of each experiment. Oral gavages took place the third week of each experiment. On days 14, 16, and 18, experimental animals were gavaged with 100 μL of one of the bacterial cocktails described above or the PBS control. All gavages occurred between 1-3 p.m. Malnourished mice were gavaged with one of the four possible gavages (EC&BAC, EC, BAC, or the PBS control), whereas controls received either the complete cocktail (EC&BAC) or the PBS control.

### 16S rRNA gene copy analysis

Jejunum lumenal contents (digesta) and feces were collected from all six mouse groups during the fifth week of the experiment, on days 28, 29, and 30, with one mouse per group per day and three mice per group (Fig. 2, A-E, and Fig. S1). The only exception was the MAL+PBS group: in this group, one mouse was euthanized on day 30 and the other two mice euthanized on day 31. However, because these mice were not exposed to bacteria, we did not anticipate that microbiota composition would change substantially over the course of a few days. The six mouse groups were staggered across two experiments, with the groups on the COM diet analyzed 1 month after the groups on the MAL diet (but on the same relative days of the experiment). Therefore, differences in microbiota composition between COM and MAL groups may be partially due to the differences in resident microbiota composition between the two experiments. Before euthanasia, each mouse was weighed in a clean plastic container, and fresh fecal pellets were collected from the weighing container. Each mouse was then euthanized by intraperitoneal (IP) euthasol injection (~250 μL of 10-fold euthasol (07-805-9296; Patterson Veterinary Supply, Devens, MA, USA) dilution in saline per mouse) and cardiac puncture. Jejunum digesta was collected from a full segment identified in the 2^nd^ quartile of the SI. Samples were stored on ice during sample collection and were moved to −80 °C within 1 h of collection.

For tail-cup intervention (Fig. 2F-G), mice were fitted with either functional or mock tail cups as previously described [38], on day 13 of the experiment and maintained on tail-cups until the end of the experiment. They were gavaged with EC&BAC cocktail on days 14, 16, and 18 of the experiment and euthanized on days 27-31, with one mouse per group per day. Fecal and jejunum digesta samples were collected following the same procedure as described above.

DNA from feces and jejunum digesta was extracted using a PowerSoil Pro kit (47016; Qiagen, Hilden, Germany) following manufacturer’s instructions with specifications [38, 40, 41]. Total bacterial 16S rRNA gene copy load was quantified by droplet digital PCR (dPCR) on a QX200 droplet generator (1864002; Bio-Rad Laboratories, Hercules, CA, USA) following our previously published protocols [40]. *Enterobacteriaceae* 16S rRNA gene copy load was also quantified by dPCR using *Enterobacteriaceae-specific* primers [64] following the same protocol except that annealing temperature was increased from 52 °C to 60 °C. Detailed summary of DNA extraction and dPCR results are provided in Table S2. 16S rRNA gene amplicon sequencing, including library amplification, sequencing, and data processing was performed following our previously published protocols [38, 40, 41]. Notably, the sequencing library was amplified following protocols that reduce amplification bias [38, 40, 41]. Detailed 16S rRNA gene amplicon sequencing results are provided in Table S1.

To calculate absolute taxon-specific 16S rRNA gene copy load (Fig. 2, C and D), the fractional taxon-specific abundance obtained by 16S rRNA gene amplicon sequencing (Fig. 2A) was first anchored by the total bacterial 16S rRNA gene copy load obtained by dPCR (Fig. 2B) for each sample (black circles), and then these absolute taxon-specific 16S gene copy loads were averaged for each treatment group (color bars). For 16S rRNA gene amplicon sequencing, the LOD was defined as detection of ≥3 reads per sample (Fig. 2, C and D, and Fig. S1, B and D); according to Poisson distribution, this corresponds to ≥95% probability of successful detection. Notably, according to this definition, a taxon can still be detected at fewer than 3 reads but less than 95% of the time. To translate this LOD definition to 16S rRNA gene copies per gram, the fractional LOD value (3 reads over total reads) was first anchored by the total 16S rRNA gene copy load obtained by dPCR for each sample, and then these values were averaged for each treatment group (Fig. 2, C and D, and Fig. S1, B and D, dashed lines). For dPCR with *Enterobacteriaceae* primers, the LOD was defined as *μ_background_* + 3*σ_background_*, where *μ_background_* and *σ_background_* are the mean and standard deviation of the background, respectively (Fig. 2, E-F, and Fig. S1, A and C). These parameters were calculated across all samples from mice not exposed to *E. coli* (COM+PBS, MAL+PBS, and MAL+BAC), yielding *μ_background_*, *σ_background_*, and LOD equal to 0.9, 1.1, and 4.2 copies/μL in dPCR reaction, respectively (Table S2). Thus, if the target is detected below this LOD value, it cannot be distinguished from background noise. The starting sample mass, DNA extraction recovery, and DNA extract dilution into the dPCR reaction were then used to translate this LOD definition to 16S rRNA gene copies per gram for each sample (Fig. 2, E-F, Fig. S1, A and C, and Table S2).

### RT-qPCR analysis of host gene expression

Jejunum (2^nd^ quarter of the SI), ileum (4^th^ quarter of the SI) and proximal colon tissue scrapings were collected from the four MAL groups (MAL+PBS, MAL+BAC, MAL+EC, and MAL+EC&ABC) on days 28, 29, 30, and 31, with one mouse per group per day and four mice per group. Tissues for RT-qPCR and imaging were collected from the same mice that were fasted for 1 h (see below). The tissue scrapings were collected in this order: jejunum (2^nd^ quartile of the SI), ileum (4^th^ quartile of the SI), and proximal colon; this order was chosen because, in our experience and as previously reported [65], mRNA is least stable in jejunum tissue, followed by ileum tissue and proximal colon tissue. Tissue scrapings were transferred immediately to 600 μL of 1x DNA/RNA Shield (R1100; Zymo, Irvine, CA, USA) in a Lysing Matrix D tube containing 1.4 mm ceramic spheres (116913050-CF; MP Biomedicals, Santa Ana, CA, USA). During the sample-collection procedure, the samples were stored and handled on ice. At the end of the procedure, they were homogenized by bead beating (6.5 m/s for 60 s on FastPrep24 (MP Biomedicals)) and stored at −80 °C.

After defrosting on ice, homogenized tissue samples preserved in DNA/RNA shield were immediately extracted using a Quick RNA Miniprep Plus kit (R1057; Zymo) per manufacturer’s instructions. Prior to RNA extractions, a proteinase K (D3001-2-20; Zymo) digestion step was included (60 μL of Proteinase K digestion buffer and 30 μL of Proteinase K were added to 600 μL of the homogenate, and the digestion was carried out for 30 min at 55 °C). During RNA purification, an on-column DNase treatment was performed per manufacturer’s instructions. Purified RNA was eluted into 50 μL of nuclease free water (AM9935; ThermoFisher Scientific). A 1 μL aliquot of sample was diluted 1:20 for RNA quantification and quality analysis, prior to storing samples at −80 °C. RNA quantification was performed using Qubit HS RNA kit (Q32856; ThermoFisher Scientific) and RNA quality was assessed using Agilent 2200 TapeStation system with high sensitivity RNA screen tape and sample buffer reagents (5067; Agilent, Santa Clara, CA, USA). cDNA was created using a high-capacity reverse transcription kit (4368814; ThermoFisher Scientific). We added 1.8 μg total RNA per 20 μl reaction. Reaction setup was performed on ice per manufacturer’s instructions. A Bio-Rad C1000 thermocycler was used to perform cDNA synthesis at 25 °C for 10 min, 37 °C for 120 min, 85 °C for 5 min, and 4 °C degree hold. Glyceraldehyde 3-phosphate dehydrogenase (GAPDH), tumor necrosis factor alpha (TNF-α), and interleukin 6 (IL-6) transcripts were quantified using commercially available TaqMan probes (Mm99999915_g1, Mm00443258_m1, and Mm00446190_m1, respectively; ThermoFisher Scientific) and TaqMan Fast Advanced MasterMix (4444556; ThermoFisher Scientific). Targets were detected with the FAM channel using Bio-Rad CFX96 qPCR instrument (thermocycler conditions: 50 °C for 2 min, 95 °C for 10 min, 40 cycles of 95 °C for 15 sec, 60 °C for 1 min). The technical replicates for each probe were averaged before normalization. TNF-alpha and IL-6 transcript levels were normalized to the house keeping gene (GAPDH) by subtracting housekeeping gene Cq from target gene Cq to obtain ΔCq [39]. Additionally, ΔCq values were normalized to the corresponding ΔCq values in the MAL+PBS reference group to obtain ΔΔCq values [39].

### Preparation of acrylamide monomer mix

Two acrylamide monomer mix chemistries were used to preserve each intestinal segment (see Table S4 for compositions). First, one of the surface-gel-monomer mix chemistries were used to polymerize a protective surface hydrogel layer to stabilize loosely adherent mucosal matter (e.g. mucus and bacteria). Second, one the tissue-gel-monomer mix chemistries were used to embed the tissues into an acrylamide hydrogel. More than 1 h before a monomer mix was needed, UltraPure water (10977023; ThermoFisher Scientific) and 10xPBS (46-013-CM; Corning) were combined in a 50 mL round bottom flask and chilled on ice; from this point, the monomer mix was stored continuously on ice. Approximately 30 min before the monomer mix was needed, the remaining chilled reagents were added. To remove oxygen, the mix was degassed with house vacuum for 20 min and purged with argon for 1 min. At this point, the monomer mix was brought to anaerobic chamber.

### Hydrogel tissue embedding of empty jejunum from non-fasted mice

During the fifth week of the experiment, empty jejunum segments from non-fasted mice – one mouse in each MAL treatment group (MAL+PBS, MAL+BAC, MAL+EC, and MAL+EC&BAC) — were embedded into an acrylamide hydrogel following our previously published protocols with modifications. Each mouse was anesthetized by IP euthasol injection and transcardially perfused with sterile-filtered ice-cold PBS supplemented with 10 U/mL heparin (H3149; Sigma-Aldrich) and 0.5 w/v% sodium nitrite (237213; Sigma-Aldrich) (hPBS) at a rate of 5 mL/min for 10 min. The gastrointestinal tract was excised and quick-fixed in an excess (40 mL) of ice-cold 4% PFA for 2–3 min. We observed that rapid fixation of the outer muscle layer reduces tissue warping during the subsequent preservation protocol. The gut was then rinsed in ice-cold PBS and further dissected on an ice-cold dissection stage in a biosafety cabinet (BSC). The gut was untangled, and the mesentery was removed. A 2–3 cm long empty jejunum segment was identified in mid-SI segment and excised. The segment was placed on a microscope slide that had a thin (1 mm thick) silicone isolator glued to it (666103; Grace Bio-labs, Bend, OR, USA). The segment was opened longitudinally using fine dissection scissors (504024; World Precision Instruments, Sarasota, FL, USA) and laid out flat on the slide with the mucosa facing up. Two additional thick (1.8 mm thick) silicone isolators (666203; Grace Bio-labs) were stacked on top of the bottom isolator to create a ~2.5 mL reservoir for subsequent incubations. The reservoir was first filled with 4% PFA, covered with another microscope slide to prevent PFA evaporation, and the slide was placed on ice for 1 h to fix the tissue.

After fixation, the tissues were flipped upside down so the mucosa faced the glass slide (we have observed that the most convenient way to flip the tissues is to drag them over a cover slip, flip the cover slip upside down, and drag the tissue back to the liquid-filled reservoir). Flipped tissue was moved to an anaerobic chamber where an oxygen-sensitive polymerization of acrylamide hydrogel was performed. In the anaerobic chamber, the isolator was filled with A4B.08P4 surface-gel-monomer mix (see Table S4 for compositions) and the tissue was incubated on ice for 15 min to allow the monomer mix to displace liquid in tissue crevices. Then most of the monomer mix was removed to leave only a fine layer under the mucosa – this fine layer ultimately polymerizes into a protective surface gel. The isolator was covered with a silicone sheet (664475; Grace Bio-labs) to allow for gas exchange, and the surface gel was polymerized at 37 °C for 3 h in a humid environment (humidity was maintained by placing either an open container with water in the 37 °C incubator or wet tissue directly in a petri dish with tissue samples). After surface gel polymerization, the tissue was stored at 4 °C and proceeded to hydrogel tissue embedding within 3 days.

Tissue embedding into a hydrogel was also performed in the anaerobic chamber. The reservoirs were filled with A4B0P4 tissue-gel-monomer mix and infused on ice for 3h. After infusion, the tissue-gel-monomer mix was removed, the reservoirs were covered with silicone sheets for gas exchange, and hydrogel-tissue hybrids were polymerized at 37 °C in a humid environment for 3 h. Finally, hydrogel-tissue hybrids were removed from the anaerobic chamber, the muscle side of the tissues was glued to a polypropylene solid support (46510; Crawford Industries, Crawfordsville, IN, USA) using GLUture tissue glue (10014489; Zoetis, Parsippany, NJ, USA), and the tissues were dislodged from the glass slide with a microtome blade (15148-236; VWR). The plastic was trimmed around the tissues, leaving short overhangs on both ends of intestinal segments (Fig. 4A) to facilitate handling and mounting for imaging. The tissues were stored in 0.025% sodium azide (BDH7465-2; VWR) in PBS until lysozyme permeabilization.

### Hydrogel tissue embedding of empty jejunum from fasted mice

Only the malnourished groups were considered in this study (MAL+PBS, MAL+BAC, MAL+EC, and MAL+EC&BAC), which were staggered across four experiments separated from each other by a week. In each group, the mice were analyzed on days 28, 29, 30, and 31, with one mouse per day. Tissue preservation for imaging is time consuming; therefore, only a few mice can be analyzed over the course of a single day. Considering that circadian rhythms can affect microbiota [43], we decided to perform processing over the course of 4 days, but consistently euthanize all mice at the same time each day (11 am). To link the images to any potential host response, the tissues for imaging and RTqPCR were collected from the same mice. After euthanasia, the gut was dissected and the segments of interest were excised: 2^nd^, 3^rd^, and 4^th^ quartile of the SI as well as proximal colon. 3^rd^ quartile of the SI was subjected to tissue preservation for imaging, whereas the rest of the tissues were collected for RTqPCR analysis of host gene expression (see above). At this point, the tissues for imaging and mRNA analysis were handled in parallel.

Empty jejunum segments from fasted mice were embedded into an acrylamide hydrogel following a similar procedure with modifications (see above). First, to synchronize the passage of digesta across the mice to the best of our ability, each mouse was fasted for 1 h before euthanasia and euthanized consistently at 11 am. Second, transcardial perfusion was omitted because it swells the gut [44], which can disrupt the spatial structure of mucosal bacteria and alter the concentration of RTqPCR targets. Third, to improve the permeability of hydrogel-tissue hybrids for antibody staining, the more permeable A4B.08P1 and A1B.01P4 surface and tissue-gel-monomer mixes were used, respectively. This time, the tissues were infused with tissue-gel-monomer mix immediately after surface gel polymerization on ice for 18 h and then polymerized at 37 °C for 5 h.

### Permeabilization of bacterial peptidoglycan with lysozyme

Lysozyme treatment was performed in 10 mM Tris-HCl, pH 8.0, buffer prepared from 1 M Tris-HCl, pH 8.0 (AM9856; ThermoFisher Scientific) and UltraPure water. Before lysozyme treatment, hydrogel-tissue hybrids were incubated in lysozyme treatment buffer at RT for 1 h. Bacterial peptidoglycan layer was then permeabilized with lysozyme (90082; ThermoFisher Scientific) at 37 °C for 6 h with gentle shaking; 5 mg/mL of lysozyme was used for A4B.08P4 / A4B0P4 hydrogel chemistry, and 1 mg/mL of lysozyme was used for more permeable A4B.08P1 / A1B.01P4 hydrogel chemistry. After treatment, the hydrogel-tissue hybrids were first rinsed in PBS and then washed three times in excess of PBS over the course of one day.

### SDS clearing

Clearing solutions were prepared from 10% SDS (51213; Lonza, Rockland, ME, USA), 10xPBS, and UltraPure water. A4B.08P4 / A4B0P4 hydrogel-tissue hybrids were cleared in 8% SDS, pH 8.3, at 37 °C for 3 days with daily pH adjustments (but without clearing solution changes). These hydrogel-tissue hybrids were placed into separate cassettes (22-272416; Fisher Scientific, Waltham, MA, USA) in a single large crystallizing dish, submerged in the clearing solution, and cleared with stirring on a stirring hot plate. After clearing, these hydrogel-tissue hybrids were washed in the setup in PBS at 25 °C for 3 days with daily wash solution changes. A4B.08P1 / A1B.01P4 hydrogel-tissue hybrids were cleared in 4% SDS, pH 8.5, at 37 °C for 3 days with daily solution changes. These hydrogel-tissue hybrids were placed into separate 50 mL tubes fully filled with the clearing solution and cleared with shaking in a shaking incubator. After clearing, these hydrogel-tissue hybrids were washed in the same setup, first twice in 0.1% TritonX-100 in PBS at 37 °C for 1 day and then twice in PBS at RT for another day.

### HCR v2.0 tagging of bacteria in hydrogel-tissue hybrids from non-fasted mice

HCR v2.0 probes were designed by concatenating B4, B5, and B2 initiator sequences (Table S5) to the 3’ end of EUB338, GAM42a, and CFB560ab probes (Table S5), respectively, and purchased from IDT. EUB338 and GAM42a probes have a higher melting temperature (Tm) than CFB560ab probes; therefore, probe hybridization was performed in two steps with two different formamide concentrations. Otherwise, each hybridization step was performed at 46 °C for 16 h in 5 mL of hybridization solution per two hydrogel-tissue hybrids (two hydrogel-tissue hybrids were grouped in a single 5 mL tube (490019-562; VWR)). In the first hybridization step, hydrogel-tissue hybrids were hybridized with EUB338-B4 (10 nM) and GAM42a-B5 (10 nM) probes in a hybridization buffer consisting of 15 v/v% formamide (BP227-100; Fisher Scientific), 10 w/v% high molecular weight dextran sulfate (D8906; Sigma-Aldrich), and 2xSSC (sodium saline citrate) buffer (2xSSC was prepared from 20xSSC (PAV4261, VWR) and UltraPure water). After the first hybridization, each hydrogel-tissue hybrid was rinsed in room-temperature 2xSSCT (0.1% Tween 20 (P1379; Sigma-Aldrich) in 2xSSC) and washed in 30% formamide in 2xSSCT followed by 2xSSCT, and finally PBS; each wash performed in 30 mL of wash buffer per sample with shaking at RT for 1 h. In the second hybridization step, hydrogel-tissue hybrids were hybridized with CFB560a-B2 (10 nM) and CFB560b-B2 (10 nM) probes in the hybridization buffer that now excluded formamide. After the second hybridization, hydrogel-tissue hybrids were washed similarly except that the duration of each wash was extended to 2 h. Probes were amplified in a single step with B2-Alexafluor647, B4-Alexafluor546, and B5-Alexafluor488 amplifier pairs (Molecular Technologies, Pasadena, CA, USA) at RT for 16 h. Prior to combining all amplifiers in the amplification solution, they were heat-shocked at 95 °C for 90 s and cooled to RT for 30 min. Amplification solution consisted of 0.12 μM of each amplifier in the amplification buffer consisting of 10 w/v% high molecular weight dextran sulfate and 2xSSC. Amplification was carried out in hybridization chambers (666203; Grace Bio-Labs) glued to HybriSlip covers (716024; Grace Bio-Labs) on both sides; ~1.5 mL of amplification solution was required to fill each chamber depending on the sample size. After amplification, each hydrogel-tissue hybrid was rinsed in room-temperature 5xSSCT (0.1% Tween 20 in 5xSSC prepared from 20xSSC and UltraPure water) and washed twice in 5xSSCT and once in PBS; each wash performed in 30 mL of wash buffer per sample with shaking at RT for 2 h. The samples were then stored in 5 μg/mL DAPI (D1306; Thermo Fisher Scientific) and 0.025% sodium in PBS, 5 mL per sample, until imaging.

### Design and validation of universal bacterial HCR v3.0 degenerate probe set

While HCR v2.0 probes recognize a region of approximately 20 base pairs (bp), split HCR v3.0 probes recognize a longer 52 bp region, complicating the design of broad coverage HCR v3.0 probes, such as the universal bacterial 16S rRNA probe. For example, 18-bp long EUB338 probe covers >95% of all known bacterial 16S rRNA sequences without a mismatch, however, no single 52 bp region is shared among >95% of sequences. Thus, we designed a universal bacterial HCR v3.0 degenerate probe set. First, a 52 bp region around the EUB338 probe binding site compatible with the HCR v3.0 reaction mechanism was selected by Molecular Technologies. We then aligned this region to the 16S rRNA sequences from SILVA 138 NR99 database, considering only the most prevalent bacterial orders in mouse and human gut. After alignment, we selected the most frequent hits that maximized coverage. The universal bacterial HCR v3.0 probe set was narrowed down to 17 52-bp-long probes (Table S5), however, different combinations of split probes further increased coverage. The exact location of the split is proprietary (Molecular Instruments Corp.); therefore, we assessed the coverage of the hypothetical 26 bp long probes (Fig. S7). For most bacterial orders under consideration, both split degenerate probe sets covered >80% 16S rRNA sequences in SILVA database without mismatch (Fig. S7). The probes are not expected to differentiate between sequences with zero or one mismatch; therefore, in practice, our designed degenerate universal HCR v3.0 probes are expected to cover nearly all bacteria in the bacterial orders of interest. Universal bacterial HCR v3.0 degenerate probe set was validated against EUB338 HCR v2.0 probe in a proximal colon from SPF mice (Fig. S8 and SI Methods). The signal from the universal bacterial HCR v3.0 degenerate probe set correlated with the signal from EUB338 HCR v2.0 probe (coefficient of correlation = 0.8), suggesting that the new probes successfully recognized bacteria (Fig. S8L). That coefficient of correlation was below 1 may be explained by the competition between the two probes for the same binding site, HCR v3.0 requirement for bound probes to initiate amplification, and the propensity of HCR v2.0 toward background amplification. Importantly, unlike HCR v2.0 (Fig. S8, A, E, and J), HCR v3.0 was resistant to background amplification (Fig. S8, B, E, and K).

### Design of taxon-specific HCR v3.0 probes

Two taxon-specific HCR v3.0 probes were designed by Molecular Technologies using full 16S rRNA sequences of the gavaged bacterial isolates, one probe specific to both *E. coli* isolates but orthogonal to *Bacteroides*/*Parabacteroides* spp. and one probe specific to all five *Bacteroides/Parabacteroides* spp. but orthogonal to *E. coli* (Table S5. The probes were synthesized by Molecular Technologies and provided as 1 μM stocks. The specificity and sensitivity of the new probes were determined in silico by aligning probe sequences to complete 16S rRNA sequences from SILVA 138 NR99 database and calculating % of 16S rRNA sequences in each bacterial taxon that aligned with the probe sequences when a given number of mismatches was allowed (Fig. S9A); only the sequences of the bacterial orders detected in experimental animals by sequencing were considered (Fig. 2A). The new *E. coli HCR v3.0* probe only recognized *Enterobacteriaceae* family, solely represented by *gavaged E. coli*, even when one mismatch was allowed. In contrast, the new *Bacteroides/Parabacteroides* spp. HCR v3.0 probe recognized *Bacteroidaceae* (the family of gavaged *Bacteroides* spp. and resident *B. theta*) and *Tannerellaceae* (the family of gavaged *P. distasonis*) families without a mismatch and recognized *Muribaculaceae* family with one mismatch (Fig. S9A); thus, the new *Bacteroides/Parabacteroides* spp. probe may have still cross-reacted with *Muribaculaceae* and was referred to as *Bacteroidales* probe hereafter. The designed probes were also validated *in vitro* using *B. fragilis* and one of the *E. coli* isolates (SI Methods): the probes recognized their targets but did not cross-react (Fig. S9B). Furthermore, although the new *Bacteroidales* HCR v3.0 probe overlapped with CFB560 HCR v2.0 probe binding site (Table S5), it resolved false-positive tagging of *E. coli* deep in a hydrogel associated with CFB560 HCR v2.0 probe (Fig. S10).

### HCR v3.0 tagging of bacteria in hydrogel-tissue hybrids from fasted mice

Before hybridization, hydrogel-tissue hybrids were incubated in 30% formamide in 5xSSCT, 5 mL per sample, at 37 °C for 2 h. They were then hybridized with the universal bacterial HCR v3.0 degenerate probe set (B1 initiator, 4 nM of each degenerate probe), *Bacteroides/Parabacteroides* spp. HCR v3.0 probe (B2 initiator, 20 nM), and *E. coli* HCR v3.0 probe (B3 initiator, 20 nM) in probe hybridization buffer for cells in suspension (Molecular Technologies) at 37 °C for 20 h. Hybridization was carried out in 5 mL tubes with 4 mL of hybridization solution and two hydrogel-tissue hybrids per tube (one tissue from each of two mice from the same mouse group). After hybridization, the hydrogel-tissue hybrids were rinsed in warm (equilibrated to 37 °C) probe wash buffer (Molecular Technologies) and then washed in the dilution series of probe wash buffer in 5xSSCT (3:0 for 30 min, 2:1 for 30 min, 1:2 for 60 min, 0:3 for 60 min, and again 0:3 for 60 min), with 20 mL of wash buffer per sample per wash. Hybridized probes were amplified with B1-Alexafluor514, B2-Alexafluor647, and B3-Alexafluor594 amplifier pairs (Molecular Technologies), with 0.12 μM of each amplifier in amplification buffer for cells in suspension (Molecular Technologies), at RT for 20 h. As previously, amplifiers were heat-shocked before combining them in the amplification solution. Amplification was carried out in 5 mL tubes with 3.75 mL of amplification solution and two hydrogel-tissue hybrids per tube (one tissue from each of two mice from the same mouse group). After amplification, hydrogel-tissue hybrids were rinsed in room-temperature 5xSSCT and then washed in 5xSSCT (twice for 30 min and twice for 60 min) and finally in PBS for 60 min, with 20 mL of wash buffer per sample per wash. All hybridization and amplification washes were performed at RT with shaking. The samples were then stored in 5 μg/mL DAPI and 0.025% sodium azide in PBS, 5 mL per sample, until imaging.

### Antibody and lectin staining of hydrogel-tissue hybrids from fasted mice

To remove refractive index matching solution (RIMS) after 20xCLARITY imaging of HCR-stained hydrogel-tissue hybrids from fasted mice, the hybrids were washed for 1 day in PBS, 50 mL per sample, at RT and with gentle shaking. Before antibody and lectin staining, the hybrids were washed for another day in 100 mM Tris-HCl, pH 8.0, 50 mL per sample, at RT and with gentle shaking; this step was included to scavenge free formaldehyde that may be generated by formaldehyde reverse crosslinking [66, 67] and may react with antibodies and lectins. Small pieces of hydrogel-tissue hybrids were then cut off for antibody and lectin staining. Each piece was first blocked overnight in 1 mL of blocking buffer (2% serum (100487-948; VWR) in PBST (0.1% TritonX-100 (ICN19485450; Fisher Scientific) and 0.1% sodium azide in PBS)) at RT and with gentle shaking. The next day, each piece was transferred to 0.5 mL of the staining solution prepared in the same blocking buffer and stained for 3 days at RT and with gentle shaking. Right before preparing staining solutions, antibody and lectin stocks – 1 mg/mL WGA-A488 (WGA conjugated to Alexafluor488 (W11261; ThermoFisher Scientific)), 0.2 mg/mL anti-EpCAM-A546 (anti-EpCAM antibody conjugated to Alexafluor546 (sc-53532 AF546; Santa Cruz Biotechnology, Santa Cruz, CA, USA)), and 0.2 mg/mL anti-CD45-A546 (anti-CD45 antibody conjugated to Alexafluor546 (sc-53665 AF546; Santa Cruz Biotechnology)) – were centrifuged at 10,000 g and 4 °C to remove any large aggregates. In Fig. 5, a piece of the hydrogel-tissue hybrid with bacterial clusters in the mucosa was stained with 2 μg/mL WGA-A488, 1 ug/mL anti-EpCAM-A546, and 5 ug/mL DAPI. In Fig. 6, one piece of the hydrogel-tissue hybrid with large surface aggregates was stained with 2 ug/mL WGA-A488, 5 ug/mL anti-EpCAM-A546, and 5 ug/mL DAPI, and another piece was stained with 2 ug/mL WGA-A488, 5 ug/mL anti-CD45-A546, and 5 ug/mL DAPI. After staining, each piece was washed three times in PBST, 30 mL per sample per wash, at RT and with gentle shaking over a course of one day.

### Fluorescence microscopy

All imaging was performed on a Zeiss LSM880 confocal microscope. All imaging, image display, and image segmentation and filtering metadata are provided in Table S6.

For large-scale low-magnification tile-scanning, hydrogel-tissue hybrids were mounted in PBS. Briefly, plastic overhangs were taped to a microscope slide with tape (2097-36EC; 3M, Saint Paul, MN, USA) and then silicone isolators (depending on sample thickness, either RD481862 or RD481863; Grace Bio-Labs) were glued to the slide with the hybrid positioned in the center. The isolator was filled with PBS and covered with a cover slip (16002-264; VWR). We observed that during the long acquisition of large-scale low-magnification tile scans, air bubbles can form under the cover slip (Fig. S17), possibly due to liquid evaporation and air diffusion across silicone isolator-cover slip contact. Therefore, to reduce air bubble formation, we applied oil along the perimeter of silicone isolator and cover slip contact. Mounted hydrogel-tissue hybrids were tile-scanned with a 5x objective (EC Plan-Neofluar 5x/0.16 420330-9901; Zeiss, Oberkochen, Germany) at 10% overlap between the tiles. The tile scans were acquired in channel mode with two channels. In the first channel, 405 nm laser was used to visualize DAPI staining of the epithelium; in the second channel, 561 nm (for hydrogel-tissue hybrids from non-fasted mice, Fig. 3B) or 514 nm (for hydrogel-tissue hybrids from fasted mice, Fig. 4B and Figs. S12-S27) lasers were used to visualize HCR staining of total bacteria. After the acquisition, the tile scans were stitched in ZEN software (Zen 2.3 SP1, Zeiss).

For imaging at high magnification, hydrogel-tissue hybrids were mounted in RIMS [31] (600 mL of 0.02 M phosphate buffer (P5244; Sigma-Aldrich) + 800 g of Histodenz (CAS #66108-95-0; JINLAN Pharm-Drugs Technology Co., Hangzhou, China) supplemented with 0.01% sodium azide, pH=7.5, refractive index (RI)=1.465). Briefly, plastic overhangs were taped to the bottom of the deep petri dish (89107-632; VWR), submerged in a large volume of RIMS (50-60 mL), and incubated overnight at RT with gentle shaking. To prevent RIMS dehydration and changes to its refractive index, the petri dish was wrapped with parafilm during infusion and RIMS was covered with a with a layer of Immersion Oil Type FF (100496-526; VWR) during imaging. Hydrogel-tissue hybrids mounted in RIMS were imaged with 20x CLARITY objective (Clr Plan-Neofluar 20x/1.0 Corr nd=1.45 M32, RI=1.464; Zeiss).

HCR tagging of tagging of total bacteria and individual taxa was imaged in channel mode with 4 channels total. For hydrogel-tissue hybrids from non-fasted mice, 561, 488, and 633 nm lasers were used to visualize HCR v2.0 staining of total bacteria, *Gammaproteobacteria*, and *Bacteroidetes*, respectively (Fig. 3C and Figs. S2-S6). For hydrogel-tissue hybrids from fasted mice, 514, 561, and 633 nm lasers were used to visualize HCR v3.0 staining of total bacteria, *E. coli*,and *Bacteroidales*, respectively (Fig. 4C, Fig. 5, A, B and D, and Figs. S28-S35). In both experiments, 405 nm laser was used to visualize DAPI staining of epithelium. For each hydrogel-tissue hybrid, three images were acquired, with each image was composed of 4 (2×2) fields of view imaged at 10% overlap.

HCR tagging of total bacteria and WGA staining of mucus (and other N-acetylglucosamine and sialic acid-rich glycoprotein) (Fig. 5C) were imaged in spectral mode to mitigate potential bleed-through between abundant bacteria and mucus in the mucosa of the hydrogel-tissue hybrid. Four lasers were used to excite the fluorophores (405, 514, 561, and 633 nm), and emitted light was collected in 8.9 nm bins. Linear unmixing was performed in ZEN using fluorescence spectra for each fluorophore or tissue autofluorescence control. Briefly, to generate fluorescence spectra, hydrogel-tissue hybrids were prepared, stained, mounted, and imaged following identical protocols except that each piece of tissue was stained with a single fluorophore only (staining was omitted for the tissue autofluorescence control), generating a spectral file for each fluorophore (or tissue autofluorescence control). In each of these files, a region of interest (ROI) was selected in ZEN that was characterized by a strong but not saturated signal. For the tissue autofluorescence control, ROI was selected in the tissue. Spectral profiles in these ROIs were then used to unmix spectral imaging files in ZEN. Each image was composed of a single field of view.

Antibody staining of either epithelial or immune cells and WGA staining of mucus (and other N-acetylglucosamine and sialic acid-rich glycoprotein) (Fig. 6E) were imaged in channel mode. This sample was also previously stained by HCR v3.0 for bacteria, which may have produced bleed-through into antibody and lectin channels. However, because in this sample antibody and lectin staining was much stronger or more prevalent than HCR v3.0 staining, we did not observe significant bleed-through for visual inspection purposes. 405, 488, and 561 nm lasers were used to visualize DAPI staining of epithelium, WGA staining of mucus, and antibody staining of epithelial or total immune cells, respectively. For each antibody, a single image was acquired composed of 4 (2×2) fields of view imaged at 10% overlap.

### Image analysis

The images were segmented and filtered in Imaris software (Imaris 9.7, Oxford Instruments, Abington, United Kingdom). Identical thresholds were applied to images acquired under the same imaging settings (Table S6).

To segment fluorescence images with HCR tagging of bacteria and DAPI staining of host nuclei, iso-surfaces were created by setting the minimum fluorescence intensity threshold for each channel, yielding four categories of surfaces: DAPI, total and two taxon-specific bacteria. Various parameters were calculated for the segmented surfaces: center-of-mass position, volume, mean fluorescence intensity in each channel, as well as shortest surface-to-surface distance and overlapped volume ratio to the nearest surface in other surface categories. Imaris does not calculate the shortest surface-to-surface distance to the nearest surface in the same surface category. Therefore, to calculate this distance, we used center-of-mass position to first calculate the shortest center-to-center distance and then we corrected it to take into account the volume of surfaces.

Segmented surfaces were then filtered to remove false and double positives. Several criteria were used to remove false positives. First, candidate total and taxon-specific bacterial surfaces were filtered by volume to remove small surfaces characteristic of background noise. Specifically, under our imaging settings, the voxel size was 0.83 μm × 0.83 μm × 0.83 μm, or 0.57 μm^3^; therefore, a 1 μm wide spherical bacterial cell was expected to occupy 2×2×2 voxels and have an apparent volume of 8 × 0.57 μm^3^= 4.57 μm^3^ (we set minimum size threshold to 4 μm^3^). Second, candidate total and taxon-specific bacterial surfaces were filtered by their distance to DAPI+ surfaces to remove surfaces inside DAPI+ surface (that is, distance to DAPI+ surfaces <0 μm). Based on visual inspection, candidate bacterial surface inside the tissue corresponded to tissue or blood autofluorescence, especially in mice that were not transcardially perfused. We acknowledge that removing candidate bacterial surfaces inside DAPI+ surfaces removes translocated bacteria, if any; however, translocation was not the focus of this study. Finally, candidate taxon-specific bacterial surfaces were filtered based on the evidence for total bacterial staining, removing surfaces dim in total bacterial channel and distant from total bacterial surfaces. Notably, we allowed taxon-specific bacterial surfaces to be up to 0.5 μm away from total bacterial surfaces because, in multicolor imaging, the position of an object along the z-coordinate differs for each color wavelength due to the chromatic aberrations of lenses so setting arbitrary segmentation thresholds can separate surfaces that physically overlap.

A single criterion was applied to remove double-positive taxon-specific bacterial surfaces. Specifically, for each taxon, we removed those surfaces that overlapped substantially with the surfaces of the other taxon. We allowed up to 0.4-0.5 overlap ratio because, with 0.83 μm × 0.83 μm × 0.83 μm voxel size, individual bacterial cells in dense bacterial clusters cannot be resolved and that there is significant bleed-through between taxon-specific bacterial channels.

The remaining surfaces were then analyzed to quantify bacterial composition and spatial distribution. In Fig. 3F, ECDF was calculated by first sorting GAM42a+ surfaces based on their shortest distance to another surface and then calculating the fraction of GAM42a+ surfaces (y-axis variable) less than a certain distance away from the target surface (x-axis variable). For example, 50% of GAM42a+ surfaces were less than 19 μm away from another GAM42a+ surface but in contact with CFB560+ surfaces. In Fig. 6E, CDF was calculated by first sorting DAPI+ surfaces by size (volume) and then counting the number of these DAPI+ surfaces (y-axis variable) less than a certain volume in size (x-axis variable).

### Statistical analyses

Considering the low number of replicates (4 biological replicates in Fig. 1, C-D and 3 biological replicates in Fig. 2, A-D, and Fig. S1, A-B), we could not assume governing statistical distributions. Therefore, as previously described [40], we used a non-parametric Kruskal-Wallis rank sums test implemented with *scipy.stats.kruskal* function. To account for multiple comparisons, *P*-values were adjusted with *statsmodels.stats.multitest.multipletests* function with the Benjamini–Hochberg correction.

## Declarations

### Ethics approval and consent to participate

All animal husbandry and experiments were approved by the Caltech Institutional Animal Care and Use Committee (IACUC, protocol #1646).

### Availability of data and material

All data from this publication are available at Caltech DATA at http://dx.doi.org/10.22002/D1.20199.

### Authors’ contributions

RP Conception, animal study execution, sample collection, sample processing for imaging, imaging, data analysis, figure generation, manuscript preparation. SRB Tail cup study implementation. sample collection. AER RT-qPCR, dPCR, and sequencing data acquisition. AHD dPCR data acquisition. OMP Preliminary imaging and sequencing. HT Preliminary image analysis. RFI Project supervision and administration, acquisition of funding, manuscript review and editing. All authors read and approved the final manuscript. See Supplementary Information for detailed author contributions

## Acknowledgements

We thank the Caltech Office of Laboratory Animal Resources as well as the veterinary technicians at the Wanimal facilities for animal care, personnel training, and resources. We thank Biological Imaging Facility at Caltech (including Andres Collazo, Giada Spigolon, and Steven Wilbert) for resources, training, and technical support. We thank Brett Finley (University of British Columbia) and Prof. Emma Allen Vercoe (University of Guelph) for providing bacterial isolates. We thank Prof. Jared Leadbetter and Prof. Sarkis Mazmanian for providing feedback on study design. We thank Justin Bois for introduction to data analysis in Python. We thank Emily Savela, Mary Arrastia and Eugenia Khorosheva for reviewing and filing Institutional Biosafety Committee paperwork. We thank Jacob T. Barlow for processing sequencing data. We thank Joanne Lau for maintaining anaerobic chambers for bacterial culture. We also thank Natasha Shelby for contributions to writing and editing this manuscript.

## References

1. Christl SU, Scheppach W: Metabolic consequences of total colectomy. Scandinavian Journal of Gastroenterology 1997, 222:20–24.

2. Spencer AU, Neaga A, West B, Safran J, Brown P, Btaiche I, Kuzma-O’Reilly B, Teitelbaum DH, Anderson KD, Teitelbaum DH, et al. Pediatric short bowel syndrome: Redefining predictors of success. Annals of Surgery 2005, 242:403–412.

3. Tang Q, Jin G, Wang G, Liu T, Liu X, Wang B, Cao H: Current Sampling Methods for Gut Microbiota: A Call for More Precise Devices. Frontiers in Cellular and Infection Microbiology 2020, 10:1–10.

4. Helander HF, Fändriks L: Surface area of the digestive tract - revisited. Scandinavian Journal of Gastroenterology 2014, 49:681–689.

5. Mowat AM, Agace WW: Regional specialization within the intestinal immune system. Nature Reviews Immunology 2014, 14:667–685.

6. El Aidy S, van den Bogert B, Kleerebezem M: The small intestine microbiota, nutritional modulation and relevance for health. Current Opinion in Biotechnology 2015, 32:14–20.

7. Bohm M, Siwiec RM, Wo JM: Diagnosis and management of small intestinal bacterial overgrowth. Nutrition in clinical practice 2013, 28:289–299.

8. Donaldson GP, Lee SM, Mazmanian SK: Gut biogeography of the bacterial microbiota. Nature Reviews Microbiology 2015, 14:20–32.

9. Lazarevic V, Gaïa N, Girard M, Schrenzel J: Decontamination of 16S rRNA gene amplicon sequence datasets based on bacterial load assessment by qPCR. BMC Microbiology 2016, 16:1–8.

10. Claassen-Weitz S, Gardner-lubbe S, Mwaikono KS, Toit E, Zar HJ, Nicol MP: Optimizing 16S rRNA gene profile analysis from low biomass nasopharyngeal and induced sputum specimens. BMC Microbiology 2020, 20:1–26.

11. Atuma C, Strugala V, Allen A, Holm L: The adherent gastrointestinal mucus gel layer: thickness and physical state in vivo. American Journal of Physiology-Gastrointestinal and Liver Physiology 2001, 280:G922–G929.

12. Johansson MEV, Sjövall H, Hansson GC: The gastrointestinal mucus system in health and disease. Nature reviews Gastroenterology & hepatology 2013, 10:352–361.

13. Bevins CL, Salzman NH: Paneth cells, antimicrobial peptides and maintenance of intestinal homeostasis. Nature Reviews Microbiology 2011, 9:356–368.

14. Pabst O, Slack E: IgA and the intestinal microbiota: the importance of being specific. Mucosal Immunology 2020, 13:12–21.

15. He G, Shankar RA, Chzhan M, Samouilov A, Kuppusamy P, Zweier JL: Noninvasive measurement of anatomic structure and intraluminal oxygenation in the gastrointestinal tract of living mice with spatial and spectral EPR imaging. Proceedings of the National Academy of Sciences of the United States of America 1999, 96:4586–4591.

16. Norkina O, Burnett T, Lisle RD: Bacterial overgrowth in the cystic fibrosis transmembrane conductance regulator null mouse small intestine. Infection and immunity 2004, 72:6040–6049.

17. Clarke LL, Gawenis LR, Bradford EM, Judd LM, Boyle KT, Simpson JE, Shull GE, Tanabe H, Ouellette AJ, Franklin CL, Walker NM: Abnormal Paneth cell granule dissolution and compromised resistance to bacterial colonization in the intestine of CF mice. American Journal of Physiology-Gastrointestinal and Liver Physiology 2004, 286:G1050–G1058.

18. Sovran B, Hugenholtz F, Elderman M, Van Beek AA, Graversen K, Huijskes M, Boekschoten MV, Savelkoul HFJ, De Vos P, Dekker J, Wells JM: Age-associated Impairment of the Mucus Barrier Function is Associated with Profound Changes in Microbiota and Immunity. Scientific Reports 2019, 9:1–13.

19. Tomas J, Mulet C, Saffarian A, Cavin J-B, Ducroc R, Regnault B, Kun Tan C, Duszka K, Burcelin R, Wahli W, et al: High-fat diet modifies the PPAR-γ pathway leading to disruption of microbial and physiological ecosystem in murine small intestine. Proceedings of the National Academy of Sciences 2016:201612559–201612559.

20. Brown EM, Wlodarska M, Willing BP, Vonaesch P, Han J, Reynolds LA, Arrieta M-C, Uhrig M, Scholz R, Partida O, et al: Diet and specific microbial exposure trigger features of environmental enteropathy in a novel murine model. Nature Communications 2015, 6:1–16.

21. Khazaei T, Williams RL, Bogatyrev SR, Doyle JC, Henry CS, Ismagilov RF: Metabolic multistability and hysteresis in a model aerobe-anaerobe microbiome community. Science Advances 2020, 6:1–10.

22. Guerrant RL, Oria RB, Moore SR, Oria MO, Lima AA: Malnutrition as an enteric infectious disease with longterm effects on child development. Nutr Rev 2008, 66:487–505.

23. Humphrey JH: Child undernutrition, tropical enteropathy, toilets, and handwashing. The Lancet 2009, 374:1032–1035.

24. Korpe PS, Petri WA: Environmental enteropathy: Critical implications of a poorly understood condition. Trends in Molecular Medicine 2012, 18:328–336.

25. Lim YF, De Loubens C, Love RJ, Lentle RG, Janssen PWM: Flow and mixing by small intestine villi. Food and Function 2015, 6:1787–1795.

26. Richardson DS, Lichtman JW: Clarifying Tissue Clearing. Cell 2015, 162:246–257.

27. Choi HMT, Chang JY, Trinh La, Padilla JE, Fraser SE, Pierce Na: Programmable in situ amplification for multiplexed imaging of mRNA expression. Nature biotechnology 2010, 28:1208–1212.

28. Shah S, Lubeck E, Schwarzkopf M, He T-f, Greenbaum A, Sohn Ch, Lignell A, Choi HMT, Gradinaru V, Pierce NA, Cai L: Single-molecule RNA detection at depth via hybridization chain reaction and tissue hydrogel embedding and clearing. Development 2016, 143:2862–2867.

29. Depas WH, Starwalt-lee R, Sambeek LV, Kumar R, Gradinaru V, Newman K: Exposing the Three-Dimensional Biogeography and Metabolic States of Pathogens in Cystic Fibrosis Sputum via Hydrogel Embedding, Clearing, and rRNA Labeling. mBio 2016, 7:1–11.

30. Choi HMT, Schwarzkopf M, Fornace ME, Acharya A, Artavanis G, Stegmaier J, Cunha A, Pierce NA: Third-generation in situ hybridization chain reaction: Multiplexed, quantitative, sensitive, versatile, robust. Development 2018, 145:1–10.

31. Yang B, Treweek JB, Kulkarni RP, Deverman BE, Chen CK, Lubeck E, Shah S, Cai L, Gradinaru V: Single-cell phenotyping within transparent intact tissue through whole-body clearing. Cell 2014, 158:945–958.

32. Neckel PH, Mattheus U, Hirt B, Just L, Mack AF: Large-scale tissue clearing (PACT): Technical evaluation and new perspectives in immunofluorescence, histology, and ultrastructure. Scientific Reports 2016, 6:1–13.

33. Wang W, Zhang N, Du Y, Gao J, Li M, Lin L, Czajkowsky DM, Li X, Yang CJ, Shao Z: Three-dimensional Quantitative Imaging of Native Microbiota Distribution in the Gut. Angewandte Chemie International Edition 2021, 60:3055–3061.

34. Mondragón-Palomino O, Poceviciute R, Lignell A, Griffiths JA, Takko H, Ismagilov RF: Three-dimensional imaging for the quantification of spatial patterns in microbiota of the intestinal mucosa. Proceedings of the National Academy of Sciences of the United States of America 2022, 119:1–12.

35. Barlow JT, Leite G, Romano AE, Sedighi R, Chang C, Celly S, Rezaie A, Mathur R, Pimentel M, Ismagilov RF: Quantitative sequencing clarifies the role of disruptor taxa, oral microbiota, and strict anaerobes in the human small-intestine microbiome. Microbiome 2021, 9:1–17.

36. Cervantes J, Michael M, Hong BY, Springer A, Guo H, Mendoza B, Zeng M, Sundin O, McCallum R: Investigation of oral, gastric, and duodenal microbiota in patients with upper gastrointestinal symptoms. Journal of Investigative Medicine 2021, 69:870–877.

37. Miller LS, Vegesna AK, Sampath AM, Prabhu S, Kotapati SK, Makipour K: Ileocecal valve dysfunction in small intestinal bacterial overgrowth: A pilot study. World Journal of Gastroenterology 2012, 18:6801–6808.

38. Bogatyrev SR, Rolando JC, Ismagilov RF: Self-reinoculation with fecal flora changes microbiota density and composition leading to an altered bile-acid profile in the mouse small intestine. Microbiome 2020, 8:19.

39. Livak KJ, Schmittgen TD: Analysis of relative gene expression data using real-time quantitative PCR and the 2-ΔΔCT method. Methods 2001, 25:402–408.

40. Barlow JT, Bogatyrev SR, Ismagilov RF: A quantitative sequencing framework for absolute abundance measurements of mucosal and lumenal microbial communities. Nature Communications 2020, 11.

41. Bogatyrev SR, Ismagilov RF: Quantitative microbiome profiling in lumenal and tissue samples with broad coverage and dynamic range via a single-step 16S rRNA gene DNA copy quantification and amplicon barcoding. bioRxiv 2020.

42. Johnson JS, Spakowicz DJ, Hong BY, Petersen LM, Demkowicz P, Chen L, Leopold SR, Hanson BM, Agresta HO, Gerstein M, et al: Evaluation of 16S rRNA gene sequencing for species and strain-level microbiome analysis. Nature Communications 2019, 10:1–11.

43. Leone V, Gibbons SM, Martinez K, Hutchison AL, Huang EY, Cham CM, Pierre JF, Heneghan AF, Nadimpalli A, Hubert N, et al: Effects of Diurnal Variation of Gut Microbes and High-Fat Feeding on Host Circadian Clock Function and Metabolism. Cell Host & Microbe 2015, 17:681–689.

44. Treweek JB, Chan KY, Flytzanis NC, Yang B, Deverman BE, Greenbaum A, Lignell A, Xiao C, Cai L, Ladinsky MS, et al: Whole-body tissue stabilization and selective extractions via tissue-hydrogel hybrids for high-resolution intact circuit mapping and phenotyping. Nature Protocols 2015, 10:1860–1896.

45. Huang JY, Lee SM, Mazmanian SK: The human commensal Bacteroides fragilis binds intestinal mucin. Anaerobe 2011, 17:137–141.

46. Bjursell MK, Martens EC, Gordon JI: Functional genomic and metabolic studies of the adaptations of a prominent adult human gut symbiont, Bacteroides thetaiotaomicron, to the suckling period. Journal of Biological Chemistry 2006, 281:36269–36279.

47. Huang YL, Chassard C, Hausmann M, Von Itzstein M, Hennet T: Sialic acid catabolism drives intestinal inflammation and microbial dysbiosis in mice. Nature Communications 2015, 6:1–11.

48. Johansson MEV, Holmén Larsson JM, Hansson GC: The two mucus layers of colon are organized by the MUC2 mucin, whereas the outer layer is a legislator of host-microbial interactions. Proceedings of the National Academy of Sciences of the United States of America 2011, 108:4659–4665.

49. Holmén Larsson JM, Thomsson KA, Rodríguez-Piñeiro AM, Karlsson H, Hansson GC: Studies of mucus in mouse stomach, small intestine, and colon. III. Gastrointestinal Muc5ac and Muc2 mucin O-glycan patterns reveal a regiospecific distribution. American journal of physiology Gastrointestinal and liver physiology 2013, 305:G357–363.

50. Molloy MJ, Grainger JR, Bouladoux N, Hand TW, Koo LY, Naik S, Quinones M, Dzutsev AK, Gao JL, Trinchieri G, et al: Intraluminal containment of commensal outgrowth in the gut during infection-induced dysbiosis. Cell Host and Microbe 2013, 14:318–328.

51. MacKenzie DA, Jeffers F, Parker ML, Vibert-Vallet A, Bongaerts RJ, Roos S, Walter J, Juge N: Strain-specific diversity of mucus-binding proteins in the adhesion and aggregation properties of Lactobacillus reuteri. Microbiology 2010, 156:3368–3378.

52. Singh VP, Proctor SD, Willing BP: Koch’s postulates, microbial dysbiosis and inflammatory bowel disease. Clinical Microbiology and Infection 2016, 22:594–599.

53. Samuel BS, Hansen EE, Manchester JK, Coutinho PM, Henrissat B, Fulton R, Latreille P, Kim K, Wilson RK, Gordon JI: Genomic and metabolic adaptations of Methanobrevibacter smithii to the human gut. Proceedings of the National Academy of Sciences of the United States of America 2007, 104:10643–10648.

54. Dejea CM, Fathi P, Craig JM, Boleij A, Taddese R, Geis AL, Wu X, DeStefano Shields CE, Hechenbleikner EM, Huso DL, et al: Patients with familial adenomatous polyposis harbor colonic biofilms containing tumorigenic bacteria. Science 2018, 359:592–597.

55. Rotstein OD, Vittorini T, Kao J, McBurney MI, Nasmith PE, Grinstein S: A soluble Bacteroides by-product impairs phagocytic killing of Escherichia coli by neutrophils. Infection and Immunity 1989, 57:745–753.

56. Vatanen T, Kostic AD, D’Hennezel E, Siljander H, Franzosa EA, Yassour M, Kolde R, Vlamakis H, Arthur TD, Hämäläinen AM, et al: Variation in Microbiome LPS Immunogenicity Contributes to Autoimmunity in Humans. Cell 2016, 165:842–853.

57. Bouhnik Y, Sophie AA, Flourié B: Bacterial populations contaminating the upper gut in patients with small intestinal bacterial overgrowth syndrome. American Journal of Gastroenterology 1999, 94:1327–1331.

58. Sundin OH, Mendoza-Ladd A, Zeng M, Diaz-Arévalo D, Morales E, Fagan BM, Ordoñez J, Velez P, Antony N, McCallum RW: The human jejunum has an endogenous microbiota that differs from those in the oral cavity and colon. BMC Microbiology 2017, 17:1–17.

59. Chen RY, Kung VL, Das S, Hossain MS, Hibberd MC, Guruge J, Mahfuz M, Begum SMKN, Rahman MM, Fahim SM, et al: Duodenal Microbiota in Stunted Undernourished Children with Enteropathy. New England Journal of Medicine 2020, 383:321–333.

60. Swidsinski A, Weber J, Loening-Baucke V, Hale LP, Lochs H: Spatial organization and composition of the mucosal flora in patients with inflammatory bowel disease. Journal of Clinical Microbiology 2005, 43:3380–3389.

61. Antonioli DA: Celiac disease: A progress report. Modern Pathology 2003, 16:342–346.

62. Donowitz JR, Haque R, Kirkpatrick BD, Alam M, Lu M, Kabir M, Kakon SH, Islam BZ, Afreen S, Musa A, et al: Small intestine bacterial overgrowth and environmental enteropathy in Bangladeshi children. mBio 2016, 7:1–7.

63. Ghoshal UC, Srivastava D: Irritable bowel syndrome and small intestinal bacterial overgrowth: Meaningful association or unnecessary hype. World Journal of Gastroenterology 2014, 20:2482–2491.

64. Castillo M, Martín-Orúe SM, Manzanilla EG, Badiola I, Martín M, Gasa J: Quantification of total bacteria, enterobacteria and lactobacilli populations in pig digesta by real-time PCR. Veterinary Microbiology 2006, 114:165–170.

65. Fleige S, Pfaffl MW: RNA integrity and the effect on the real-time qRT-PCR performance. Molecular Aspects of Medicine 2006, 27:126–139.

66. Kennedy-Darling J, Smith LM: Measuring the formaldehyde protein-DNA cross-link reversal rate. Analytical Chemistry 2014, 86:5678–5681.

67. Hoffman EA, Frey BL, Smith LM, Auble DT: Formaldehyde crosslinking: A tool for the study of chromatin complexes. Journal of Biological Chemistry 2015, 290:26404–26411.

