## Supplementary Materials for "Quantitative whole-tissue 3D imaging reveals bacteria in close association with mouse jejunum mucosa"

**Contents**

- Supplementary Materials and Methods
    - *Validation of the universal degenerate HCR v3.0 probe set*
    - *Validation of the taxon-specific HCR v3.0 probes*
    - *Comparison of the taxon-specific HCR v2.0 and HCR v3.0 probe sets*
    - *Comparison of A1B.01P4 and A4B0P4 tissue gel formulations*
- Supplementary Figures S1 – S35
- Supplementary Video Caption
- Supplementary Tables S1 – S6 (see also .xlsx attachments)
- Supplementary References
- Author Contributions

### Supplementary Materials and Methods

*Validation of the universal degenerate HCR v3.0 probe set*

The universal degenerate HCR v3.0 probe set was validated on a proximal colon segment from a CHOW-fed C57BL/6J mouse (Fig. S8). The segment was preserved following the protocol for empty jejunum segments from fasted mice (see Methods). The resulting hydrogel-tissue hybrid was permeabilized with lysozyme and cleared with SDS as described in the main text except that clearing duration was extended to 5 days because proximal colon tissue is thicker than jejunum tissue. The cleared hydrogel-tissue hybrid was stained for total bacteria following HCR v3.0 protocol with modifications. Specifically, the hydrogel-tissue hybrid was hybridized with EUB338 HCR v2.0 probe linked to B4 initiator and the universal degenerate HCR v3.0 probe set linked to B2 initiator in 3 mL of probe hybridization buffer for tissues in whole-mount (Molecular Technologies). The concentration of each universal degenerate HCR v3.0 probe was 4 nM; considering that each arm of the split universal HCR v3.0 probe set contains ~10 probes (6 and 12 to be exact), we set EUB338 concentration to 40 nM to avoid out-competition. Hybridized probes were amplified with B4-AlexaFluor546 and B2-AlexaFluor647 amplifier pairs in 1.5 mL of amplification buffer for tissues in whole-mount (Molecular Technologies). After HCR, the hydrogel-tissue hybrid was stained overnight at room temperature (RT) with 5 µg/mL DAPI and 5 µg/mL WGA-AlexaFluor488 in 5 mL of PBS. After overnight infusion in RIMS, the sample was imaged with the 20x CLARITY objective. All imaging and display metadata are provided in Table S6.

*Validation of taxon-specific HCR v3.0 probes*

Taxon-specific HCR v3.0 probes were validated *in vitro* on two isolates in the bacterial cocktail: *E. coli* and *B. fragilis* (Fig. S9). *E. coli* and *B. fragilis* suspensions in PBS were prepared as for bacterial gavage described in the main text. These suspensions were then mixed with 8% PFA in 1:1 ratio and fixed on ice for 1 h. Fixed bacterial cells were spiked into A4B.08P1 surface gel monomer mix at ~$5 \cdot 10^7$ cells/mL density and polymerized into a hydrogel slab; hydrogel embedding protocol for empty jejunum segments from fasted mice was otherwise followed. The resulting hydrogel slabs were permeabilized with lysozyme and cleared with SDS as described in the main text except that clearing duration was reduced to 1 d. Processed *E. coli* and *B. fragilis* gel slabs were stained following the HCR v3.0 protocol (see Methods) except that the slabs were hybridized with *E. coli* and *Bacteroidales* HCR v3.0 probes linked to B5 and B4 initiators, respectively, each at 10 nM concentration in 1 mL of probe hybridization buffer for tissues in whole-mount, and then amplified with B5-AlexaFluor488 and B4-AlexaFluor546 amplifiers in 0.5 mL of amplification buffer for tissues in whole-mount. After HCR, the gel slabs were stained overnight with 1 µM TO-PRO-3 Iodide DNA stain (T3605; ThermoFisher Scientific) in PBS and then imaged with 20x water immersion objective. Imaging and display settings are provided in Table S6.

*Comparison of taxon-specific HCR v2.0 and HCR v3.0 probe sets*

Taxon-specific HCR v2.0 and HCR v3.0 probes have been compared *in vitro* on one of the *E. coli* isolates only (Fig. S10). *E. coli* gel slab was prepared and processed as described above. *E. coli* gel slab was then stained following HCR v3.0 protocol except that it was hybridized with CFB560b-B5 HCR v2.0 and *Bacteroidales*-B3 HCR v3.0 probes, each at 10 nM in 1 mL of probe hybridization buffer for tissues in whole-mount, and then amplified with B5-AlexaFluor488 and B3-AlexaFluor594 amplifiers in 0.5 mL of amplification buffer for tissues in whole-mount. After HCR tagging of bacteria, the hydrogels were counterstained with 5 µg/mL of DAPI overnight and imaged with 20x water immersion objective. Imaging and display settings are provided in Table S6.

*Comparison of A1B.01P4 and A4B0P4 tissue gel formulations*

The effect of tissue gel formulation on antibody staining was compared on jejunum segments from a CHOW-fed C57BL/6J mouse (Fig. S11). Two segments were preserved following the protocol for empty jejunum segments from non-fasted mice with modifications. Specifically, transcardial perfusion as well as PFA in the surface gel monomer mix were omitted, and one segment was embedded into A4B0P4 tissue gel with 3 h polymerization duration. The other segment was embedded into A1B.01P4 tissue gel with 5 h polymerization duration (see Table S4 for monomer mix compositions). The hydrogel-tissue hybrids were then permeabilized with lysozyme and cleared with SDS as described in the main text except that clearing duration was extended to 5 d. Clearing progress was monitored daily by measuring hydrogel-tissue hybrid absorbance with a portable spectrophotometer (80-2116-30; Harvard Bioscience, Holliston, MA, USA) (Fig. S11D). After clearing, net protein loss was quantified with the microBCA Protein Assay Kit (23235, ThermoFisher Scientific) following manufacturer's instructions (Fig. S11C). Processed hydrogel-tissue hybrids were stained with 5 µg/mL anti-CD45 antibody and 5 µg/mL DAPI as described in the main text except that staining duration was reduced to 24 h. Finally, stained samples were mounted in RIMS and imaged with 20x CLARITY objective (imaging metadata is provided in Table S6). One z-stack was acquired for each condition, for a total of 2 z-stacks.

Antibody penetration (Fig. S11B) was quantified in FIJI. Briefly, a region of interest (ROI) was first drawn to mark the tip of a villus so that the z-distance from the villus tip could later be quantified. Then, at various planes along the villus, four ROIs were selected: three marking immune cells in the core of the villus (signal) and one marking epithelial cells (background). Average ROI fluorescence was calculated in FIJI, and the results were exported for further analysis in Python. In Python, the signal-background ratio was calculated as the ratio of average fluorescence in signal ROI to average fluorescence in background ROI at the same z-plane. Each data point in Fig. S11B represents signal-background ratio averaged over three signal ROIs at the same plane.

### Supplementary Figures

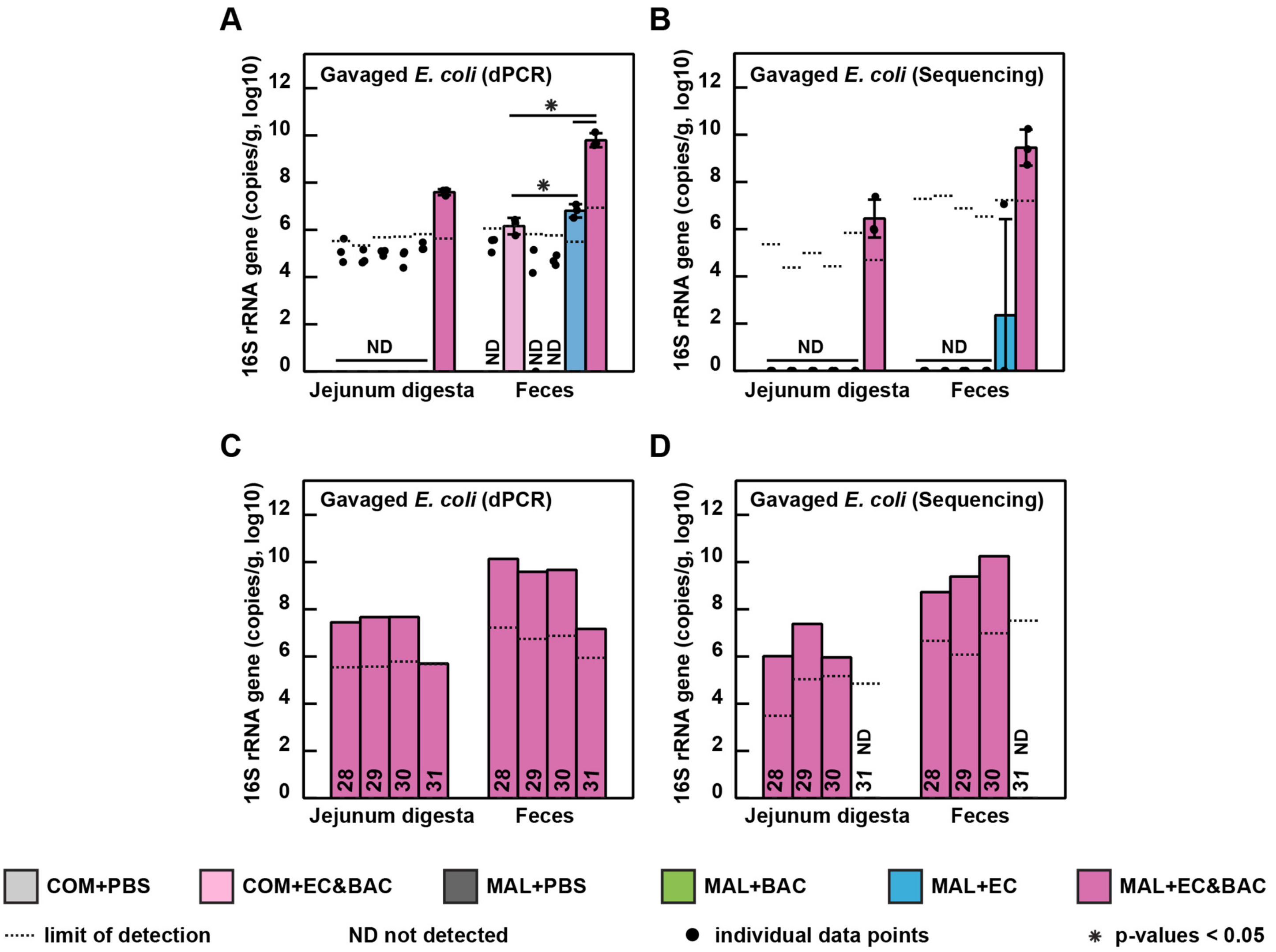


**Figure S1. Absolute abundance of *E. coli* 16S rRNA gene copies in jejunum digesta and feces across treatment groups.** (A) Log-absolute 16S rRNA gene copy abundance of gavaged *E. coli* quantified by digital PCR (dPCR) with *Enterobacteriaceae* primers. (B) Extrapolation of log-absolute 16S rRNA gene copy abundance of gavaged *E. coli* using 16S rRNA gene amplicon sequencing data (Fig. 2A) and total 16S rRNA gene copy quantification by dPCR with universal bacterial primers (Fig. 2B) [1]. Panel B is identical to Fig. 2D and displayed here for comparison purposes. In panels A and B, the analysis consisted of six groups: control mice not gavaged with bacteria (COM+PBS), control mice gavaged with an *E. coli* and *Bacteroides*/*Parabacteroides* spp. cocktail (COM+EC&BAC), malnourished mice not gavaged with bacteria (MAL+PBS), malnourished mice gavaged only with *Bacteroides*/*Parabacteroides* spp. isolates (MAL+BAC), malnourished mice gavaged only with *E. coli* isolates (MAL+EC), and malnourished mice gavaged with the full bacterial cocktail (MAL+EC&BAC). Each group contained three mice, which were euthanized on days 28, 29, and 30, with one mouse per group per day. Limit of detection is expressed as group average. Statistical significance was evaluated using Kruskal-Wallis tests. (C) Log-absolute 16S rRNA gene copy abundance of gavaged *E. coli* quantified by dPCR with *Enterobacteriaceae* primers in MAL+EC&BAC group. (D) Extrapolation of log-absolute 16S rRNA gene copy abundance of gavaged *E. coli* in MAL+EC&BAC group using 16S rRNA gene amplicon sequencing data (Fig. 2A) and total 16S rRNA gene copy quantification by dPCR with universal bacterial primers (Fig. 2B). In C and D, each bar represents an individual mouse euthanized on the specified day of the experiment.

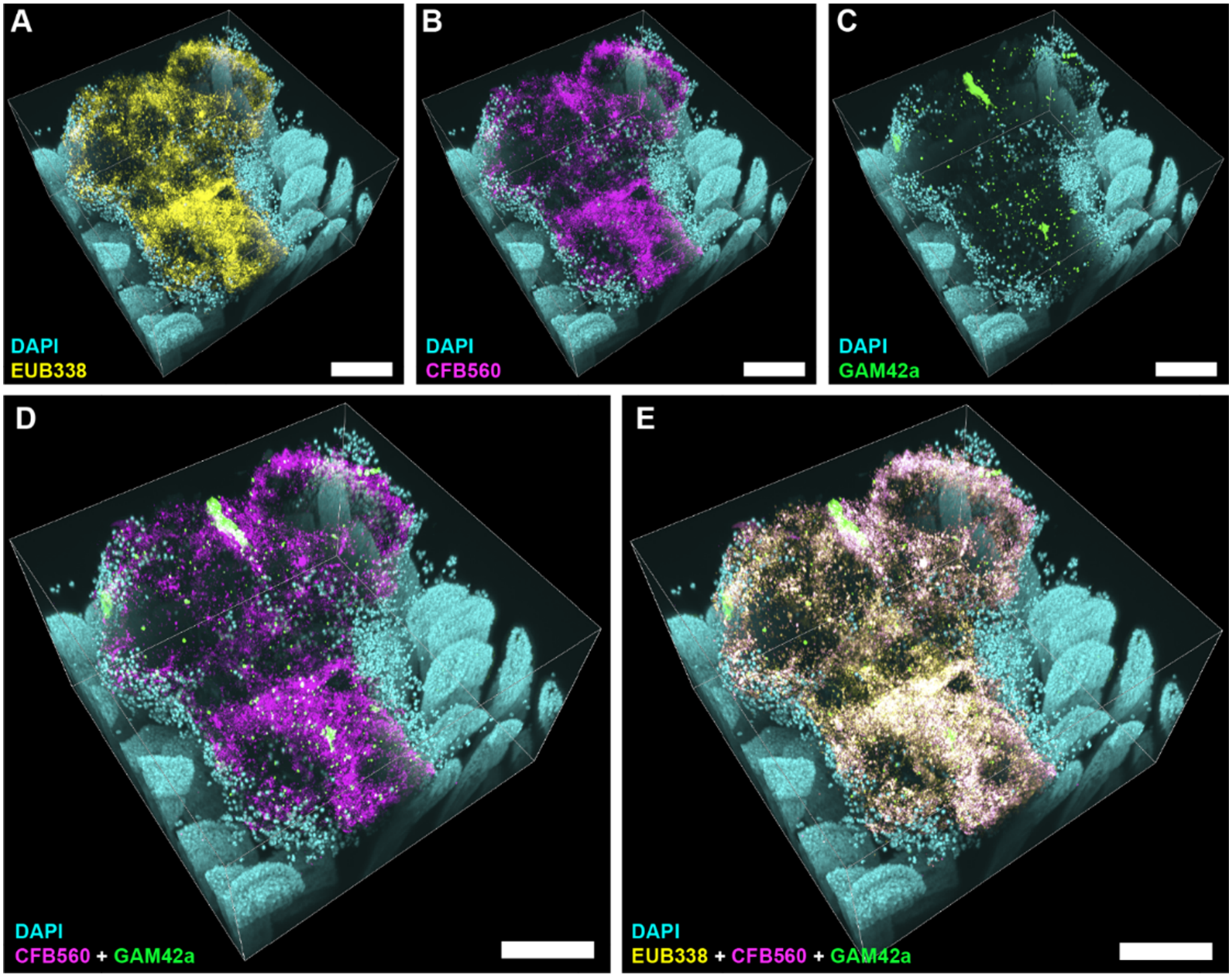


**Figure S2. High-magnification 3D imaging of a large surface aggregate detected in a MAL+EC&BAC mouse.** (A-E) High-magnification 3D fluorescence image of a large surface aggregate seen in Fig. 3B (red square) showing DAPI staining of epithelium (cyan) and HCR v2.0 staining of (A) total bacteria with EUB338 probe (yellow), (B) *Bacteroidetes* with CFB560 probe (magenta), (C) *Gammaproteobacteria* with GAM42a probe (green), (D) *Bacteroidetes* and *Gammaproteobacteria*, and (E) total bacteria, *Bacteroidetes*, and *Gammaproteobacteria*. Panel E shows the same large surface aggregate as shown Fig. 3C but displayed here at an enlarged scale. All scale bars 200 µm.

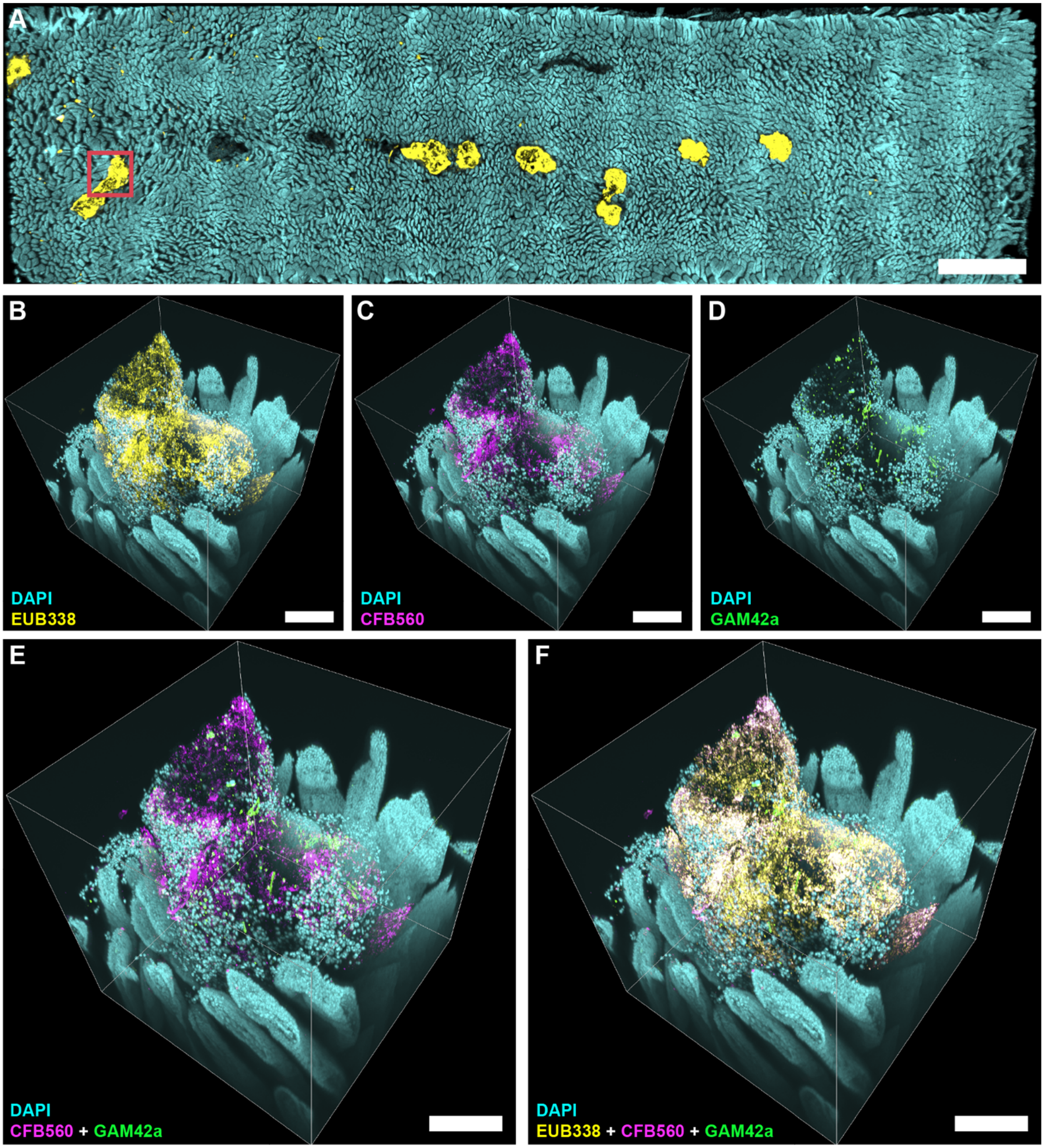


**Figure S3. High-magnification 3D imaging of a large surface aggregate detected in a MAL+EC&BAC mouse.** (A) Large-scale low-magnification fluorescence tile scans of a whole empty jejunum segment showing DAPI staining of epithelial surface (cyan) and HCR v2.0 staining of total bacteria with EUB338 probe (yellow). Red square marks the position where high magnification 3D image shown in panels B-F was acquired. Scale bar 2 mm. (B-F) High-magnification 3D fluorescence image of a large surface aggregate DAPI staining of epithelium (cyan) and HCR v2.0 staining of (A) total bacteria with EUB338 probe (yellow), (B) *Bacteroidetes* with CFB560 probe (magenta), (C) *Gammaproteobacteria* with GAM42a probe (green), (D) *Bacteroidetes* and *Gammaproteobacteria*, and (E) total bacteria, *Bacteroidetes*, and *Gammaproteobacteria*. This is the second of two large surface aggregates imaged at higher magnification; the other is shown in Fig. 3C. All scale bars 200 µm.

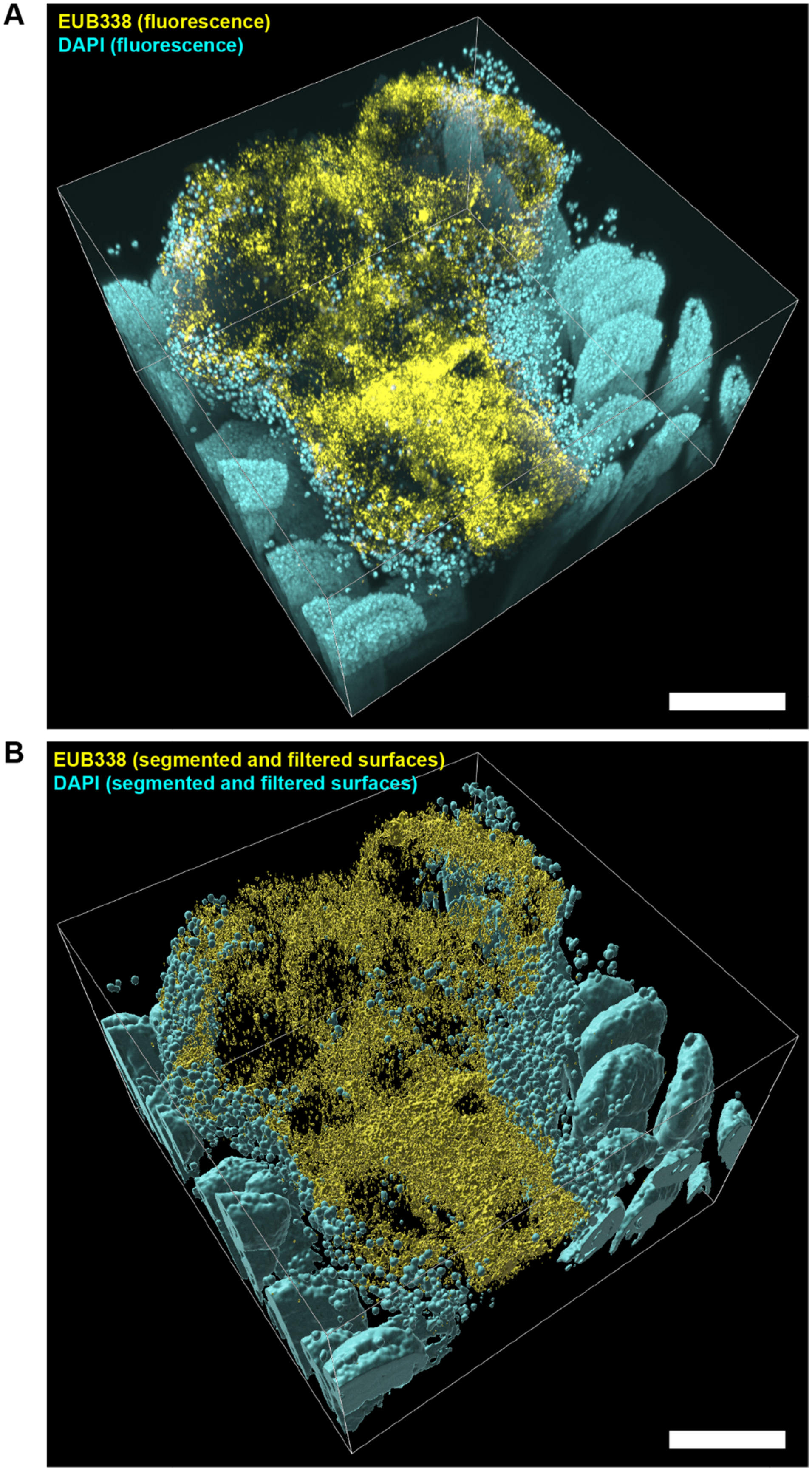


**Figure S4. Example image segmentation and filtering of total bacteria and host epithelium for a large surface aggregate detected in a MAL+EC&BAC mouse.** (A) High-magnification 3D fluorescence image before segmentation and filtering of a large surface aggregate seen in Fig. 3B (red square) showing DAPI staining of epithelium (cyan) and HCR v2.0 staining of total bacteria with EUB338 probe (yellow). (B) 3D rendering of segmented and filtered image showing DAPI+ and EUB338+ surfaces. All scale bars 200 µm.

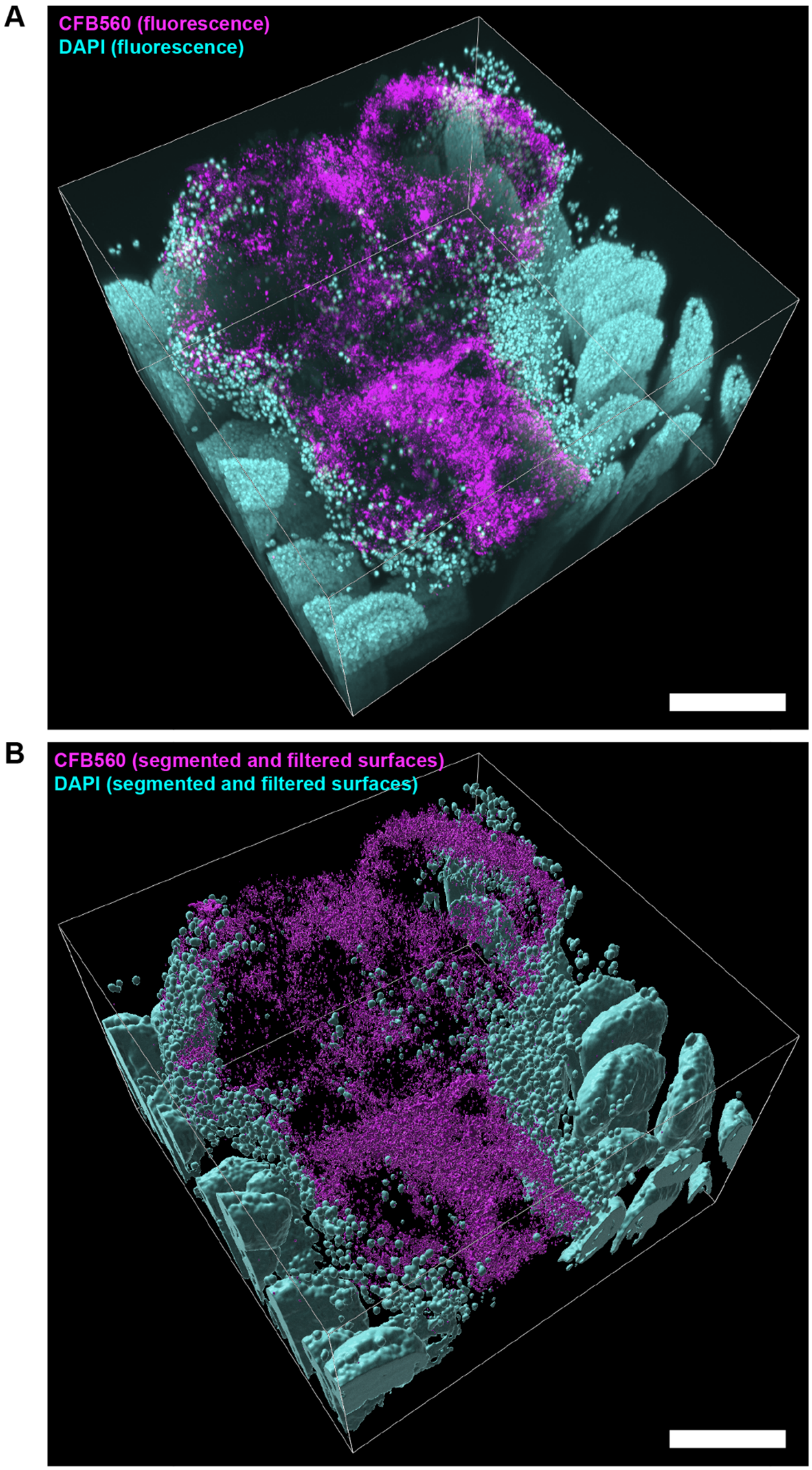


**Figure S5. Example image segmentation and filtering of *Bacteroidetes* and host epithelium for a large surface aggregate detected in a MAL+EC&BAC mouse.** (A) High-magnification 3D fluorescence image before segmentation and filtering of a large surface aggregate seen in Fig. 3B (red square) showing DAPI staining of epithelium (cyan) and HCR v2.0 staining of *Bacteroidetes* with CFB560 probe (magenta). (B) 3D rendering of segmented and filtered image showing DAPI+ and CFB560+ surfaces. All scale bars 200 µm.

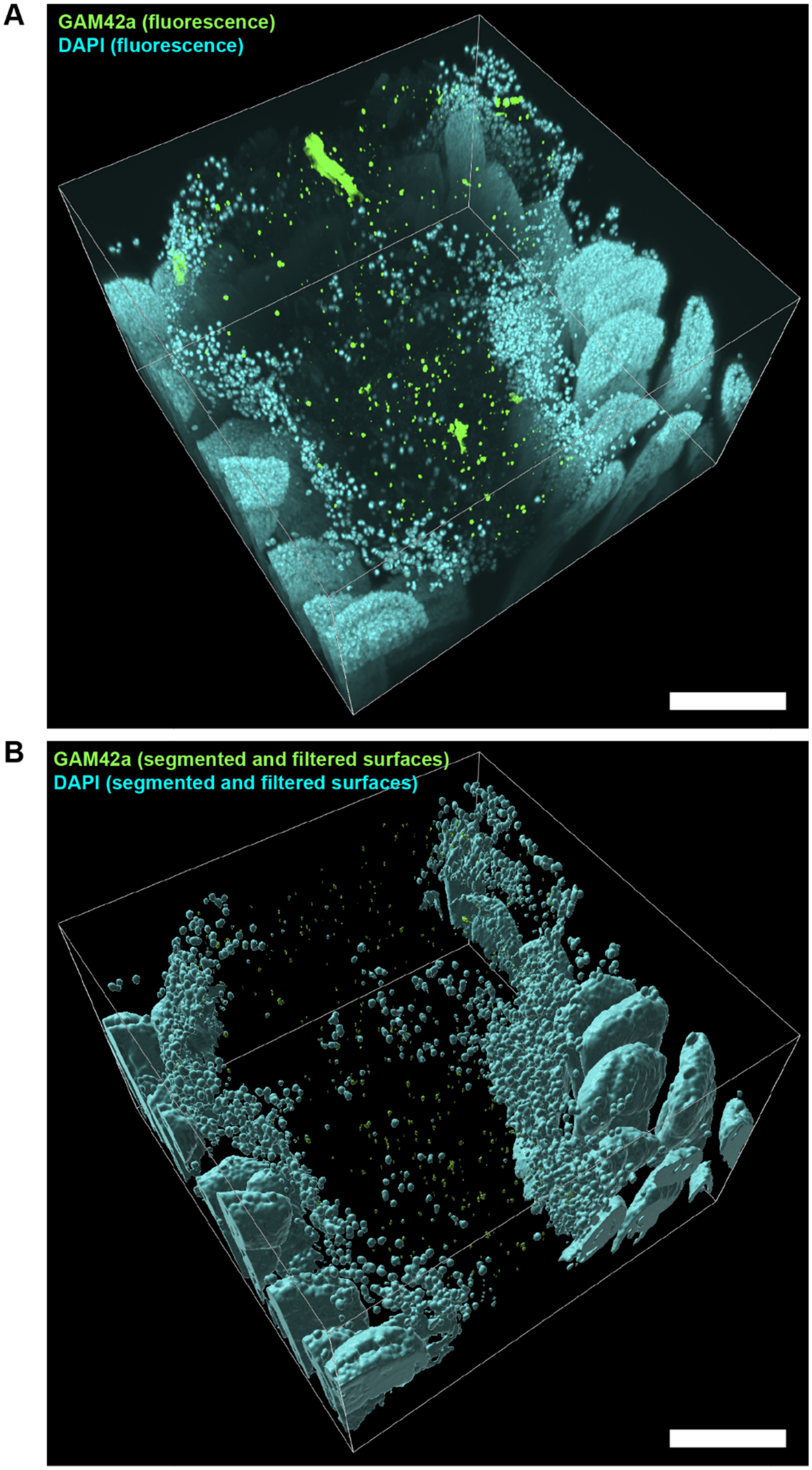


**Figure S6. Example image segmentation and filtering of *Gammaproteobacteria* and host epithelium for a large surface aggregate detected in a MAL+EC&BAC mouse.** (A) High-magnification 3D fluorescence image before segmentation and filtering of a large surface aggregate seen in Fig. 3B (red square) showing DAPI staining of epithelium (cyan) and HCR v2.0 staining of *Gammaproteobacteria* with GAM42a probe (magenta). (B) 3D rendering of segmented and filtered image showing DAPI+ and GAM42a+ surfaces. All scale bars 200 µm.

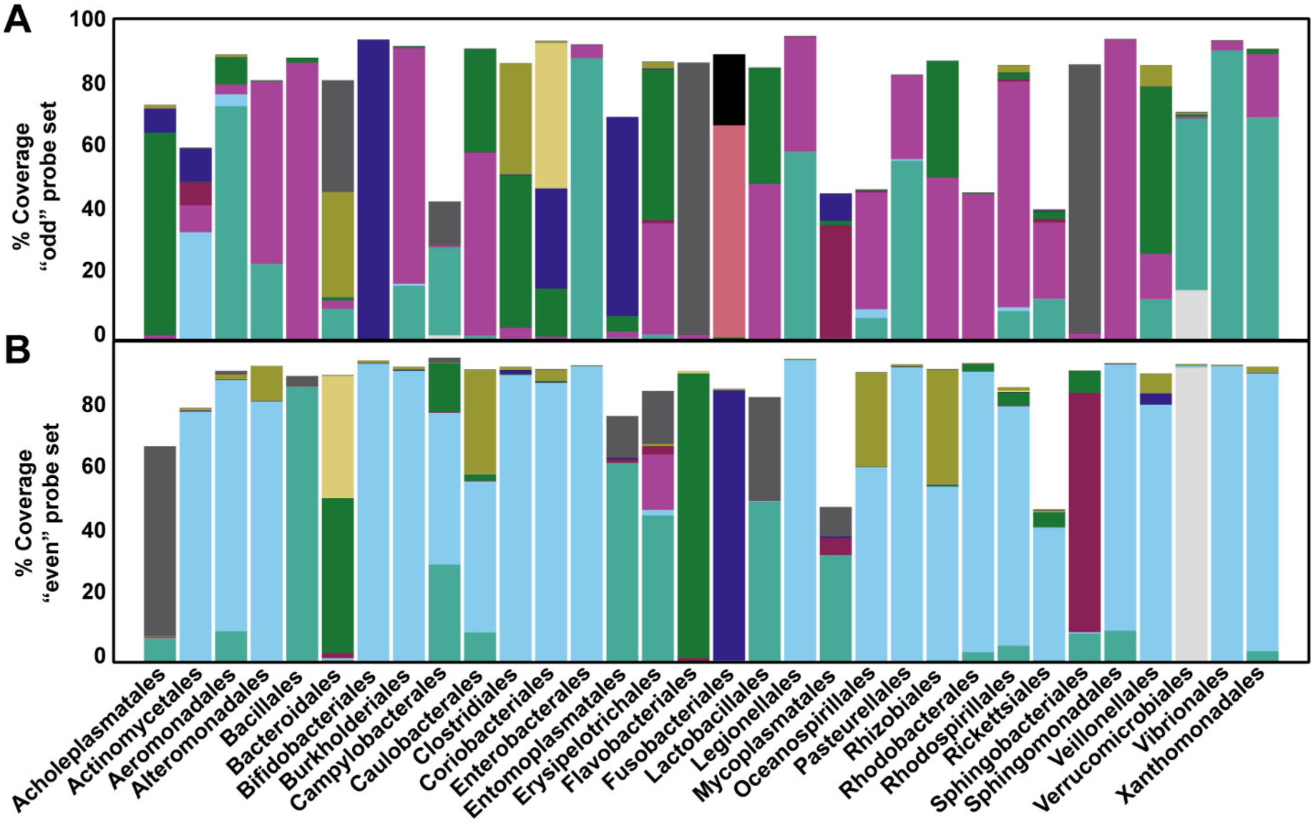


**Figure S7. Design of the universal degenerate HCR v3.0 probe set.** HCR v3.0 achieves suppression of off-target signal amplification by requiring two target-bound probes to initiate signal amplification, which decreases the likelihood that the non-specifically bound probes are amplified [2]. To design HCR v3.0 probes for total bacteria, a 52-bp region that overlaps with the EUB338 binding site was first selected by Molecular Technologies. To improve coverage, we expanded this region to 17 full degenerate probes, and analyzed the coverage of 10 and 12 split probes split in the center, referred here as “odd” (A) and “even” (B). The location of the split is proprietary (and it actually yields 6 and 12 probes); therefore, we analyzed the coverage of the hypothetical 26-bp-long split probes. The height of each colored bar represents the percentage of coverage without a mismatch by one degenerate split probe, and the net bar height represents cumulative coverage by the degenerate probe set of the bacterial order of interest. Coverage is defined as % of order-specific 16S rRNA sequences in the SILVA 138 NR99 database that align with the probe without a mismatch. Only common bacterial orders were considered in the analysis.

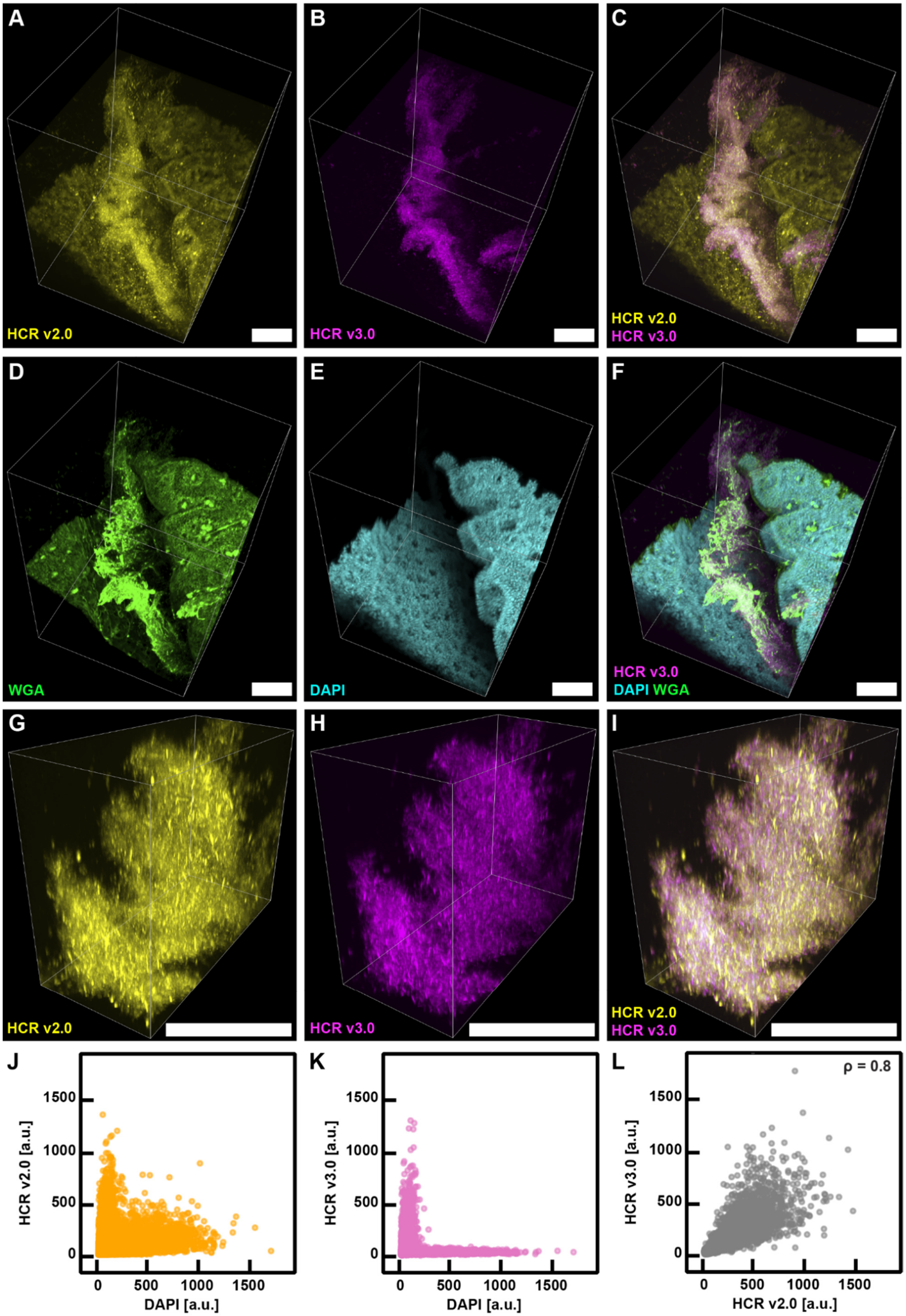


**Figure S8. Comparison of the universal degenerate HCR v3.0 probe set and EUB338 HCR v2.0 probe in a proximal colon of CHOW-fed control mouse.** (A-I) CLARITY imaging of a mouse proximal colon with the full acquired z-stack shown in (A-F) and the zoomed in portions focused on bacteria shown in (G-I). All scale bars 100 µm. (A) Bacteria tagging with EUB338 HCR v2.0 probe (yellow). (B) Bacteria tagging with the universal degenerate HCR v3.0 probe set (magenta). (C) Overlay of A and B showing that the signals overlapped in the center of the image, i.e. in between two proximal colon folds. (D) Epithelium staining with DAPI (cyan) showing proximal colon folds and a void in between. (E) Mucus staining with WGA lectin (green) showing mucus-filled goblet cells in the tissue and secreted mucus in between proximal colon folds. (F) Overlay of (B), (D), and (E) showing spatial structure of tissue, secreted mucus and bacteria tagged by HCR v3.0. (J-K) Voxel intensity analysis of a sub-sampled voxel population in panels (A-F). (J) HCR v2.0 signal vs DAPI signal, showing that voxels well-stained by DAPI also had elevated HCR v2.0 signal. (K) HCR v3.0 signal vs DAPI signal, showing that voxels well-stained by DAPI segregate from voxels well-stained by HCR v3.0. (L) Voxel intensity analysis of the zoomed in images shown in (G-I), showing that HCR v3.0 signal positively correlated with HCR v2.0 signal (Pearson's correlation coefficient = 0.8). These probes compete for the same binding site, partially explaining the <1 correlation coefficient.

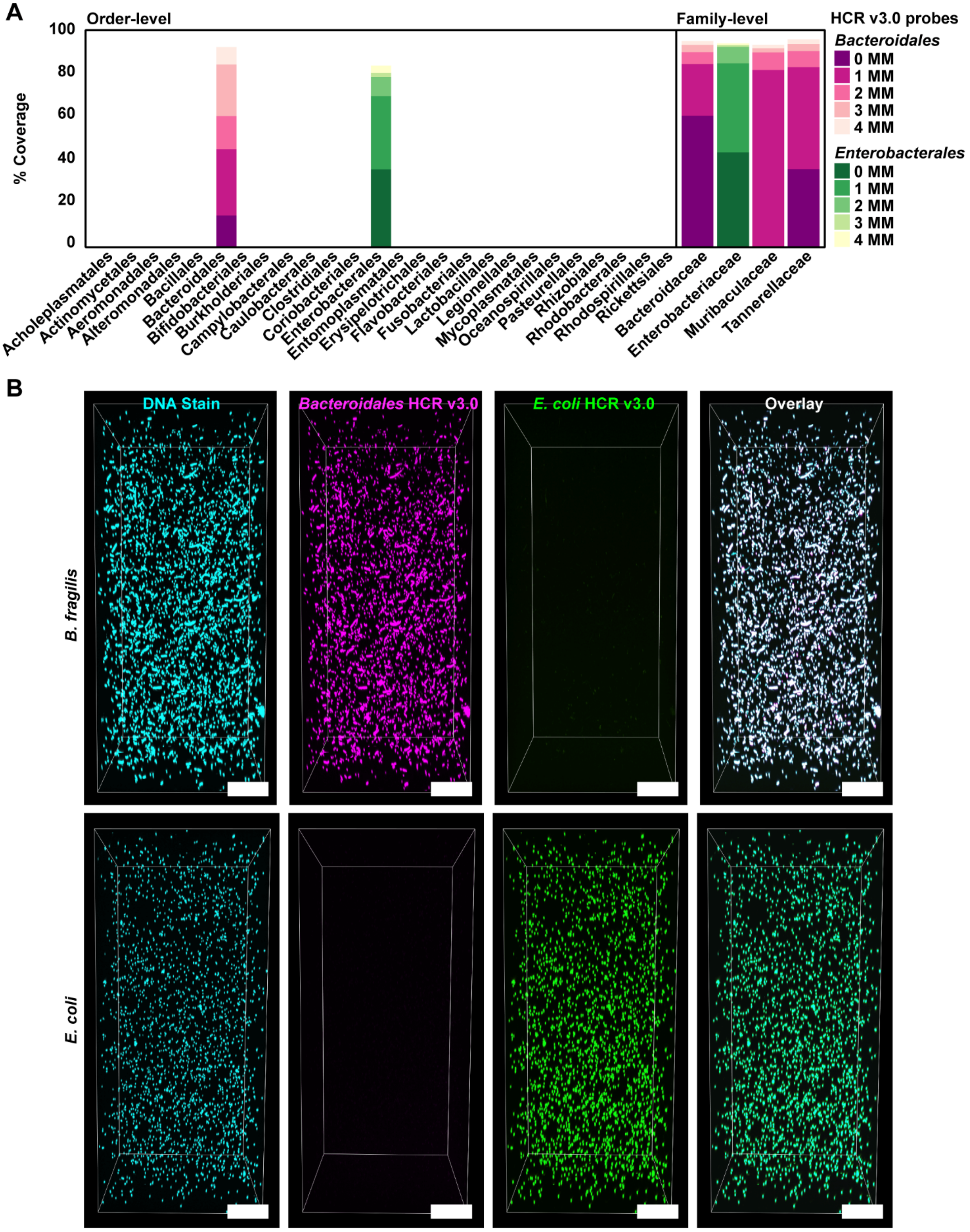


**Figure S9. Design and validation of the taxon-specific HCR v3.0 probes.** HCR v3.0 probes specific to *Bacteroidales* and *E. coli* were designed by Molecular Technologies to have no mismatches with the full 16S rRNA gene sequences of the gavaged *Bacteroides*/*Parabacteroides* spp. and *E. coli* isolates, respectively. (A) Probe coverage with no mismatches (0 MM) or up to 4 mismatches (1-4 MM) of all bacterial orders (left) and relevant bacterial families (right) detected (by sequencing) in the experimental mice. The coverage is expressed as % of sequences in SILVA database that align with the full 52-bp long HCR v3.0 probes when the specified number of mismatches is allowed. (B) Probe validation *in vitro* showing that the probes recognized their target, but did not cross-react with a model off-target even deep in the hydrogel. The validation was performed using single-isolate *in vitro* hydrogels, either *E. coli* or *B. fragilis*. Both hydrogels were stained with the two taxon-specific HCR v3.0 probes (*Bacteroidales*: magenta; *E. coli*: green) and counterstained with a nuclear stain (cyan). All scale bars 100 μm.

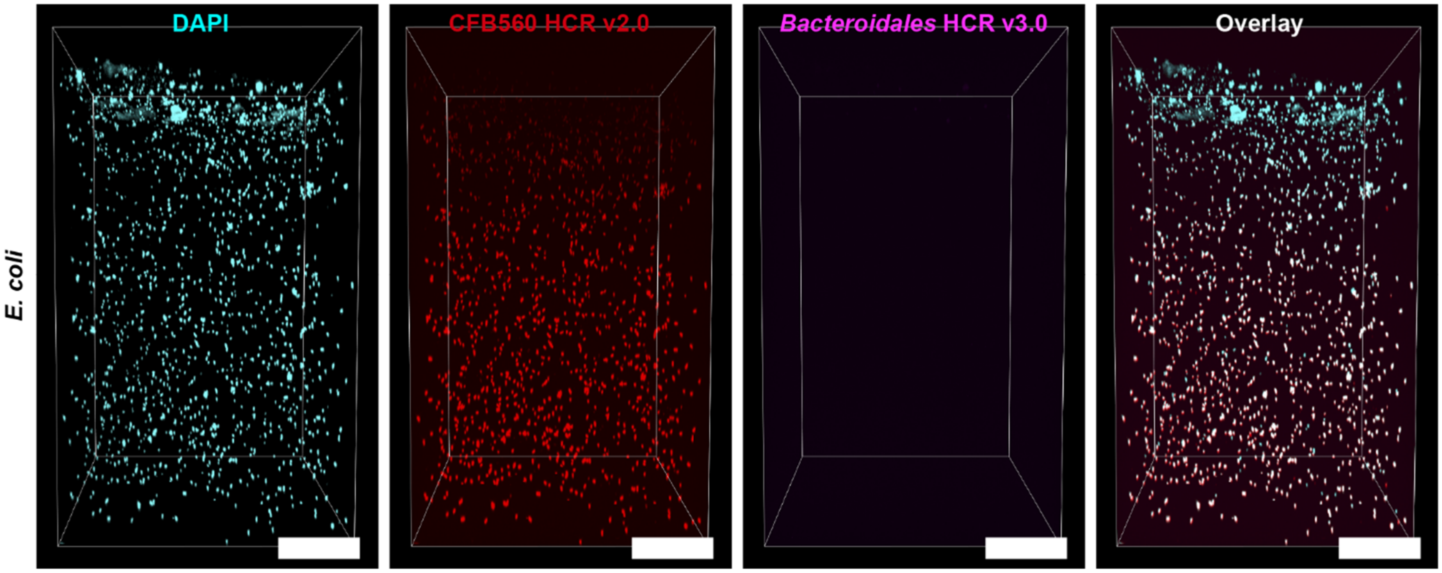


**Figure S10. Comparison of the taxon-specific *Bacteroidales* HCR v3.0 and CFB560 HCR v2.0 probes in an *in vitro* hydrogel with *E. coli*.** *E. coli* was stained DAPI for DNA (cyan), *Bacteroidetes*-specific CFB560 HCR v2.0 probe (red), and *Bacteroidales* HCR v3.0 probe (magenta). The CFB560 HCR v2.0 probe suffered from increasing false-positive signal amplification with depth, whereas the *Bacteroidales* HCR v3.0 probe successfully suppressed false-positive signal amplification deep into the hydrogel. All scale bars 100 µm.

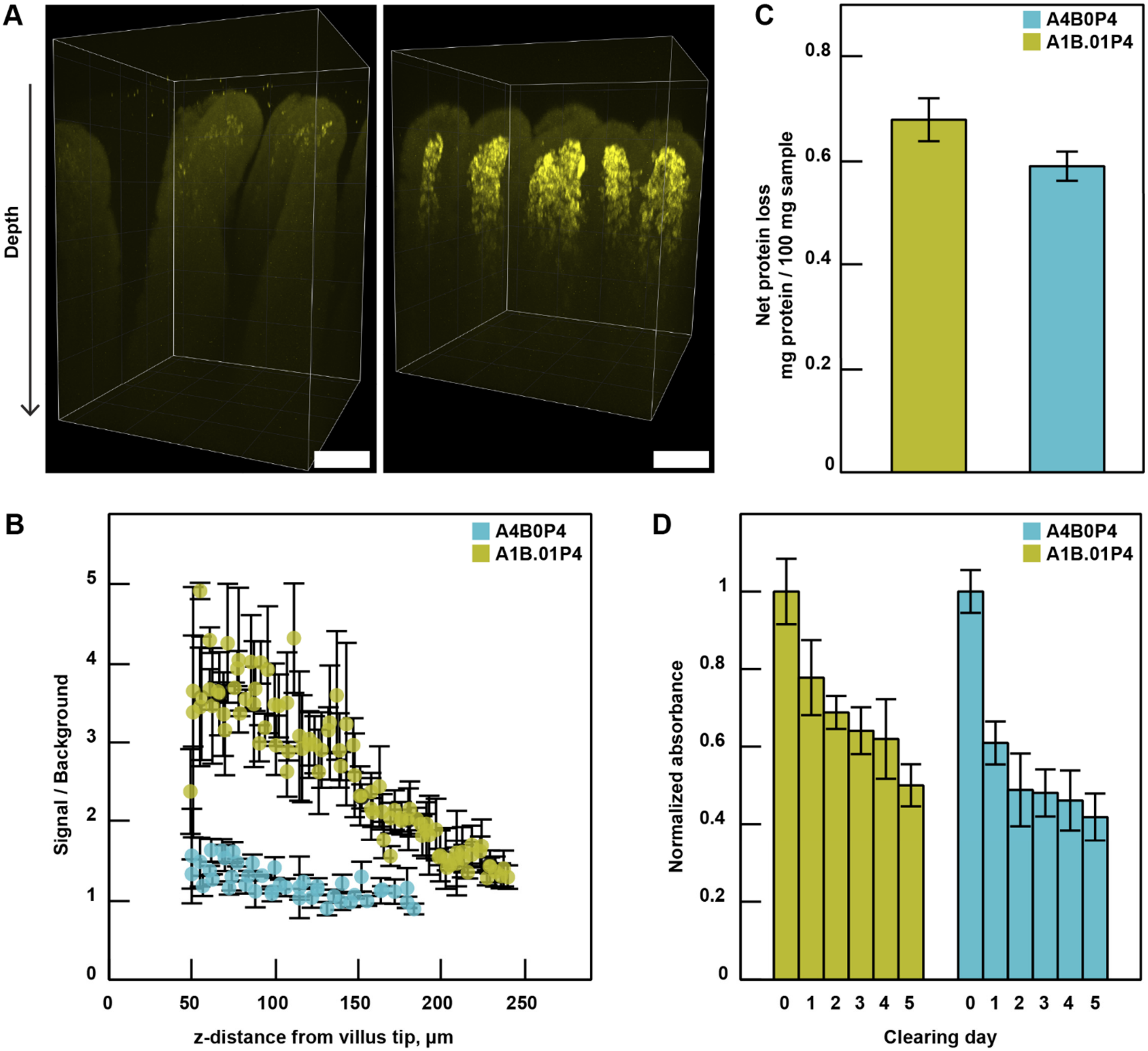


**Figure S11. Comparison of the more permeable A1B.01P4 and the less permeable A4B0P4 tissue gel formulations.** (A) 3D imaging z-stacks of anti-CD45 antibody staining of total immune cells (yellow) in a small intestine of CHOW-fed control mouse. In both images, the mucosa was protected by the same surface hydrogel, but the tissues were additionally fortified with either a 4% acrylamide + 4% paraformaldehyde formulation (A4B0P4) (left) or a 1% acrylamide + 0.01% bis-acrylamide + 4% paraformaldehyde (A1B.01P4) hydrogel formulation (right). All scale bars 100 µm. (B) Quantification of antibody penetration into the hydrogel-tissue hybrids shown in panel (A), demonstrating that A1B.01P4 chemistry increased the antibody signal/background ratio deep in the sample. (C) Net protein loss during clearing shows that a lower percentage of acrylamide hydrogel did not increase protein loss. (D) Normalized tissue absorbance during clearing shows that tissue clarity was similar for both hydrogel formulations.

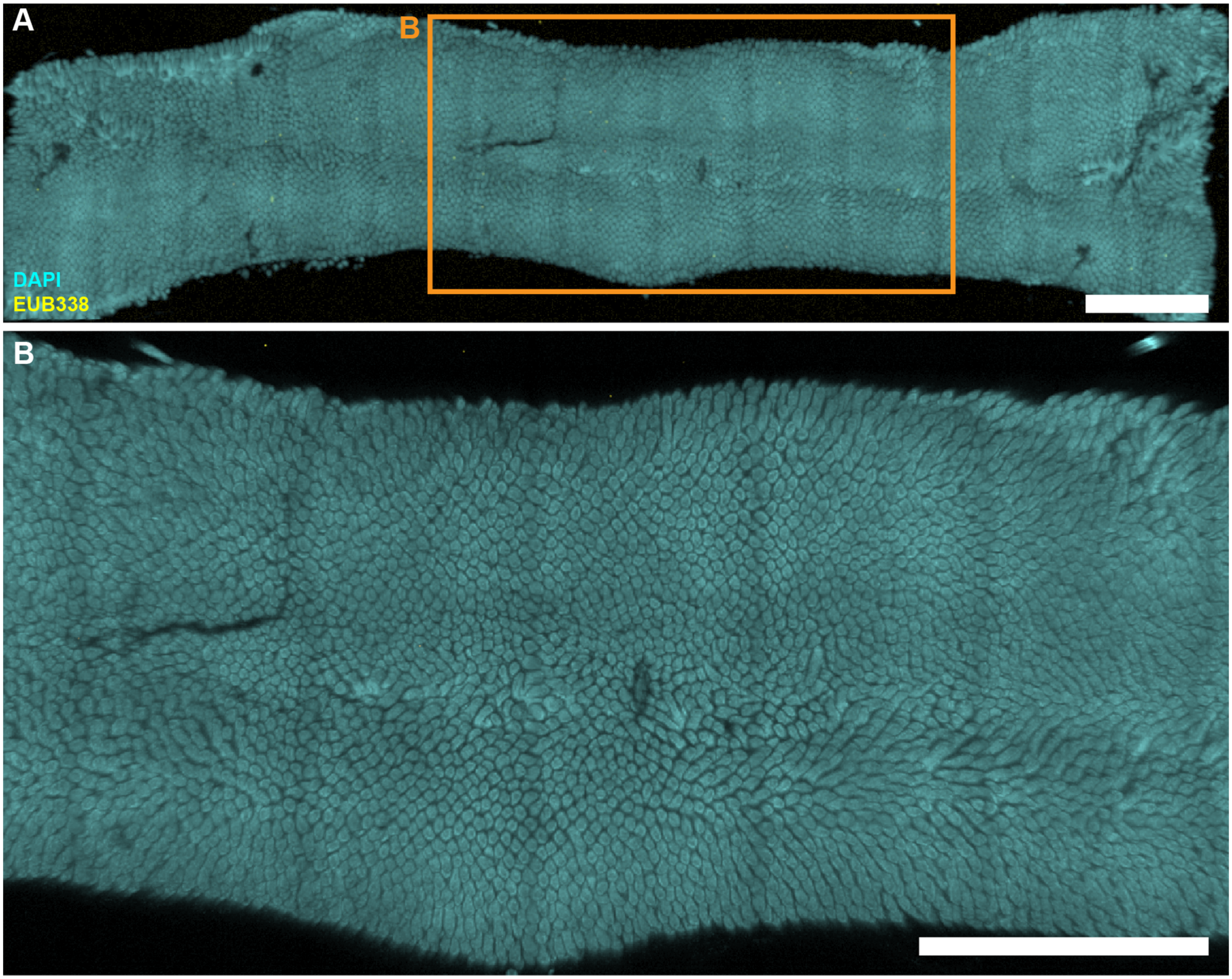


**Figure S12. Large-scale low-magnification fluorescence imaging of empty jejunum from MAL+PBS mouse on day 28 of the experiment.** (A) Large-scale low-magnification tile scan of a whole empty jejunum segment without digesta showing DAPI staining of epithelial surface (cyan) and HCR v3.0 staining of total bacteria (yellow). (B) Enlarged portion of the image shown in A (orange rectangle). All scale bars 2 mm.

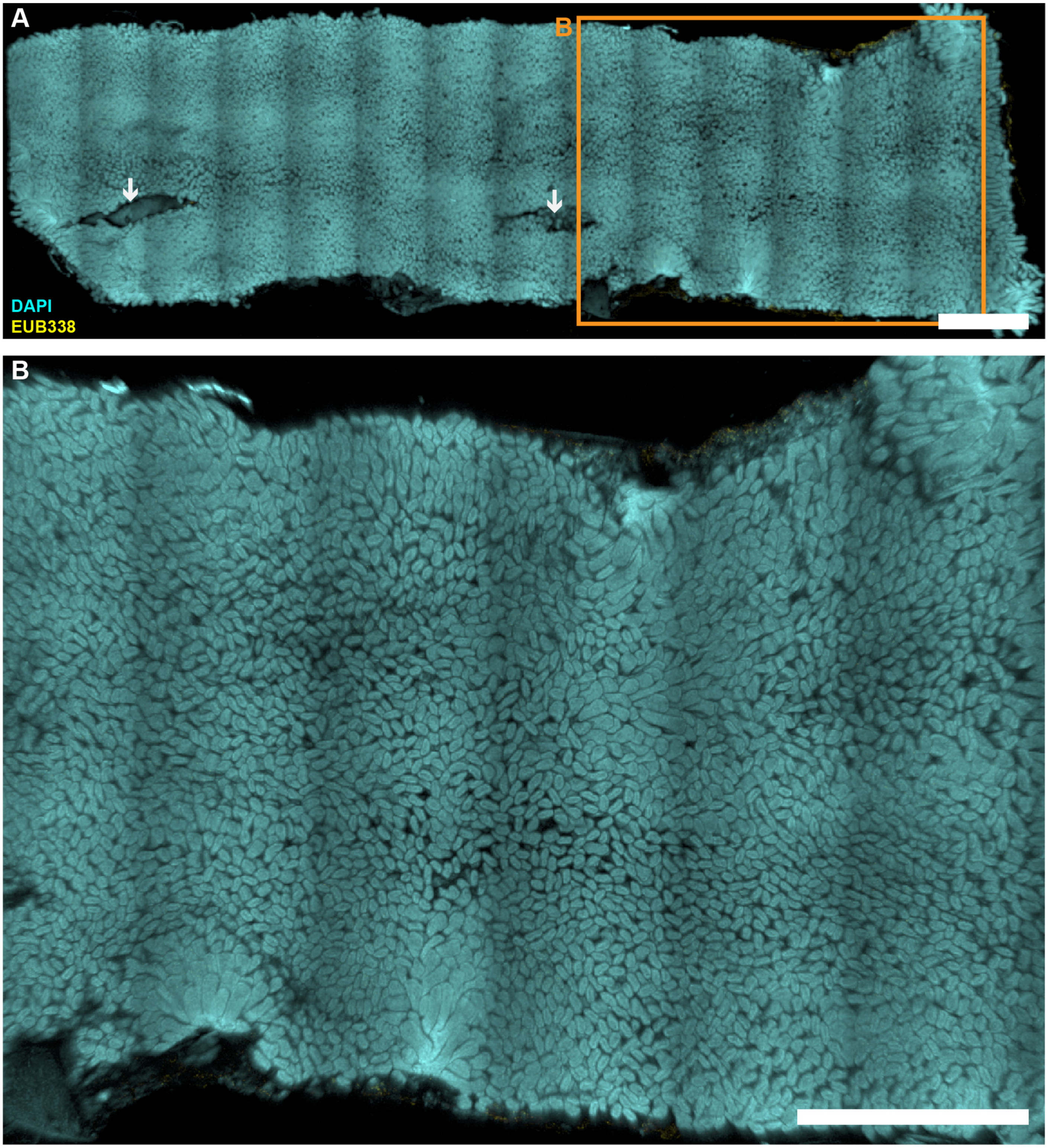


**Figure S13. Large-scale low-magnification fluorescence imaging of empty jejunum from MAL+PBS mouse on day 29 of the experiment.** (A) Large-scale low-magnification tile scan of whole empty jejunum segment without digesta showing DAPI staining of epithelial surface (cyan) and HCR v3.0 staining of total bacteria (yellow). Streaks (white arrows) represent scissor incisions during sample preparation. (B) Enlarged portion of the image shown in A (orange rectangle). All scale bars 2 mm.

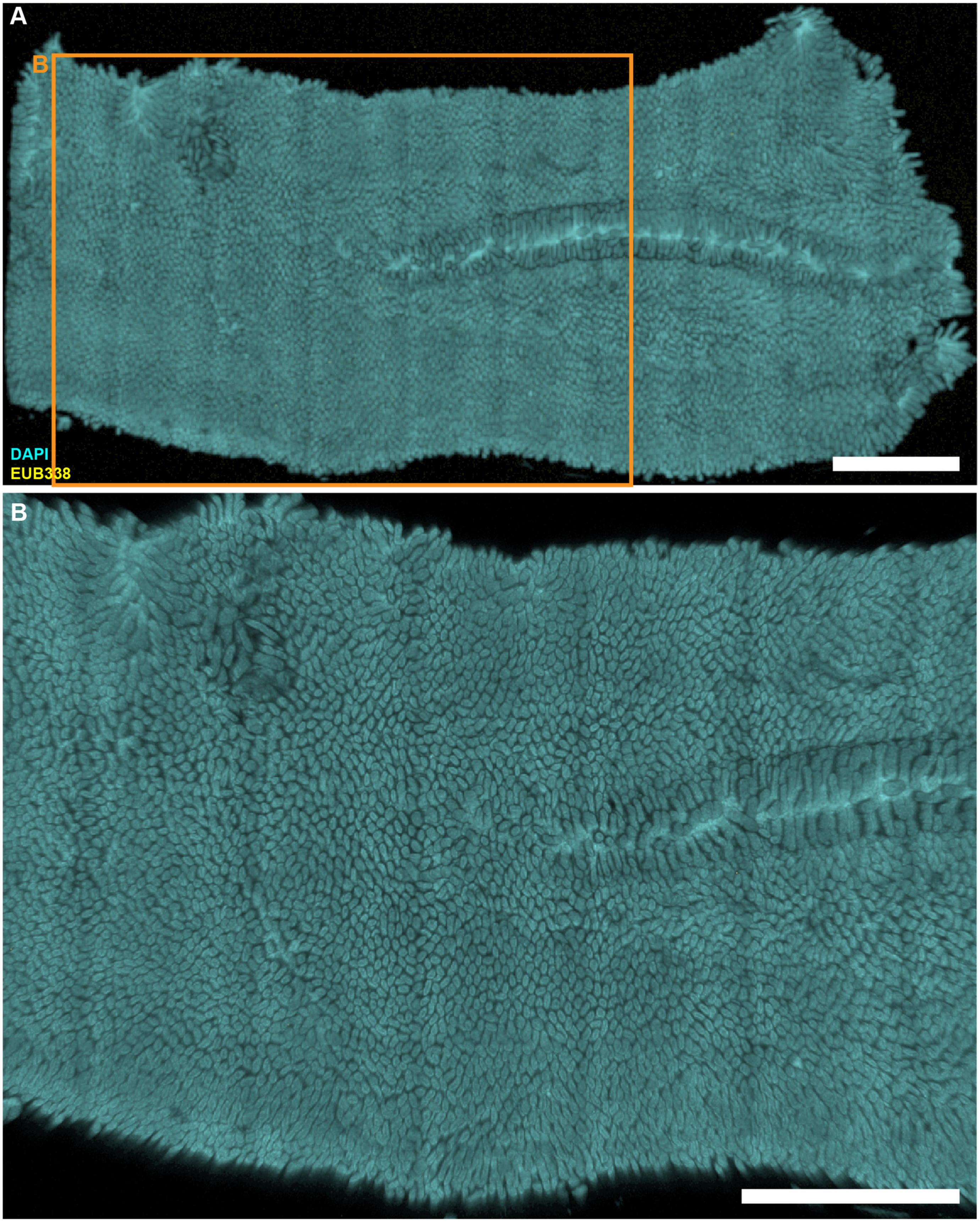


**Figure S14. Large-scale low-magnification fluorescence imaging of empty jejunum from MAL+PBS mouse on day 30 of the experiment.** (A) Large-scale low-magnification tile scan of whole empty jejunum segment without digesta showing DAPI staining of epithelial surface (cyan) and HCR v3.0 staining of total bacteria (yellow). (B) Enlarged portion of the image shown in A (orange rectangle). All scale bars 2 mm.

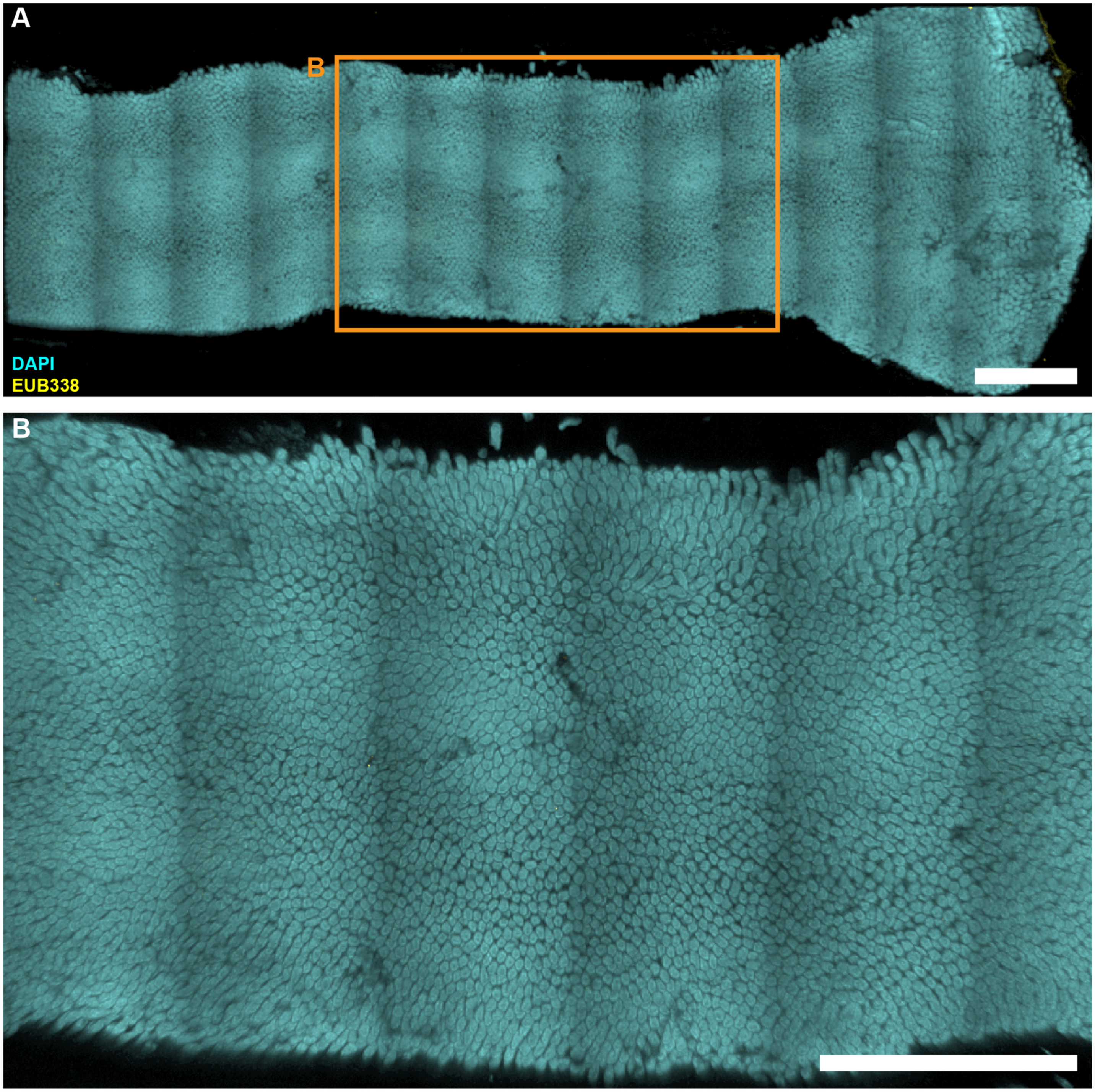


**Figure S15. Large-scale low-magnification fluorescence imaging of empty jejunum from MAL+PBS mouse on day 31 of the experiment.** (A) Large-scale low-magnification tile scan of whole empty jejunum segment without digesta showing DAPI staining of epithelial surface (cyan) and HCR v3.0 staining of total bacteria (yellow). (B) Enlarged portion of the image shown in A (orange rectangle). All scale bars 2 mm.

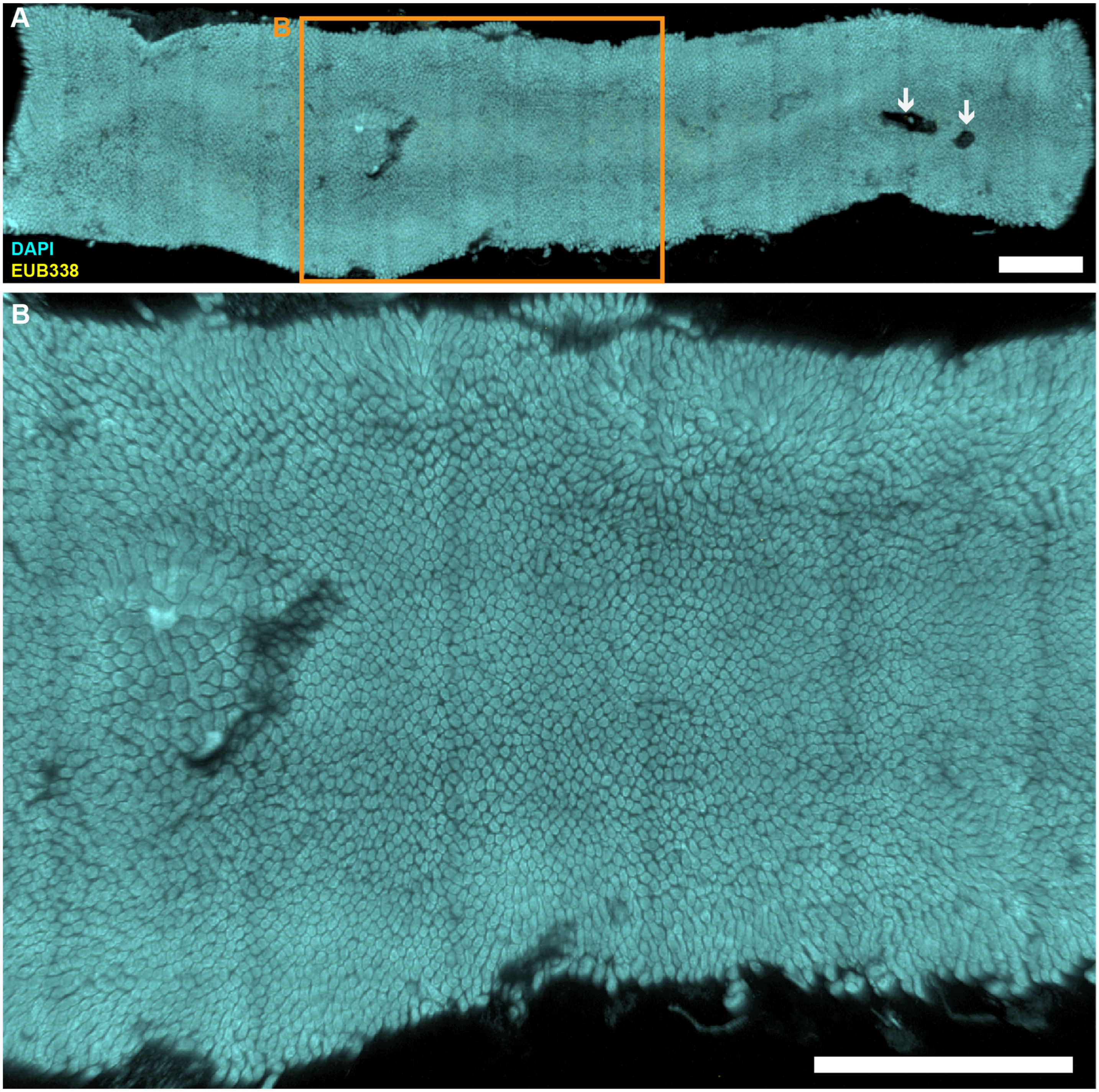


**Figure S16. Large-scale low-magnification fluorescence imaging of empty jejunum from MAL+BAC mouse on day 28 of the experiment.** (A) Large-scale low-magnification tile scan of whole empty (without visible digesta) jejunum segment showing DAPI staining of epithelial surface (cyan) and HCR v3.0 staining of total bacteria (yellow). Dark spots (white arrows) may represent large surface aggregates that were not discernable by naked eye (Fig. 4A). (B) Enlarged portion of the image shown in A (orange rectangle). All scale bars 2 mm.

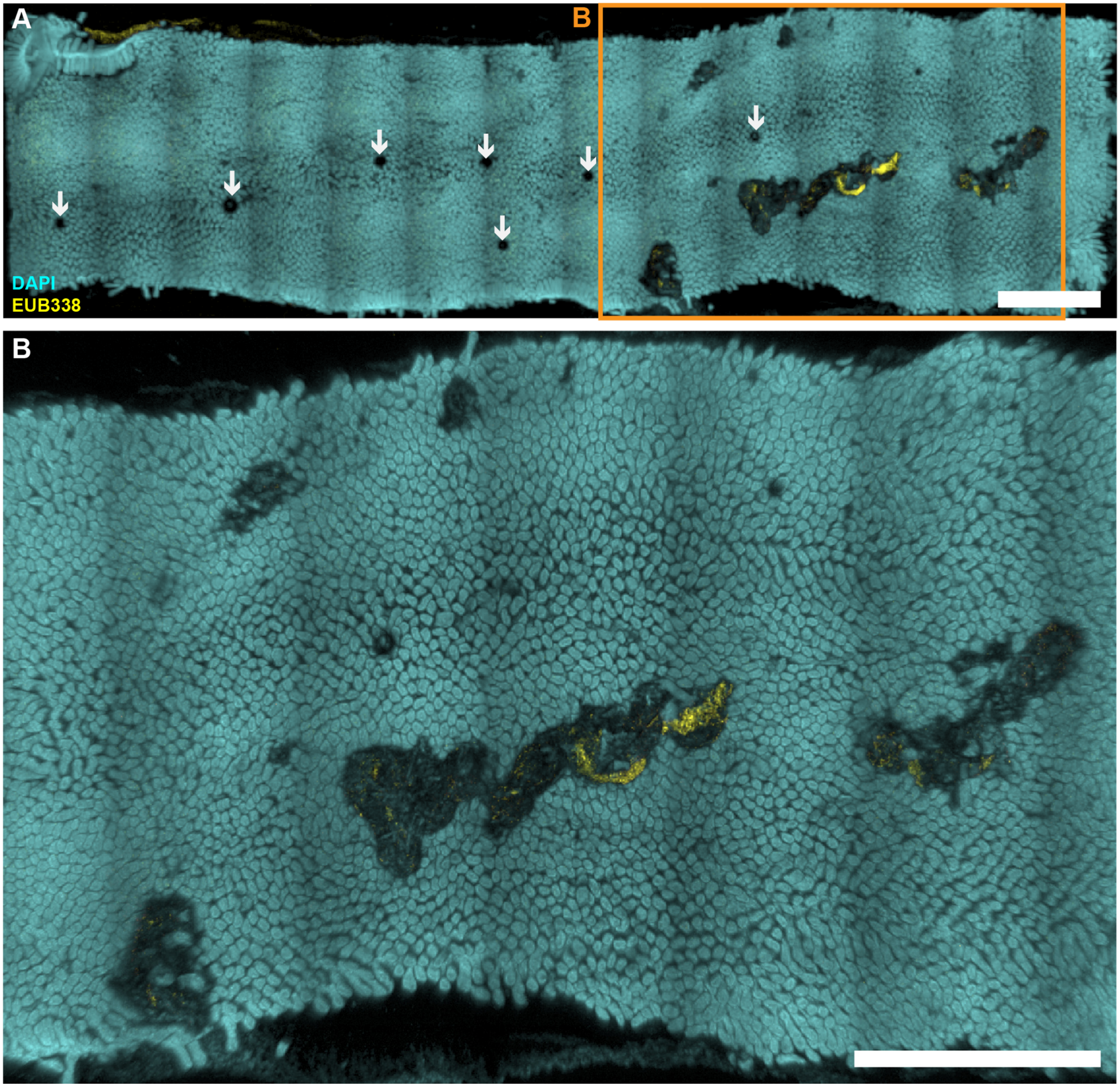


**Figure S17. Large-scale low-magnification fluorescence imaging of empty jejunum from MAL+BAC mouse on day 29 of the experiment.** (A) Large-scale low-magnification tile scan of whole empty jejunum segment without digesta showing DAPI staining of epithelial surface (cyan) and HCR v3.0 staining of total bacteria (yellow). Small circles (white arrows) are imaging artifacts created by air bubbles trapped under the cover slip. (B) Enlarged portion of the image shown in A (orange rectangle). All scale bars 2 mm.

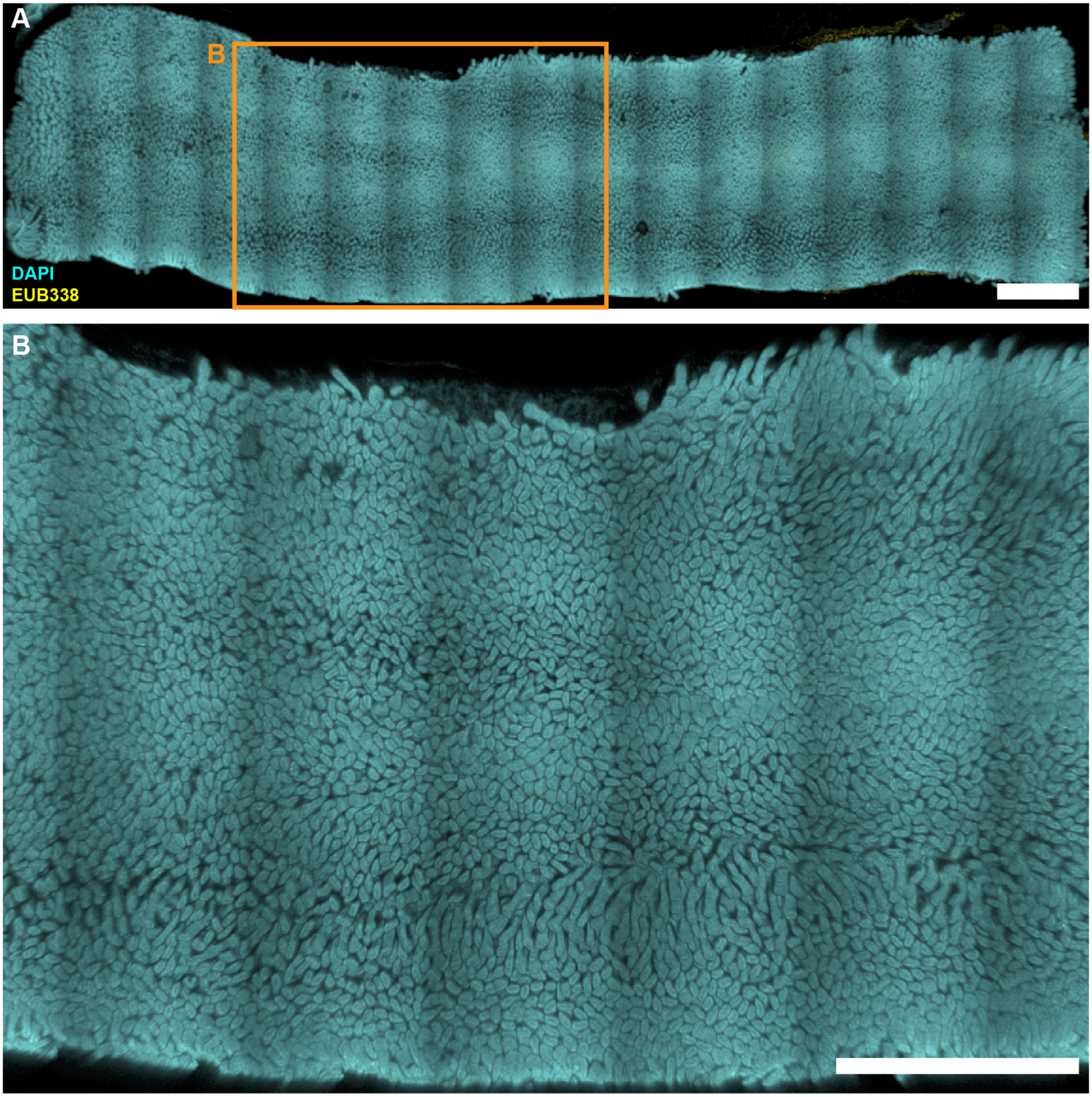


**Figure S18. Large-scale low-magnification fluorescence imaging of empty jejunum from MAL+BAC mouse on day 30 of the experiment.** (A) Large-scale low-magnification tile scan of whole empty jejunum segment without digesta showing DAPI staining of epithelial surface (cyan) and HCR v3.0 staining of total bacteria (yellow). (B) Enlarged portion of the image shown in A (orange rectangle). All scale bars 2 mm.

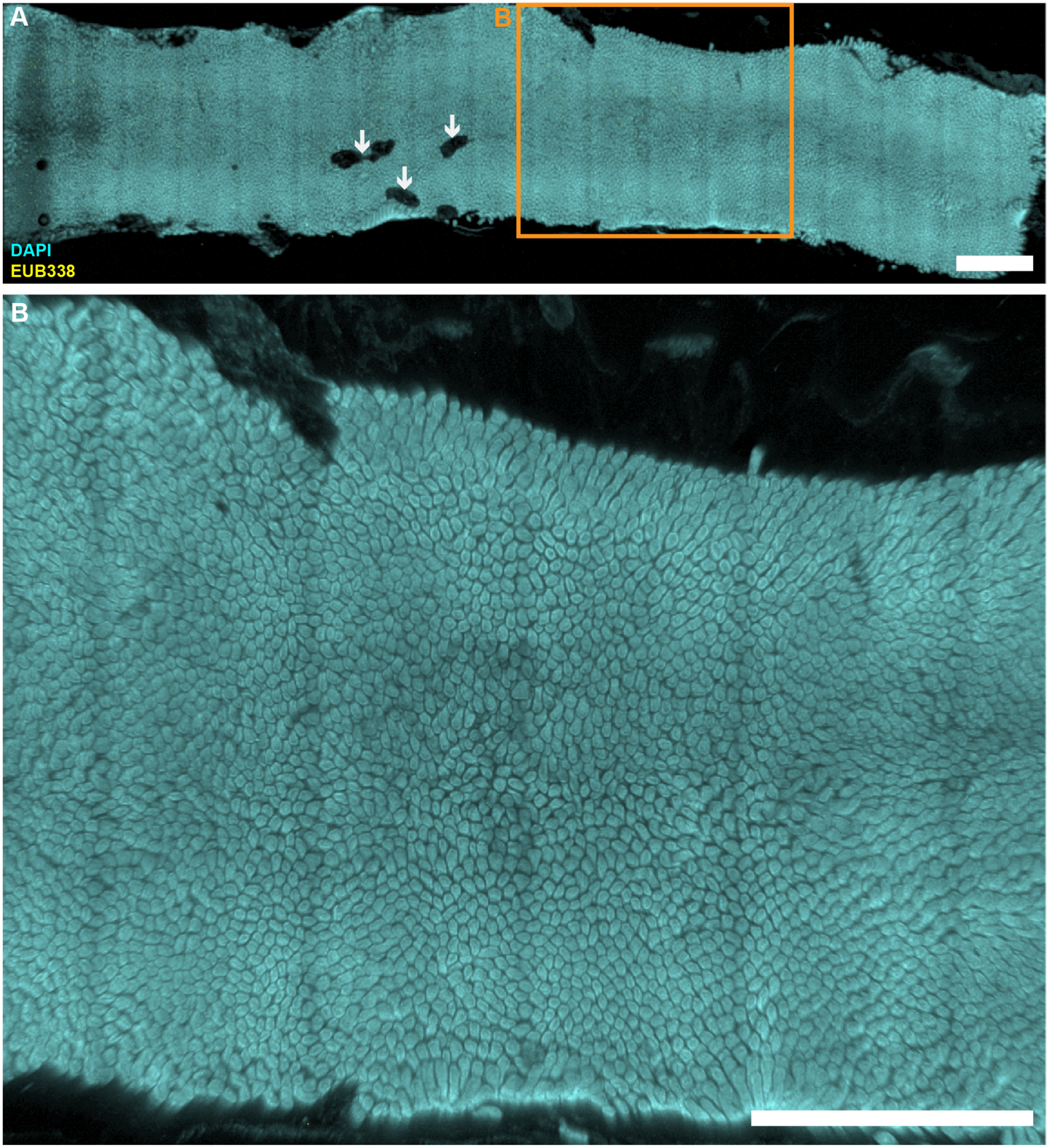


**Figure S19. Large-scale low-magnification fluorescence imaging of empty jejunum from MAL+BAC mouse on day 31 of the experiment.** (A) Large-scale low-magnification tile scan of whole empty (without visible digesta) jejunum segment without digesta showing DAPI staining of epithelial surface (cyan) and HCR v3.0 staining of total bacteria (yellow). Dark spots (white arrows) may represent large surface aggregates that were not discernable by naked eye (Fig. 4A). (B) Enlarged portion of the image shown in A (orange rectangle). All scale bars 2 mm.

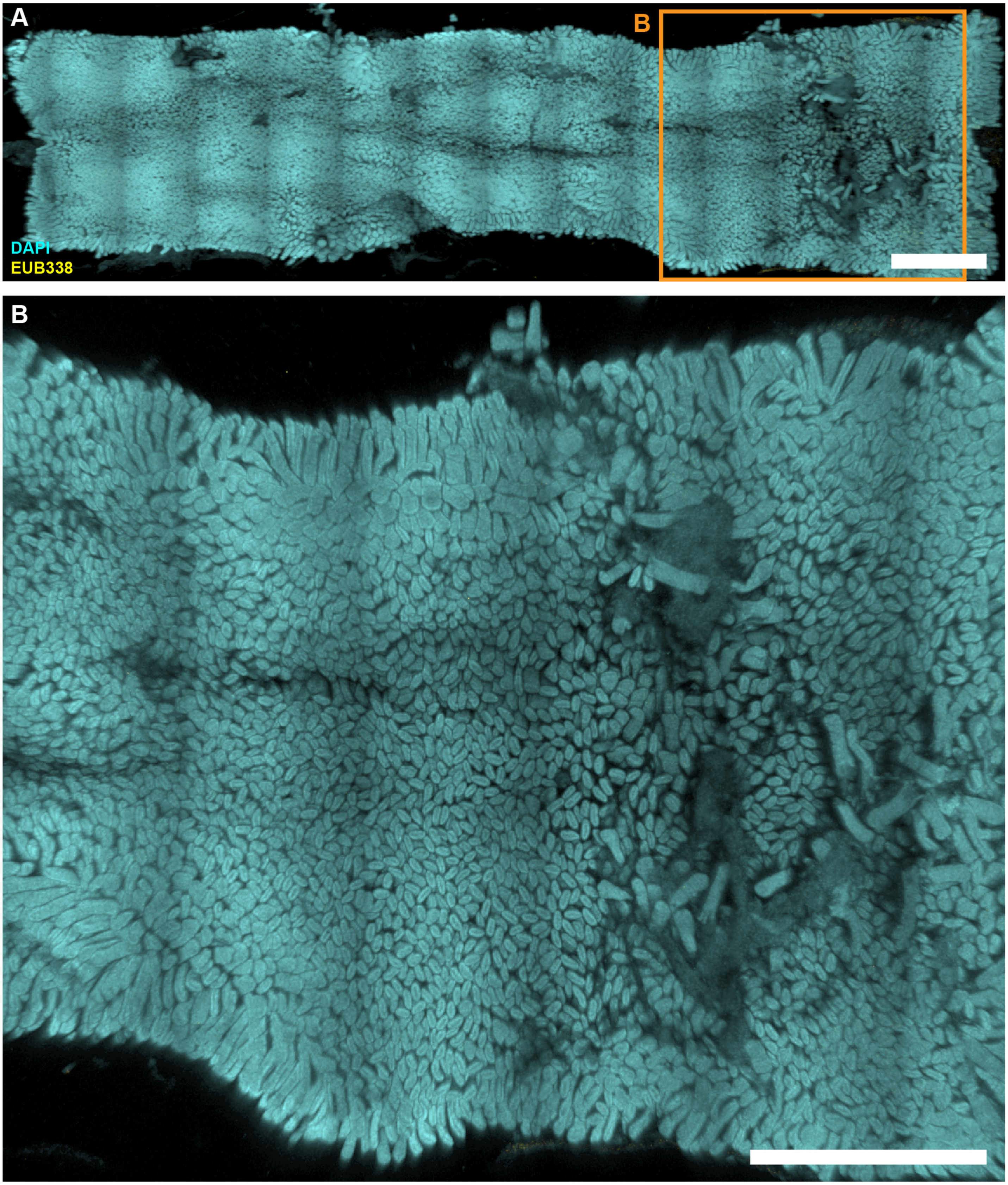


**Figure S20. Large-scale low-magnification fluorescence imaging of empty jejunum from MAL+EC mouse on day 28 of the experiment.** (A) Large-scale low-magnification tile scan of whole empty jejunum segment without digesta showing DAPI staining of epithelial surface (cyan) and HCR v3.0 staining of total bacteria (yellow). (B) Enlarged portion of the image shown in A (orange rectangle). In this part of the sample, villus loss is clearly visible. All scale bars 2 mm.

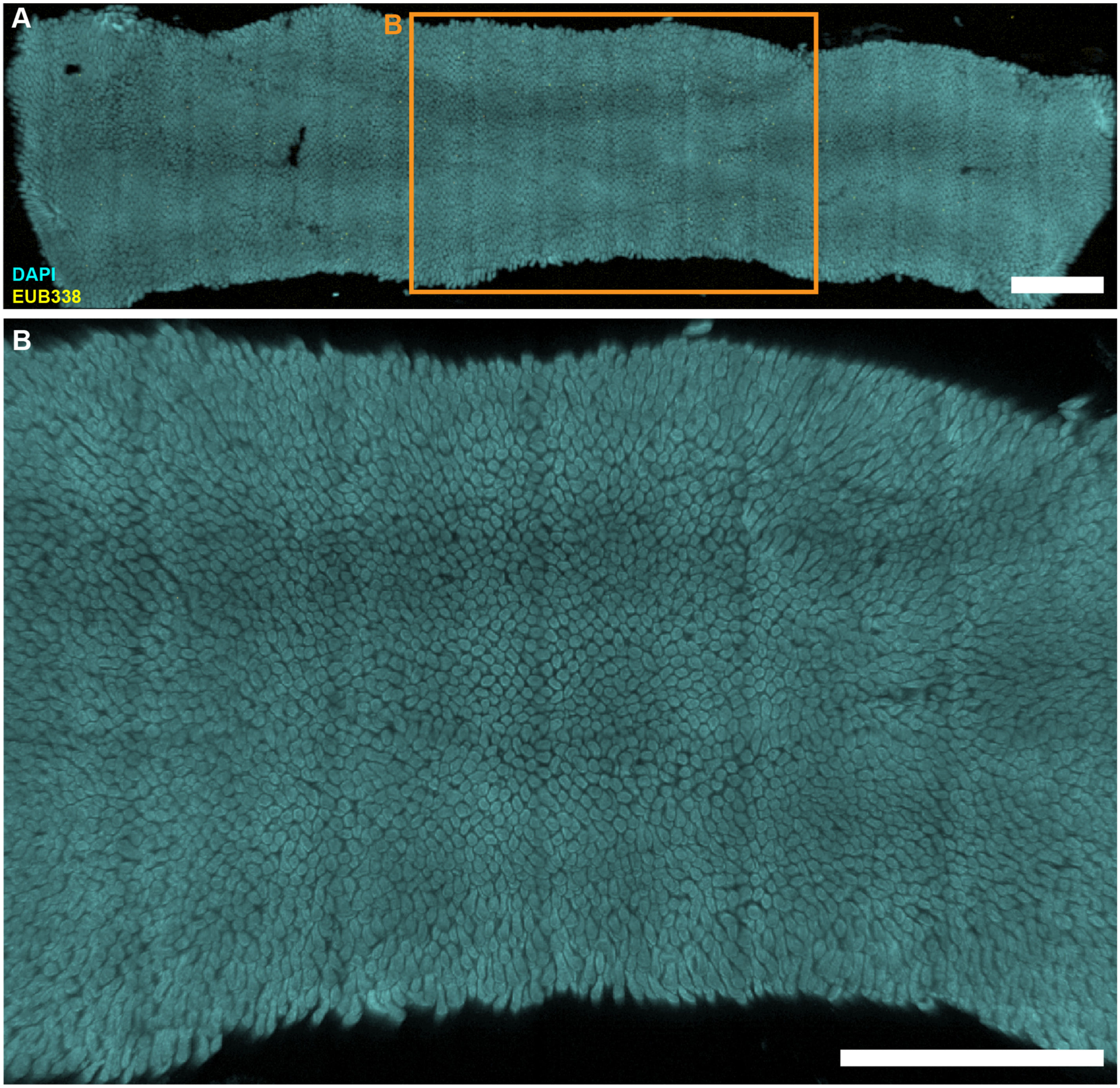


**Figure S21. Large-scale low-magnification fluorescence imaging of empty jejunum from MAL+EC mouse on day 29 of the experiment.** (A) Large-scale low-magnification tile scan of whole empty jejunum segment without digesta showing DAPI staining of epithelial surface (cyan) and HCR v3.0 staining of total bacteria (yellow). Small yellow dots are rendering artifacts rather than bacteria, and they are not visible when the image is enlarged as seen in B. (B) Enlarged portion of the image shown in A (orange rectangle). All scale bars 2 mm.

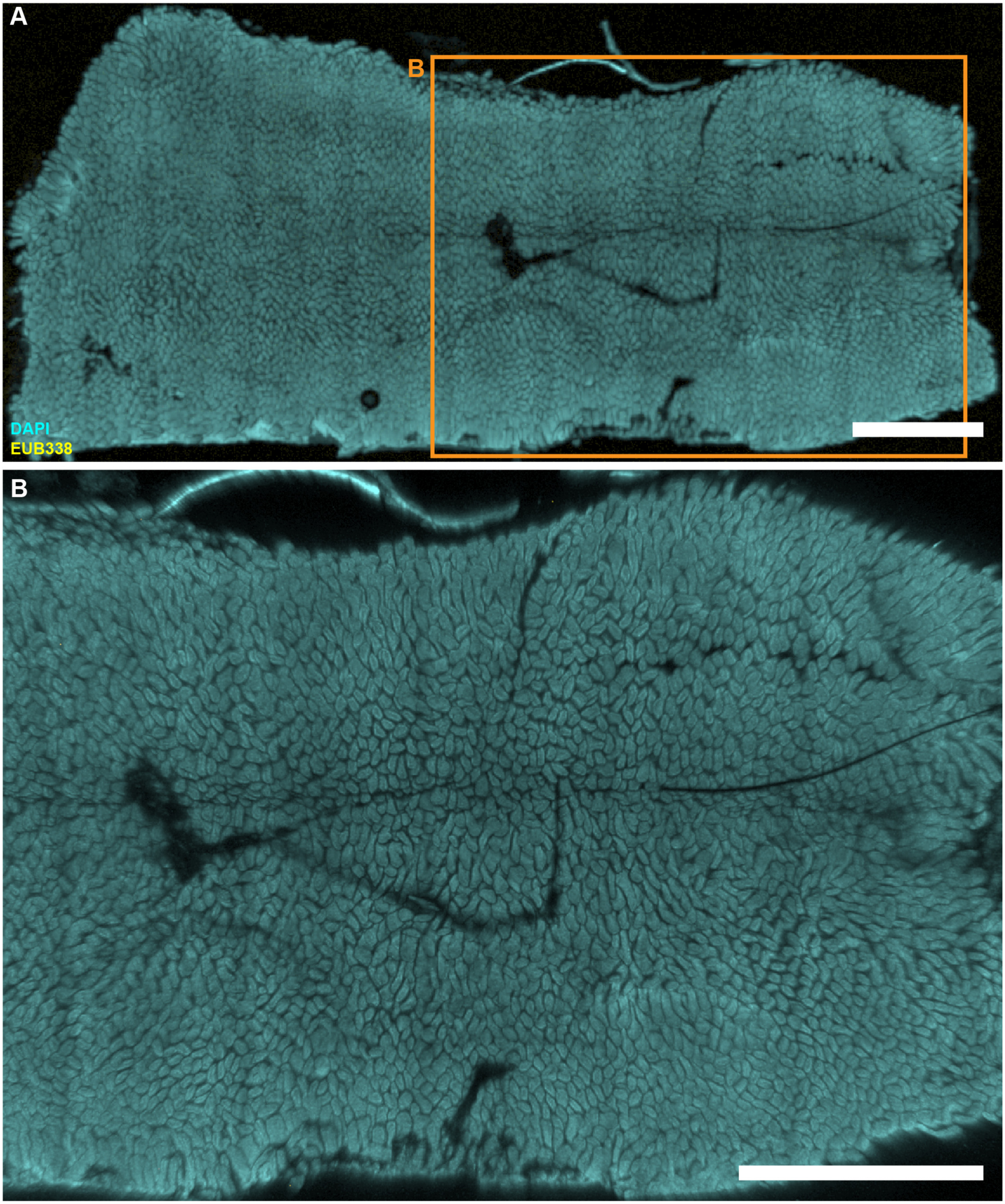


**Figure S22. Large-scale low-magnification fluorescence imaging of empty jejunum from MAL+EC mouse on day 30 of the experiment.** (A) Large-scale low-magnification tile scan of whole empty jejunum segment without digesta showing DAPI staining of epithelial surface (cyan) and HCR v3.0 staining of total bacteria (yellow). (B) Enlarged portion of the image shown in A (orange rectangle). Fine lines are shadows cast by mouse hairs that were either ingested or adhered to the epithelium during sample preparation. All scale bars 2 mm.

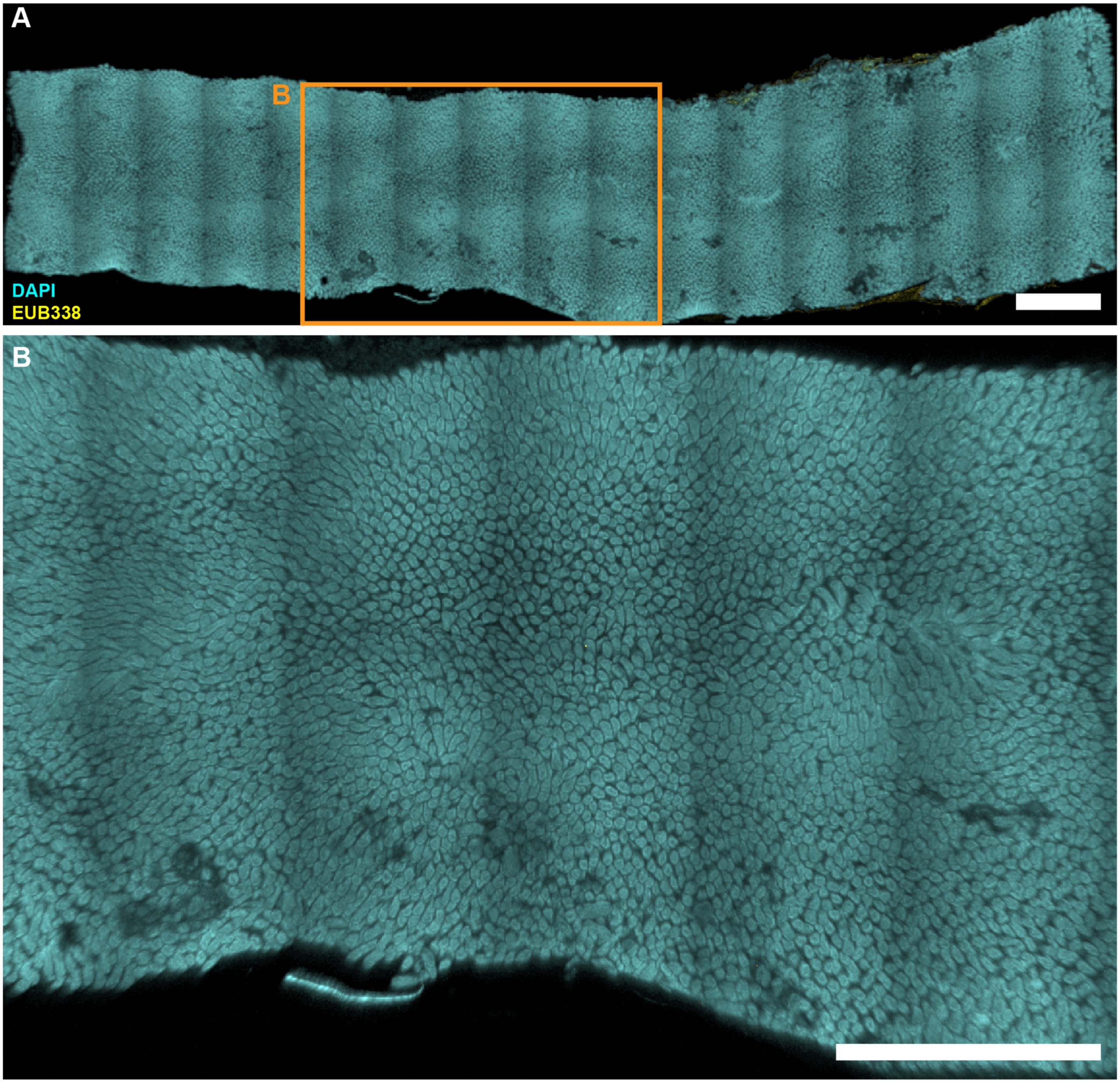


**Figure S23. Large-scale low-magnification fluorescence imaging of empty jejunum from MAL+EC mouse on day 31 of the experiment.** (A) Large-scale low-magnification tile scan of whole empty jejunum segment without digesta showing DAPI staining of epithelial surface (cyan) and HCR v3.0 staining of total bacteria (yellow). (B) Enlarged portion of the image shown in A (orange rectangle). All scale bars 2 mm.

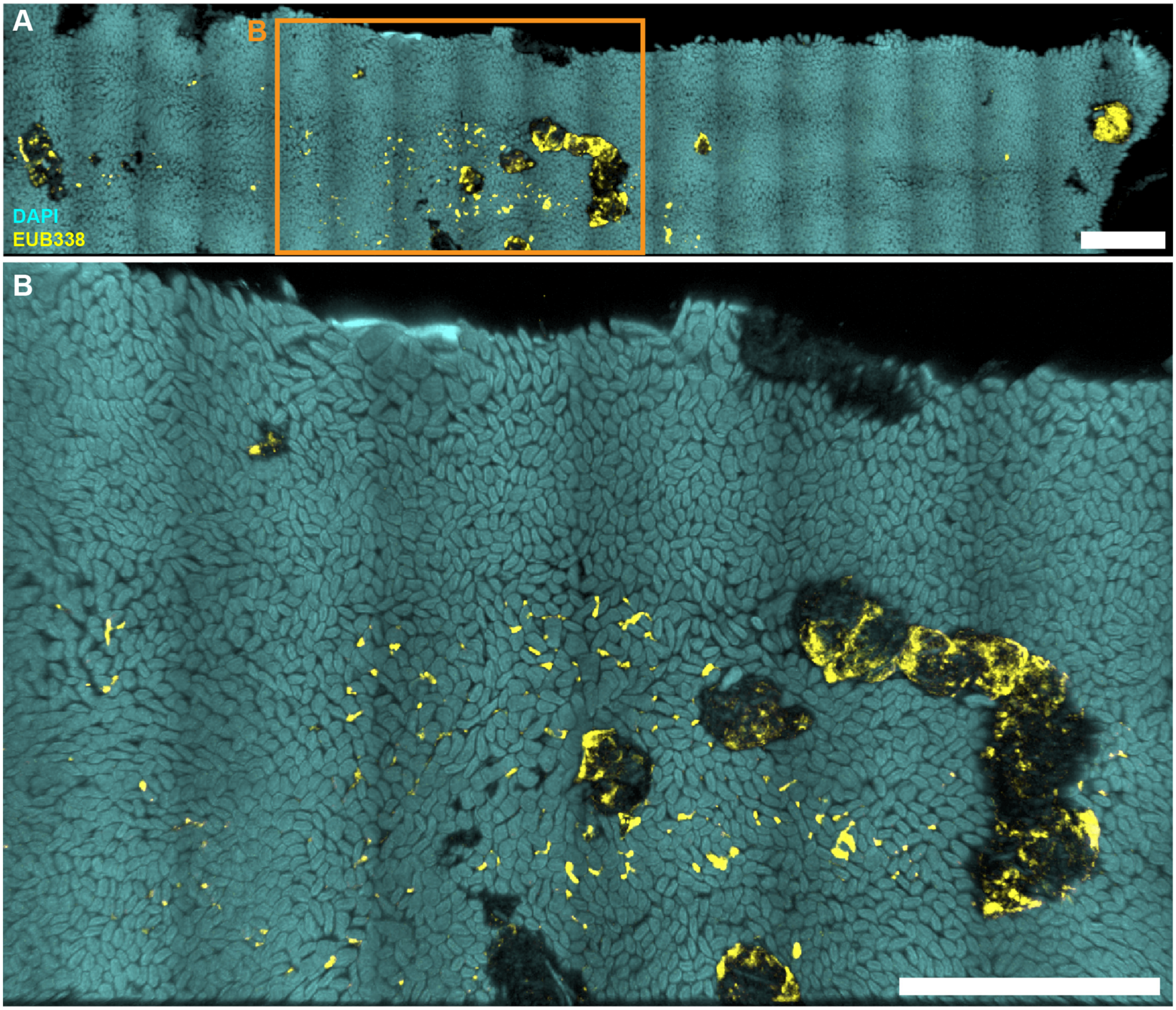


**Figure S24. Large-scale low-magnification fluorescence imaging of empty jejunum from MAL+EC&BAC mouse on day 28 of the experiment.** (A) Large-scale low-magnification tile scan of whole empty jejunum segment without digesta showing DAPI staining of epithelial surface (cyan) and HCR v3.0 staining of total bacteria (yellow). (B) Enlarged portion of the image shown in A (orange rectangle). All scale bars 2 mm.

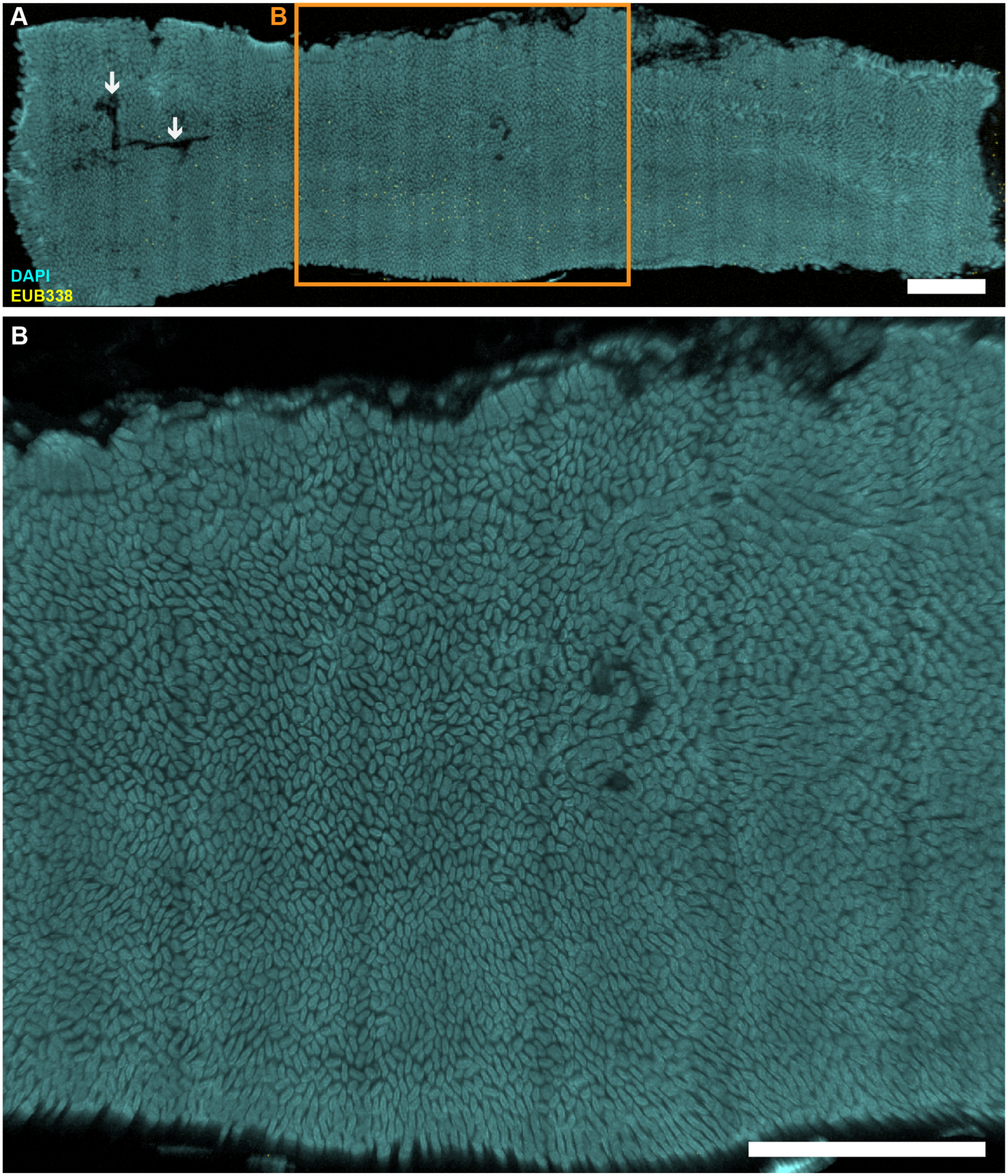


**Figure S25. Large-scale low-magnification fluorescence imaging of empty jejunum from MAL+EC&BAC mouse on day 29 of the experiment.** (A) Large-scale low-magnification tile scan of whole empty jejunum segment without digesta showing DAPI staining of epithelial surface (cyan) and HCR v3.0 staining of total bacteria (yellow). Dark streaks (white arrows) represent scissor incisions during sample preparation. Small yellow dots are rendering artifacts rather than bacteria, and they are not visible when the image is enlarged as seen in B. (B) Enlarged portion of the image shown in A (orange rectangle). All scale bars 2 mm.

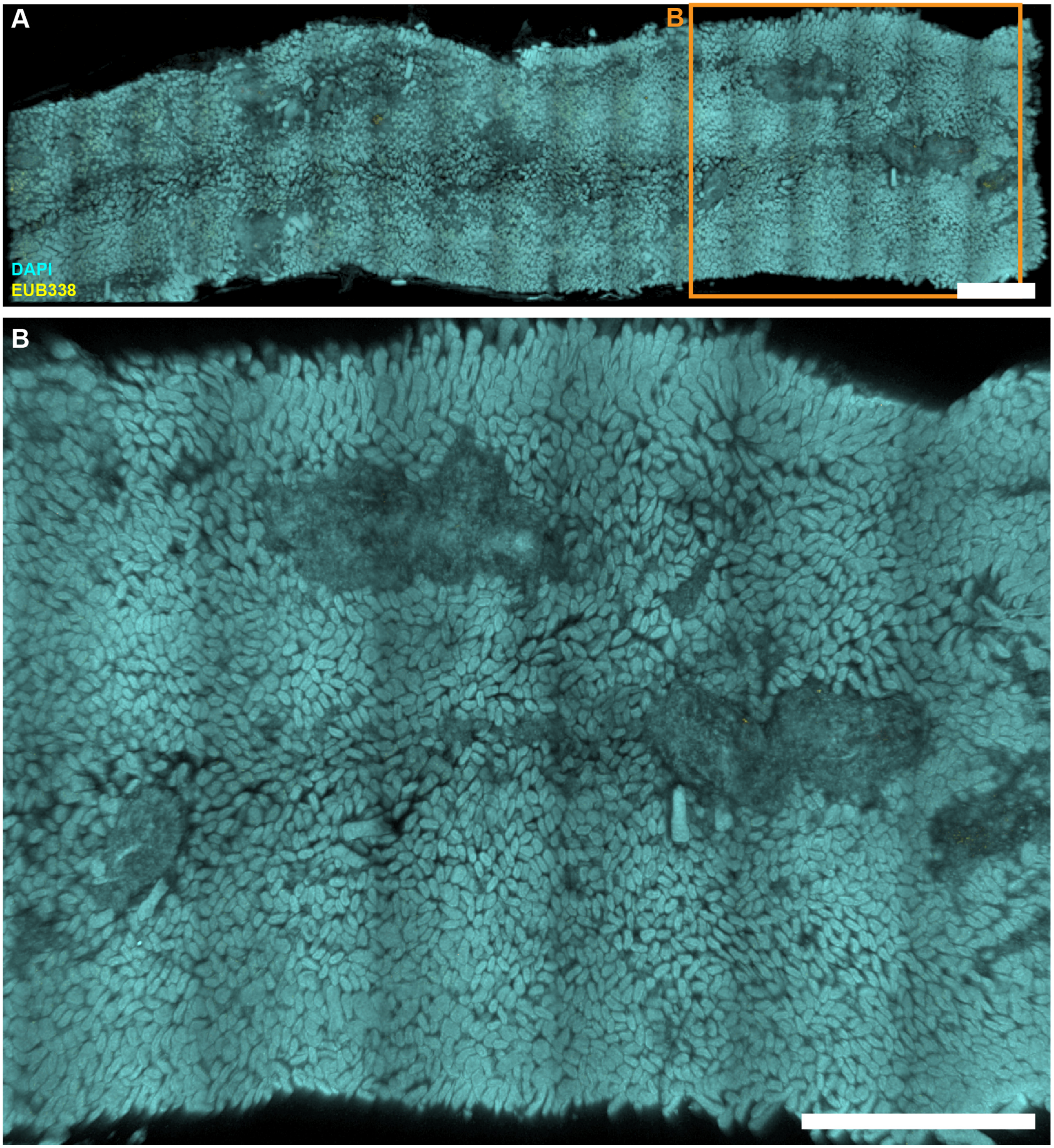


**Figure S26. Large-scale low-magnification fluorescence imaging of empty jejunum from MAL+EC&BAC mouse on day 31 of the experiment.** (A) Large-scale low-magnification tile scan of whole empty jejunum segment without digesta showing DAPI staining of epithelial surface (cyan) and HCR v3.0 staining of total bacteria (yellow). (B) Enlarged portion of the image shown in A (orange rectangle). All scale bars 2 mm.

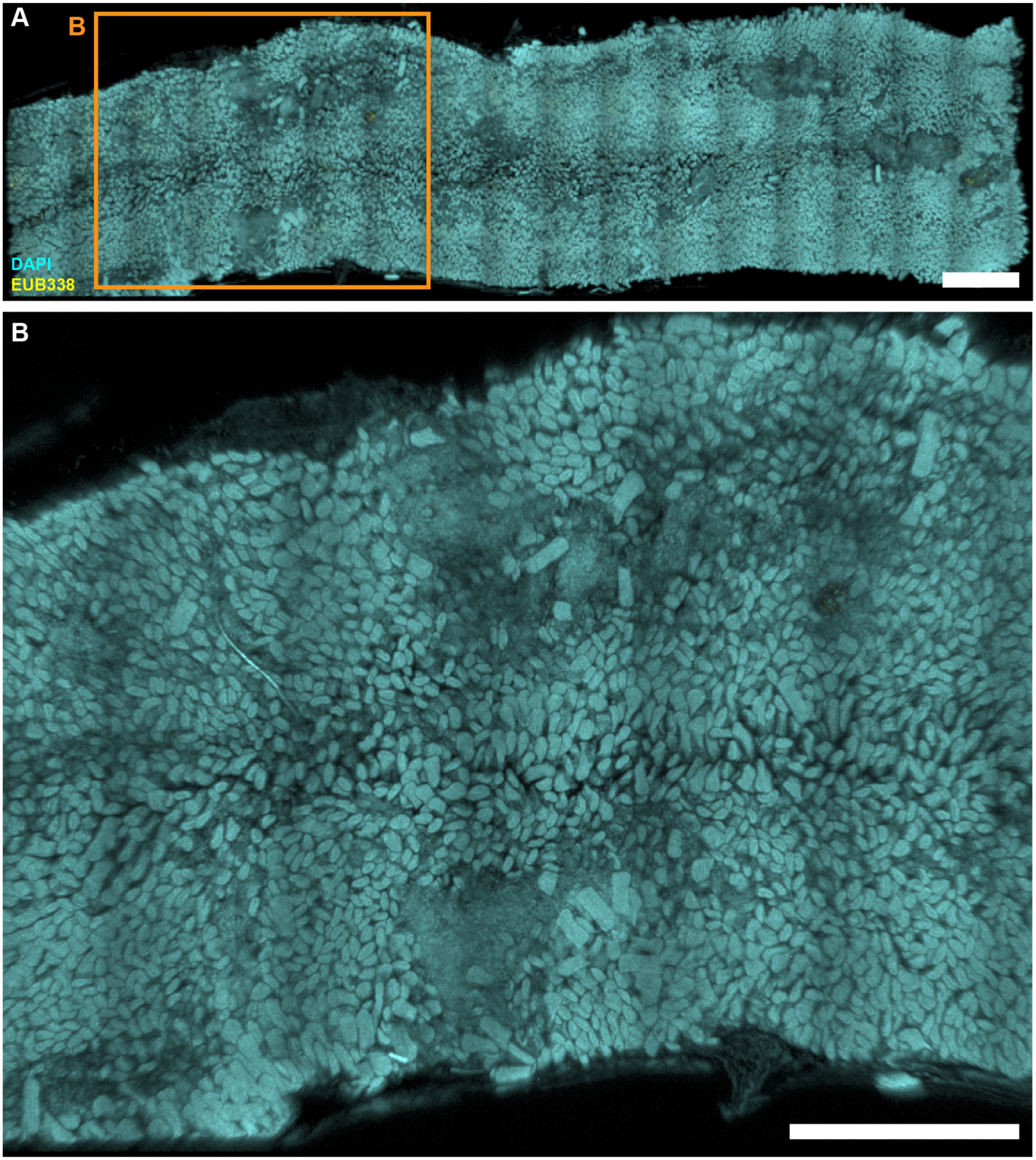


**Figure S27. Large-scale low-magnification fluorescence imaging of empty jejunum from MAL+EC&BAC mouse on day 31 of the experiment.** (A) Large-scale low-magnification tile scan of whole empty jejunum segment without digesta showing DAPI staining of epithelial surface (cyan) and HCR v3.0 staining of total bacteria (yellow). (B) Enlarged portion of the image shown in A (orange rectangle). In this part of the sample, villus loss is clearly visible. All scale bars 2 mm.

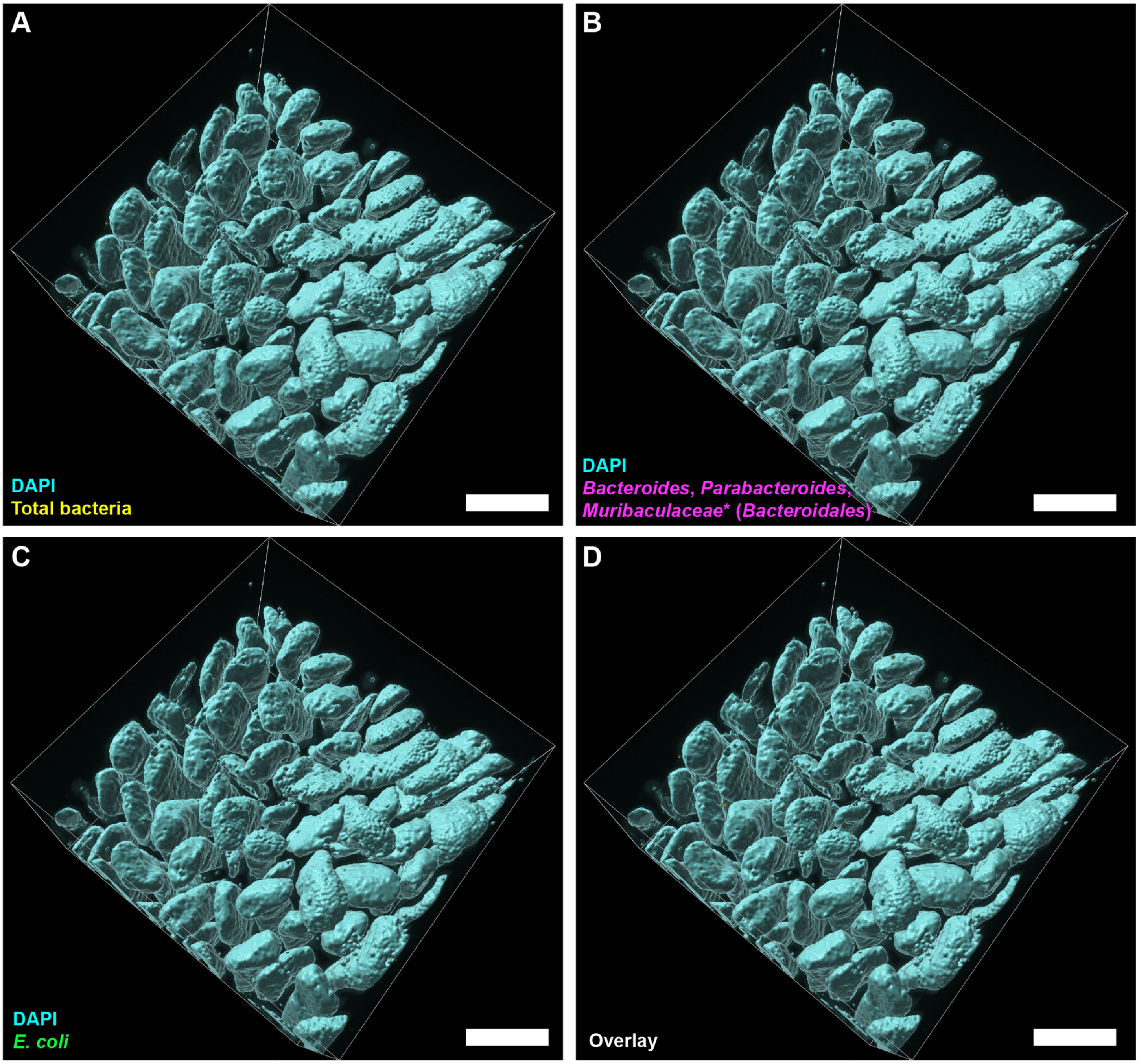


**Figure S28. High-magnification 3D fluorescence imaging of bacteria after 1-h fast in an empty jejunum of MAL+PBS mouse on day 29 of the experiment.** (A-D) 3D rendering of the segmented and filtered surfaces displaying DAPI staining of epithelium (cyan) and (A) HCR v3.0 staining for total bacteria (yellow), (B) HCR v3.0 staining for *Bacteroidales*, (C) HCR v3.0 staining for *E. coli* (green), and (D) overlay with all groups of bacteria. No significant number of bacteria was detected in this field of view. All scale bars 200 µm.

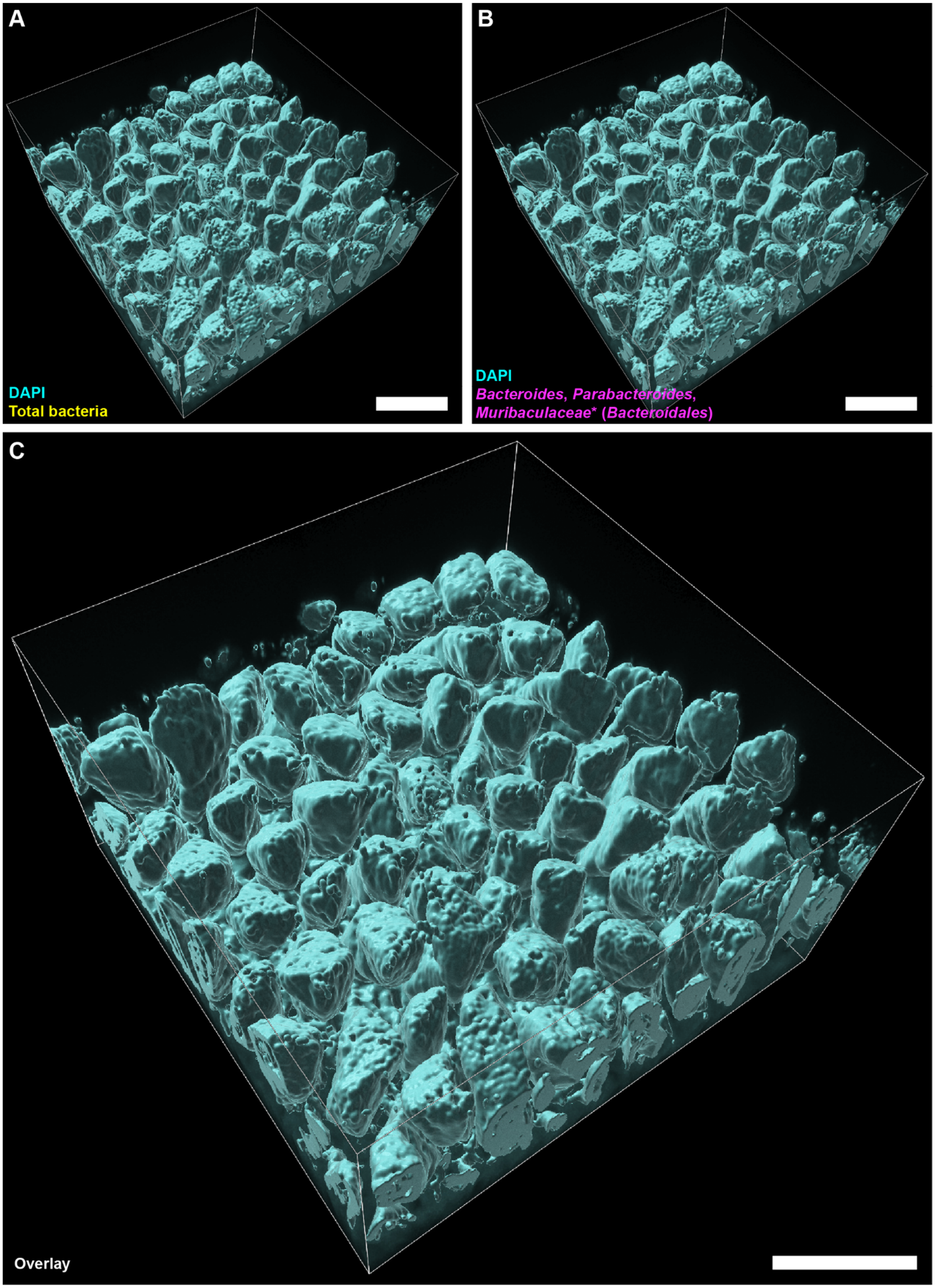


**Figure S29. High-magnification 3D fluorescence imaging of bacteria after 1-h fast in an empty jejunum of MAL+PBS mouse on day 31 of the experiment.** (A-C) 3D rendering of the segmented and filtered surfaces displaying DAPI staining of epithelium (cyan) and (A) HCR v3.0 staining for total bacteria (yellow), (B) HCR v3.0 staining for *Bacteroidales*, and (C) overlay with all groups of bacteria. No significant number of bacteria and no *E. coli* was detected in this field of view. All scale bars 200 µm.

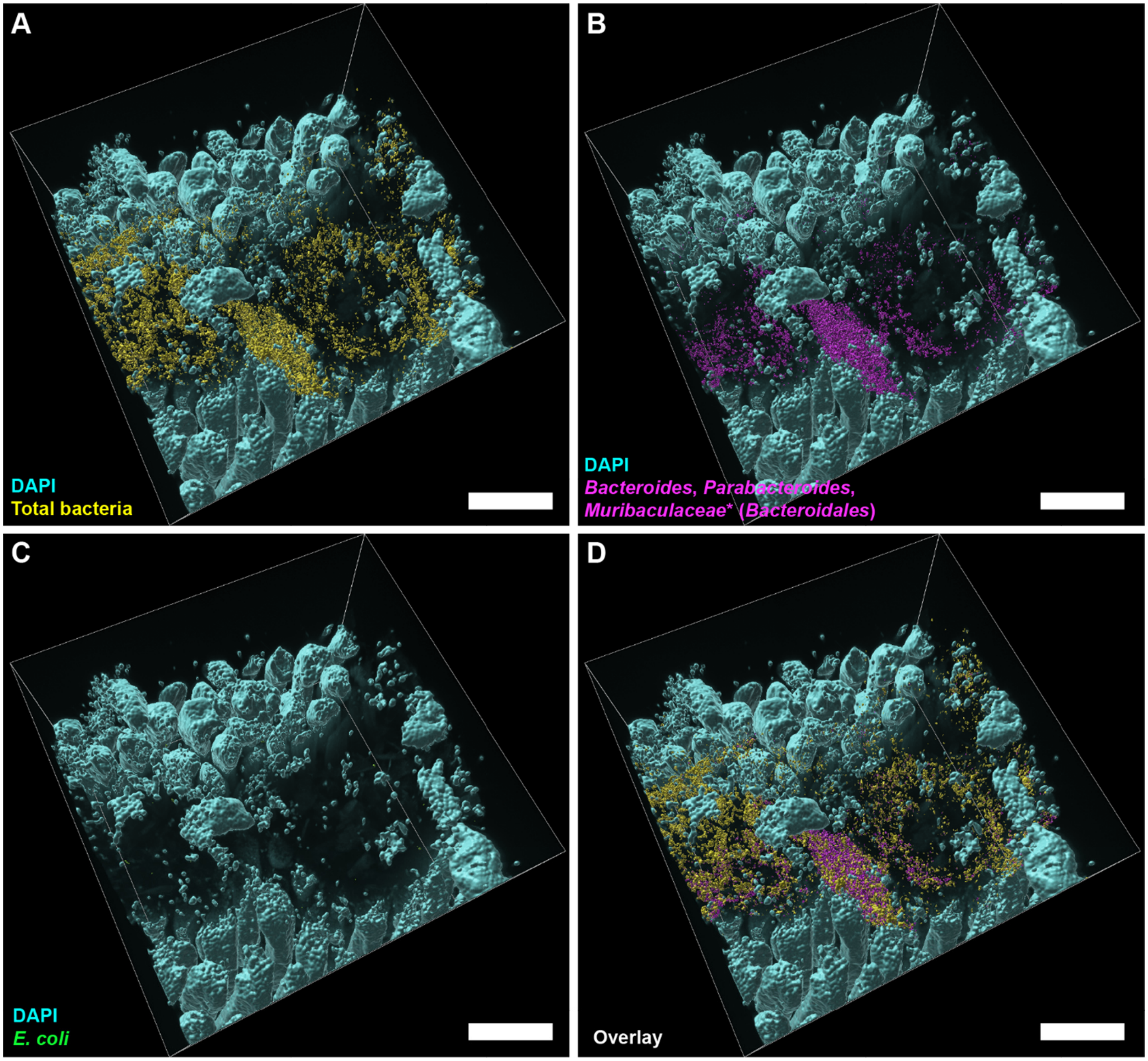


**Figure S30. High-magnification 3D fluorescence imaging of bacteria after 1-h fast in an empty jejunum of MAL+BAC mouse on day 29 of the experiment.** (A-D) 3D rendering of the segmented and filtered surfaces displaying DAPI staining of epithelium (cyan) and (A) HCR v3.0 staining for total bacteria (yellow), (B) HCR v3.0 staining for *Bacteroidales*, (C) HCR v3.0 staining for *E. coli* (green), and (D) overlay all groups of bacteria. A large surface aggregate with abundant bacteria is visible in this field of view. All scale bars 200 µm.

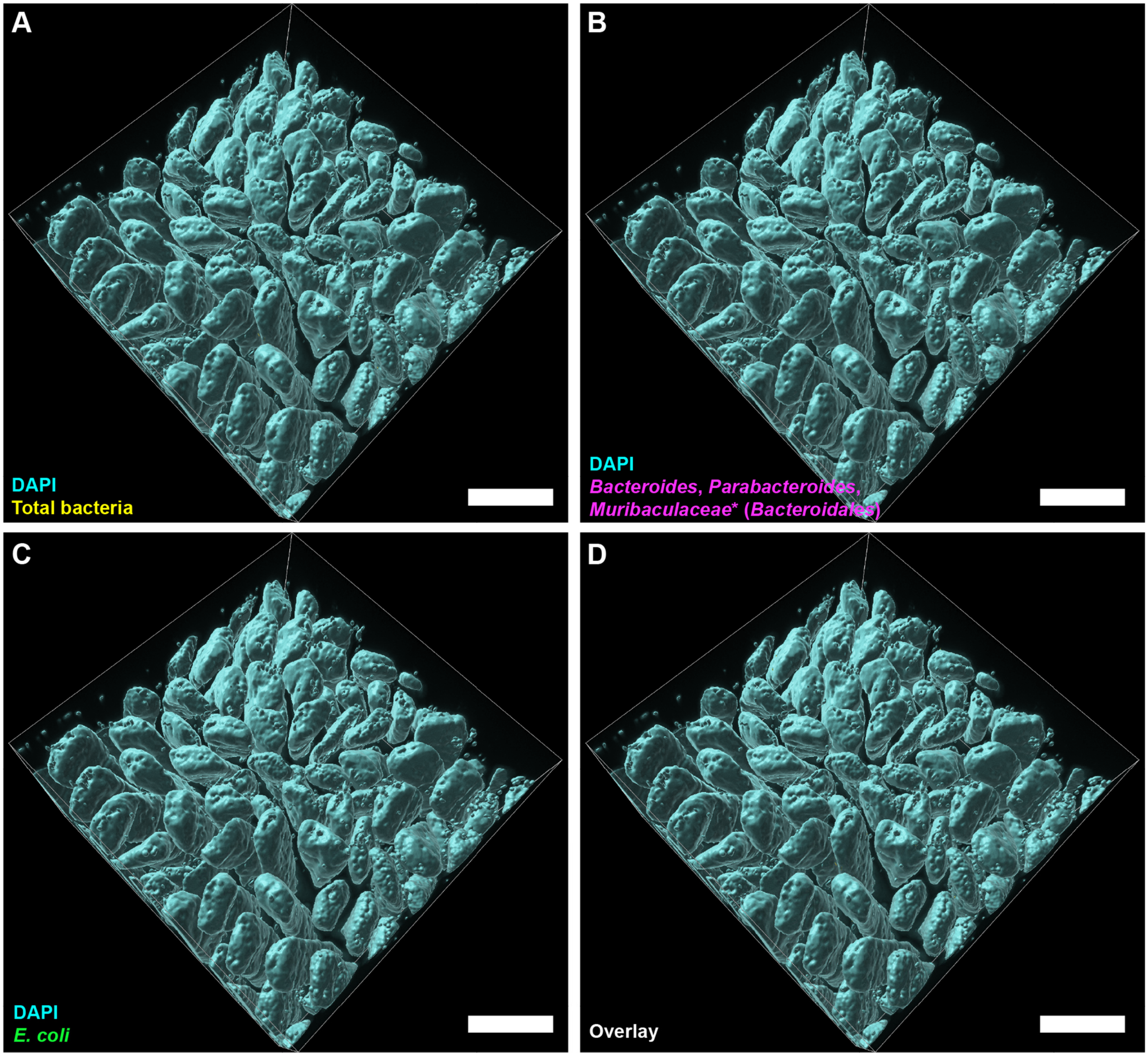


**Figure S31. High-magnification 3D fluorescence imaging of bacteria after 1-h fast in an empty jejunum of MAL+BAC mouse on day 30 of the experiment.** (A-D) 3D rendering of the segmented and filtered surfaces displaying DAPI staining of epithelium (cyan) and (A) HCR v3.0 staining for total bacteria (yellow), (B) HCR v3.0 staining for *Bacteroidales*, (C) HCR v3.0 staining for *E. coli* (green), and (D) overlay with all groups of bacteria. No appreciable number of bacteria was detected in this field of view. All scale bars 200 µm.

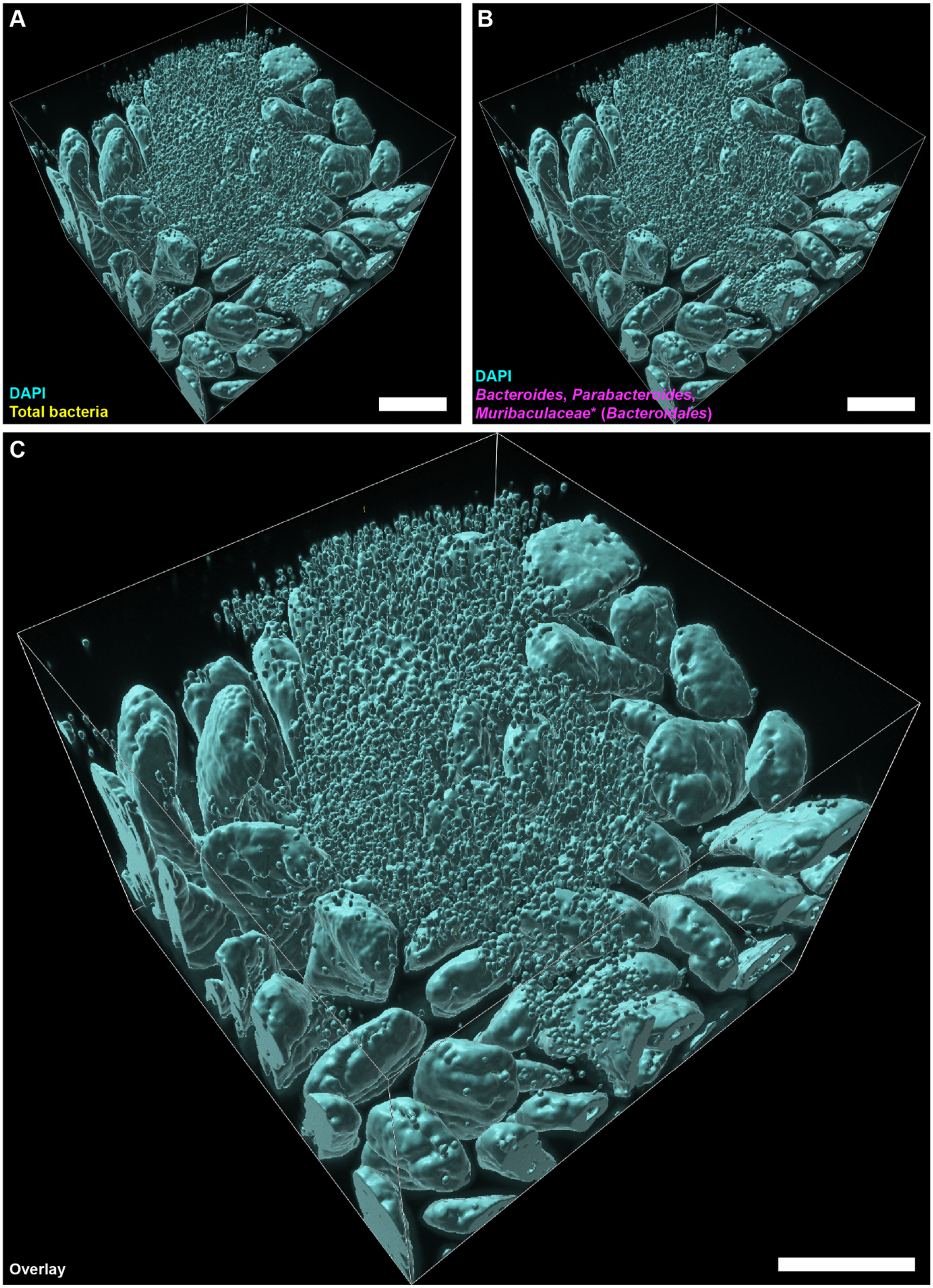


**Figure S32. High-magnification 3D fluorescence imaging of bacteria after 1-h fast in an empty jejunum of MAL+EC mouse on day 28 of the experiment.** (A-C) 3D rendering of the segmented and filtered surfaces displaying DAPI staining of epithelium (cyan) and (A) HCR v3.0 staining for total bacteria (yellow), (B) HCR v3.0 staining for *Bacteroidales*, and (C) overlay with all groups of bacteria. In this field of view, no appreciable number of bacteria and no *E. coli* was detected, but a cluster of abundant free mammalian nuclei is visible. All scale bars 200 µm.

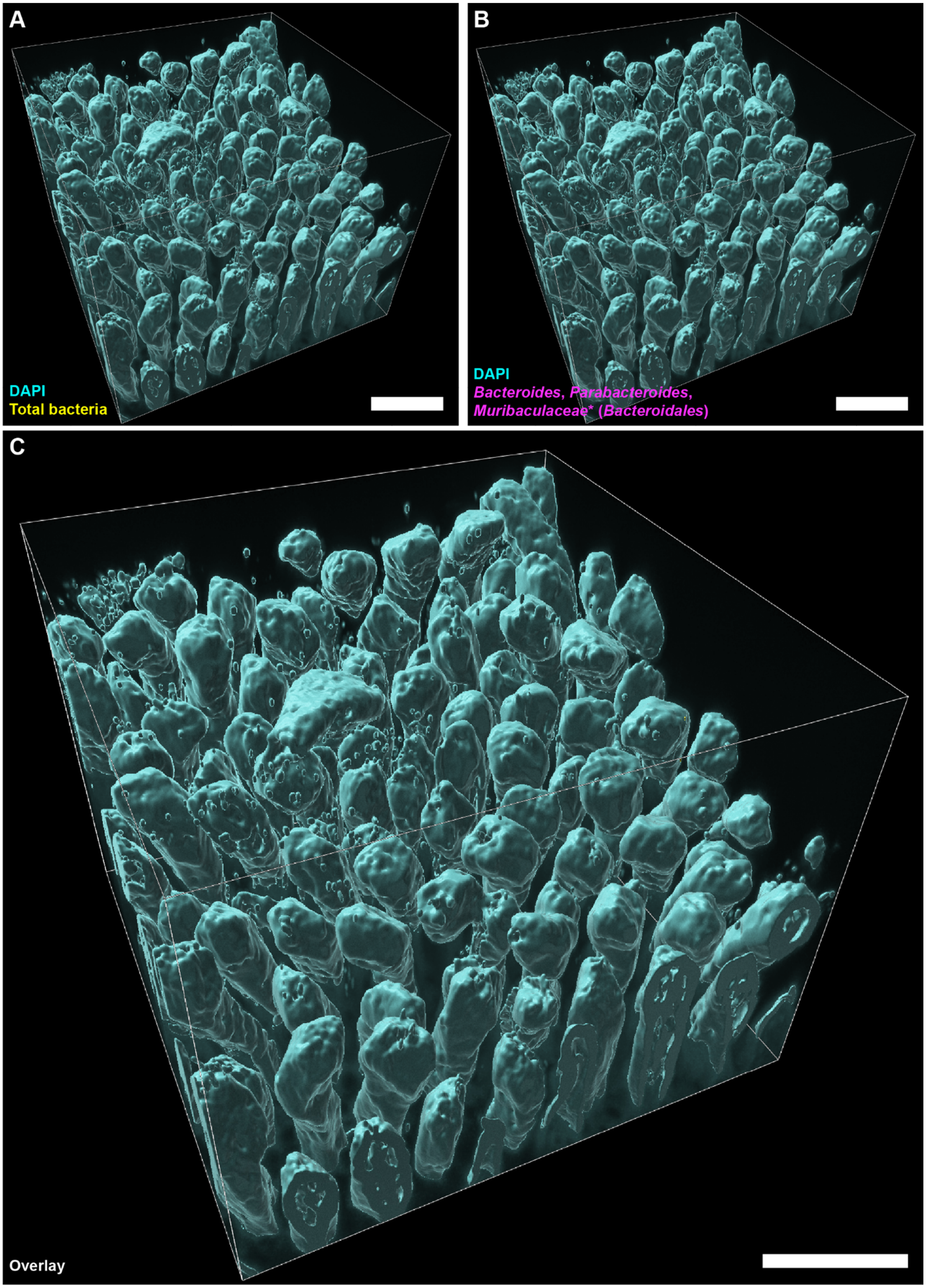


**Figure S33. High-magnification 3D fluorescence imaging of bacteria after 1-h fast in an empty jejunum of MAL+EC mouse on day 31 of the experiment.** (A-C) 3D rendering of the segmented and filtered surfaces displaying DAPI staining of epithelium (cyan) and (A) HCR v3.0 staining for total bacteria (yellow), (B) HCR v3.0 staining for *Bacteroidales*, and (C) overlay with all groups of bacteria. In this field of view, no appreciable number of bacteria and no *E. coli* was detected. All scale bars 200 µm.

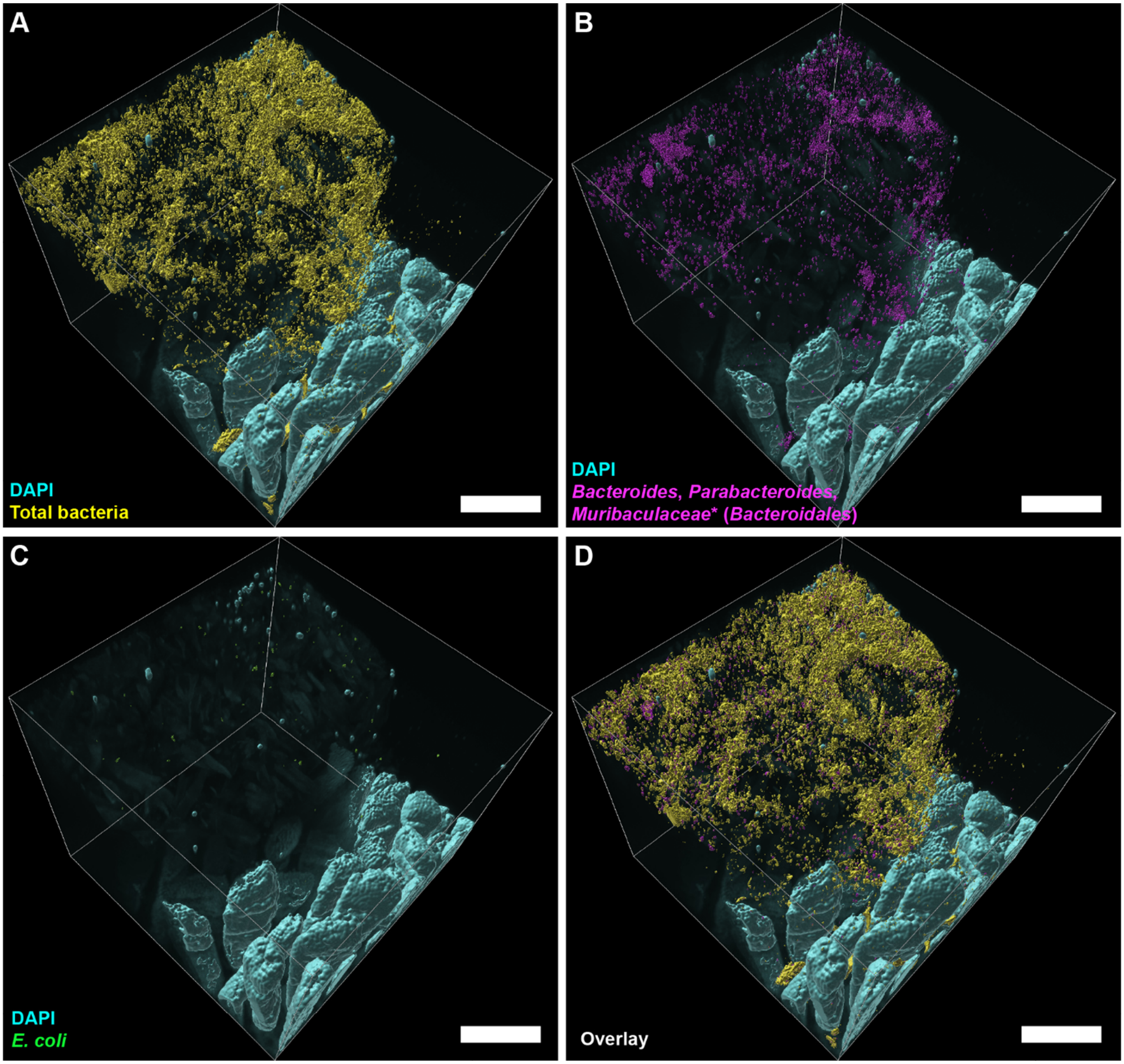


**Figure S34. High-magnification 3D fluorescence imaging of bacteria after 1-h fast in an empty jejunum of MAL+EC&BAC mouse on day 28 of the experiment.** (A-D) 3D rendering of the segmented and filtered surfaces displaying DAPI staining of epithelium (cyan) and (A) HCR v3.0 staining for total bacteria (yellow), (B) HCR v3.0 staining for *Bacteroidales*, (C) HCR v3.0 staining for *E. coli* (green), and (D) overlay all groups of bacteria. A large surface aggregate with abundant bacteria is visible in this field of view. Next to this large surface aggregate, abundant bacteria in between the villi are also visible. All scale bars 200 µm.

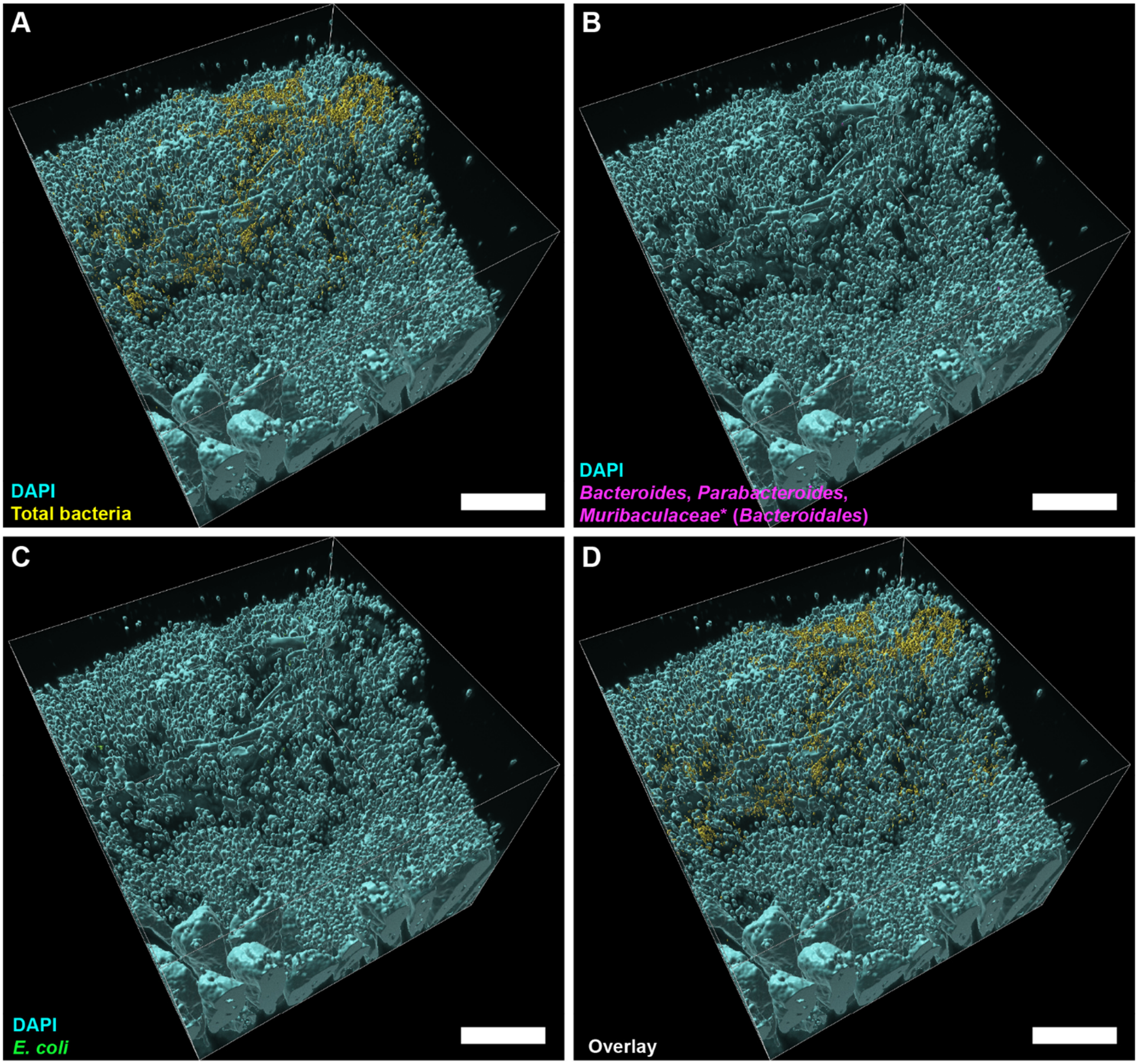


**Figure S35. High-magnification 3D fluorescence imaging of bacteria after 1-h fast in an empty jejunum of MAL+EC&BAC mouse on day 31 of the experiment.** (A-D) 3D rendering of the segmented and filtered surfaces displaying DAPI staining of epithelium (cyan) and (A) HCR v3.0 staining for total bacteria (yellow), (B) HCR v3.0 staining for *Bacteroidales*, (C) HCR v3.0 staining for *E. coli* (green), and (D) overlay all groups of bacteria. A large surface aggregate with abundant bacteria and free mammalian nuclei is visible in this field of view. All scale bars 200 µm.

### Supplementary Video Caption

**Video S1. High-magnification 3D imaging of bacterial colonization of SI mucosa in the empty jejunum of a MAL+EC&BAC mouse.** (0:00:00 – 0:00:02) 3D rendering of the fluorescence intensity data. (0:00:02 – 0:00:09). Fluorescence intensity data across z-sections. (0:00:09 – 0:00:09) 3D rendering of the fluorescence intensity data. (0:00:11 – 00:00:18) Transformation of fluorescence intensity data to the segmented surfaces. 0:00:18-0:00:30) 3D rendering of the segmented surfaces. Cyan: DAPI staining of epithelium. Yellow: HCR v3.0 staining of total bacteria. The same image is displayed in Video S1 as in Figure 4B.

### Supplementary Tables

**Table S1. 16S rRNA gene amplicon sequencing data presented as the number of reads per taxon per sample (TableS1-16SrRNA-GeneAmpliconSequencing.xlsx).** Analysis at different taxonomic levels is provided in separate tabs.

**Table S2. Data for DNA extraction and 16S rRNA gene copy quantification by dPCR (TableS2-16SrRNA-GeneCopyQuantification.xlsx).**

**Table S3. Compositions of malnourished (MAL) and complete control (COM) diets (TableS3-AnimalDiets.xlsx).** Diet compositions are identical as previously described [3] except that food dyes have been omitted.

**Table S4. Surface-gel and tissue-gel monomer mix chemistries used to embed tissues into a hydrogel matrix.**

| | Total % and amount | Acrylamide, 40%[a] | Bis-acrylamide, 2%[b] | PFA, 32%[c] | PBS, 10x | UltraPure Water | VA044 thermal initiator, CAS NO. 27776-21-2[d] |
|---|---|---|---|---|---|---|---|
| Surface-gel-monomer mix chemistries | | | | | | | |
| A4B.08P4 | 100% | 4% | 0.08% | 4.05% | NA | NA | 0.25 w/v% |
| | 30 mL | 3 mL | 1.2 mL | 3.8 mL | 3 mL | 19 mL | 75 mg |
| A4B.08P1 | 100% | 4% | 0.08% | 1.07% | NA | NA | 0.25 w/v% |
| | 30 mL | 3 mL | 1.2 mL | 1 mL | 3 mL | 21.8 mL | 75 mg |
| Tissue-gel-monomer mix chemistries | | | | | | | |
| A4B0P4 | 100% | 4% | 0% | 4.05% | NA | NA | 0.25 w/v% |
| | 30 mL | 3 mL | 0 mL | 3.8 mL | 3 mL | 20.2 mL | 80 mg |
| A1B.01P4 | 100% | 1.25% | 0.0125% | 4% | NA | NA | 0.25 w/v% |
| | 32 mL | 1 mL | 0.2 mL | 4 mL | 3.2 mL | 23.6 mL | 80 mg |

[a]01697; Sigma-Aldrich, St. Louis, MO, USA
b161-0142; Bio-Rad Laboratories, Hercules, CA, USA
c100504-858; VWR, Randor, PA, USA
[d]011-193365; Fujifilm Wako Chemicals, Osaka, Japan

**Table S5. Sequences of HCR v2.0 probes, HCR v3.0 probes, and HCR initiators (TableS5-HCR-InitiatorsAndAmplifiers.xlsx).** Regions of HCR v3.0 probes that overlap with the corresponding HCR v2.0 probes are highlighted in bold.

**Table S6. Imaging, display, segmentation, and filtering metadata (TableS6-ImagingMetadata.xlsx).**

### Supplementary References


1. Barlow JT, Bogatyrev SR, Ismagilov RF: **A quantitative sequencing framework for absolute abundance measurements of mucosal and lumenal microbial communities.** *Nature Communications* 2020, **11:2590**.
2. Choi HMT, Schwarzkopf M, Fornace ME, Acharya A, Artavanis G, Stegmaier J, Cunha A, Pierce NA: **Third-generation in situ hybridization chain reaction: Multiplexed, quantitative, sensitive, versatile, robust.** *Development* 2018, **145:**1-10.
3. Brown EM, Wlodarska M, Willing BP, Vonaesch P, Han J, Reynolds LA, Arrieta M-C, Uhrig M, Scholz R, Partida O, et al: **Diet and specific microbial exposure trigger features of environmental enteropathy in a novel murine model.** *Nature Communications* 2015, **6:**1-16.

### Author Contributions

Roberta Poceviciute (RP)

1. **Idea generation.** Conceived the project with RFI. Conceived the idea of imaging bacteria in the empty intestinal segments of malnourished mice gavaged with bacteria. Conceived the idea of imaging bacteria after 1 h fast. Wrote or contributed to writing grant proposals to fund the project.
2. **Preliminary experiments.** Performed preliminary 16S rRNA gene amplicon sequencing (in collaboration with OMP), 16S rRNA gene copy quantification by dPCR, and imaging (in collaboration with OMP). Supervised preliminary machine-learning based image analysis (performed by HT). Used preliminary data to refine the project.
3. **Method development (HCR v3.0 probe design and validation, hydrogel chemistry optimization).** Starting with the 52 bp region that overlaps with EUB338 binding site (selected by Molecular Technologies), designed the universal degenerate HCR v3.0 probe set. Validated HCR v3.0 probes. Screened different hydrogel chemistries for optimal transport properties.
4. **Data acquisition.** Set up all animal experiments (tail cup experiment was set up in collaboration with SB). Collected and preserved samples for imaging. Collected samples for host gene expression analysis by RTqPCR and 16S rRNA gene analysis by dPCR and sequencing with SB. Extracted RNA for RTqPCR in collaboration with AR. Acquired all imaging data.
5. **Data analysis.** Analyzed RT-qPCR and dPCR data, processed 16S rRNA gene amplicon sequencing, and imaging data. Conceived image segmentation and filtering pipeline.
6. **Figure generation.** Created all figures in the main text and the SI.
7. **Outline writing.** Conceived and wrote outlines.
8. **Manuscript writing.** Wrote and edited the manuscript.

Said R. Bogatyrev

1. **Data acquisition.** Set up tail cup experiment with RP. Collected samples for host gene expression analysis by RTqPCR and 16S rRNA gene analysis by dPCR and sequencing with RP.

Anna E. Romano

1. **Data acquisition.** Extracted RNA with RP and quantified host gene expression by RTqPCR (Fig. 1, C and D). Extracted DNA and quantified 16S rRNA gene copy load (Fig. 2, F and G). Prepared 16S rRNA gene amplicon library for sequencing (Fig. 2A).

Amanda H. Dilmore

1. **Data acquisition.** Extracted DNA for 16S rRNA gene amplicon sequencing and 16S rRNA gene copy quantification (Fig. 2, A-E). Quantified 16S rRNA gene copies by dPCR (Fig. 2, B-E, and Fig. S1).

Octavio Mondragón-Palomino

1. **Preliminary experiments.** Performed preliminary 16S rRNA gene amplicon sequencing (in collaboration with RP) and 3D imaging of tissue samples from the small intestines of mice in the environmental enteropathy model (in collaboration with RP). Contributed practical and technical guidance on spectral confocal imaging of clarified tissues.

Heli Takko

1. **Preliminary experiments.** Performed preliminary machine-learning based image analysis (supervised by RP).

Rustem F. Ismagilov

1. **Idea generation.** Conceived the project with RP.
2. **Data acquisition.** Provided feedback on experimental design.
3. **Funding.** Secured funding for the project.
4. **Data analysis.** Provided feedback on data analysis.
5. **Outline writing.** Provided feedback on outlines written by RP.
6. **Manuscript writing.** Edited the manuscript written by RP.
